# Dimension Reduction using Local Principal Components for Regression-based Multi-SNP Analysis in 1000 Genomes and the Canadian Longitudinal Study on Aging (CLSA)

**DOI:** 10.1101/2024.05.13.593724

**Authors:** Fatemeh Yavartanoo, Myriam Brossard, Shelley B. Bull, Andrew D. Paterson, Yun Joo Yoo

## Abstract

For genetic association analysis based on multiple SNP regression of genotypes obtained by dense DNA sequencing or array data imputation, multi-collinearity can be a severe issue causing failure to fit the regression model. In this study, we proposed a method of Dimension Reduction using Local Principal Components (DRLPC) which aims to resolve multi-collinearity by removing SNPs under the assumption that the remaining SNPs can capture the effect of a removed SNP due to high linear dependency. This approach to dimension reduction is expected to improve the power of regression-based statistical tests. We apply DRLPC to chromosome 22 SNPs of two data sets, the 1000 Genomes Project (phase 3) and Canadian Longitudinal Study on Aging (CLSA), and calculated Variance Inflation Factors (VIF) in various SNP-sets before and after implementing DRLPC as a metric of collinearity. Notably, DRLPC addresses multi-collinearity by excluding variables with a VIF exceeding a predetermined threshold (VIF=20), thereby improving applicability for subsequent regression analyses. The number of variables in a final set for regression analysis is reduced to around 20% on average for larger-sized genes, whereas for smaller ones, the proportion is around 48%; suggesting that DRLPC is more effective for larger genes. We also compare the power of several multi-SNP statistics constructed for gene-specific analysis to evaluate power gains achieved by DRLPC. In simulation studies based on 100 genes with ≤500 SNPs per gene, DRLPC effectively increased the power of the multiple regression Wald test from 60% to around 80%.

## 1 INTRODUCTION

Genetic association studies investigate associations between single nucleotide polymorphisms (SNPs) and a trait of interest (Yu, 2012; Yoo et al., 2017; Xue et al., 2020). High-density SNP genotypes generated from genome-wide genotyping arrays and imputed to a reference panel, or Next-Generation Sequencing (NGS) technologies are analyzed to detect SNP-trait association signals (Gauderman et al., 2007; Kim et al., 2010; Slavin et al., 2011; Wu et al., 2011). A single-SNP analysis considers the association between a trait and one SNP at a time (Syvänen, et al., 2001; Spencer et al., 2009; Kim et al., 2010). On the other hand, multi-SNP analysis investigates the association between a trait and multiple SNPs simultaneously. In the multi-SNP approach, a set of SNPs is considered together in region-level analysis, e.g. SNPs within a region defined by gene boundaries, that obtain a global statistic to test for the combined effect of the SNP set. Global testing can yield robust, powerful, and informative results (Asimit et al., 2009; Chapman & Whittaker, 2008). In particular, SNP genotypes within a gene can be analyzed using a multi-SNP regression model, and joint effects of a SNP set tested by a large sample Wald statistic with multiple *df* (Clayton et al., 2004; Gauderman et al., 2007; Wang et al., 2012; Yoo et al., 2013).

In multi-SNP joint regression analysis, issues with multi-collinearity can occur when there are a large number of predictors or when the ratio of number of observations relative to the number of predictors is not large. In regression, multi-collinearity means linear or near-linear dependency among two or more predictors, which corresponds to a lack of orthogonality among them (Alin, 2010). The linkage disequilibrium (LD) structure of high-density SNP genotype data often shows clusters of highly correlated SNP, which may or may not be consecutively located (Kim et al., 2018; Kim et al., 2019). High-density SNP genotype predictors in proximity often yield multi-collinearity due to LD (Wang et al., 2012). Multi-collinearity can cause a singular covariance matrix due to independent variables. This singularity arises because the matrix’s determinant approaches zero, making it mathematically unstable for inversion (Farrar et al., 1967). Multi-collinearity can lead to misleading or unexpected signs of the regression coefficient, as their signs may deviate from the expected relationship between predictors and the response variable. The most severe effect of multi-collinearity is the inflation of standard errors associated with regression coefficients, signifying large sampling variability (Alin, 2010). Ways to detect collinearity among predictors, include pairwise correlations, eigenvalues, and the Variance Inflation Factor (VIF). The correlation matrix and eigenvalues can provide indications of the existence of multi-collinearity; however, they cannot accurately measure the extent of multi-collinearity. The VIF, which measures how much the variance of the estimated regression coefficient is inflated due to collinearity among the variables, quantifies multi-collinearity and effects on computation and can be interpreted quickly and clearly (Gwelo et al., 2019).

Applying Principal Component Analysis (PCA, Edgeworth, 1884) for both SNP genotype and haplotype-based approaches within multiple regression analysis is an efficacious dimension reduction^1^ technique (Wang et al., 2008; Chapman et al., 2003; and Clayton et al., 2004). Moreover, this approach offers a valuable statistical technique for detecting, quantifying, and adjusting for multi-collinearity in a dataset (Lafi et al., 1992). Several methods have been developed to reduce the dimension of complex data; however, PCA is a conventional dimension reduction approach for multivariate data that depends on an orthogonal linear combination of variables, called principal components (PCs) (Li, 2010; Abdi & Williams, 2010; Park et al., 2020). Wang & Abbott (2008) have applied principal components regression (PCReg) to test for the association of a set of SNPs with the phenotype, assuming that a small number of variables can model a sufficient amount of variation in the joint distribution of all SNPs. As a result of this, a few PCs, which are computed from the sample covariance matrix of the SNP genotype, are used as regressors in multiple regression.

In the context of high-dimensional genetic association analysis, when the number of genetic predictors exceeds the sample size, the model cannot be fit at all using standard least squares methods. It leads to overfitting, high variance stability, singular and collinearity, reduced statistical power, misleading interpretation, and inflated type II errors. Dimension reduction streamlines the model by focusing on a reduced set of features. This strategy counters the risk of overfitting, enhancing the power of statistical tests. Replacing SNP variables with principal components can improve the power of regression-based statistical tests by addressing the challenges associated with high dimensionality. Gauderman et al., (2007) highlighted the advantages of the PC approach over joint-SNP and haplotype tests. Their findings emphasized that PCs capture SNP variation within the genetic locus. Substantive dimension reduction was achieved by adopting an 80% explained variance threshold for the disease model, which gained statistical power. However, the PCs constructed from all SNP variables in a region are hard to interpret as biological entities and are not helpful for localization and fine mapping.

While PCA can project high-dimensional data onto a lower-dimensional space that captures most of the original data’s variance using the variables’ correlation structure, it cannot represent local information for data with complicated distributions (Wold et al., 1987; Yu, 2012). If the variables have non-linear dependencies, PCA will require a more significant dimensional representation than would be found by a non-linear technique. Kambhatla & Leen (1997) introduced an extension to PCA using a local linear approach called Local PCA (LPCA). In contrast to global PCA, which projects an entire set of variables onto a low-dimensional space, LPCA focuses on subsets or “local” regions of the data to capture more nuanced or complex relationships that may be missed by global PCA. Similar to PCA, LPCA can relieve multi-collinearity and reduce data dimensionality. The LPCA algorithm for *n*-dimensional input data can be stated as follows:

*Step 1*. First, partition the input data into Q disjoint regions {*R*^(1)^, *R*^(2)^, . . ., *R*^(*Q*)^}.

*Step* 2. Compute the local covariance matrix ∑^(*i*)^ = *E*[(*x* − *Ex)(x* − *Ex*)^*T*^|*x* ∈ *R*^(*i*)^]; *i* = 1, …, *Q*, for the variables in region and their eigenvectors 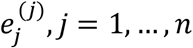, *j* = 1, …, *n*. Without loss of generality, the eigenvectors can be relabeled so that the corresponding eigenvalues are in descending order 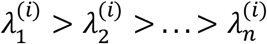.

*Step* 3. Select a target dimension *m* and retain the leading *m* eigenvector directions for reducing the data dimension.

In the present article, we propose a new approach to gene-level association analysis based on LPCA that aims to improve the power of global multi-SNP test statistics while preserving the interpretability of the localized effects. In regression analysis, Dimension Reduction using Local Principal Components (DRLPC) proceeds by first selecting clusters of SNPs in high correlation and replacing each cluster with a local principal component constructed from the SNPs in the cluster. DRLPC also aims to resolve multi-collinearity among the updated variables by removing variables with VIF values greater than a predefined threshold iteratively until the highest value of VIF falls under the threshold. DRLPC, using an LPCA technique and removing variables with a high VIF with the underlying assumption that the remaining variables possess the capacity to capture the impact of the removed variables due to their high linear dependency, can simultaneously reduce the dimensionality of the data and resolve multi-collinearity.

To examine the behavior of DRLPC in achieving dimension reduction, we applied the algorithm to SNP genotypes from two data sets. The first is the 1000 Genomes Project data (phase 3), chromosome 22 of three major super-populations: European, East Asian, and African. The second dataset is the European ancestry subset of the Canadian Longitudinal Study on Aging (CLSA), also for chromosome 22 SNPs. The dimension of each dataset was reduced separately, considering several choices of threshold values for clustering and principal components selection. We also designed simulation studies to generate quantitative trait values from the 1000 Genomes Projects genotype under gene-level regression models for genetic association. Analyses of the original SNP genotype variables and the DRLPC processed variables were then conducted to compare type I error and power of several multi-variable statistics including generalized linear regression Wald (Wald, 1943), multiple linear combination (MLC) (Yoo et al., 2017, generalized Wald with global principal components (Gauderman et al., 2007), and sequence kernel association (SKAT) and SKAT-O ((Ionita-Laza et al., 2013; Lee et al., 2012).

The rest of the paper is organized into the following sections: Section 2 includes a description of the DRLPC method. Section 3 discusses the results obtained from the proposed method for actual data application using two different SNP sets for two datasets. Section 4 reports simulation study results for the power and type I error of multi-marker statistics obtained using DRLPC processed datasets. The discussion and conclusion are given in Section 5 and Section 6, respectively.

## 2 MATERIALS AND METHODS

### 2.1 Dimension reduction using local principal components

Suppose that the genotypes of *m* SNPs are coded as 0, 1, or 2 based on an additive genetic model and denoted by *X* = (*X*_1_, *X*_2_, …, *X*_*m*_). The multi-SNP joint regression model of *m* SNPs consider *E*[*Y*|*X*] as the expected value of quantitative trait *Y* conditional on the given SNP genotypes and is formulated as follows:

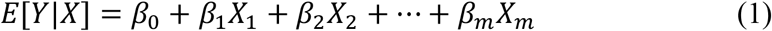

Global test statistics based on the above regression analysis can be constructed from the beta estimates 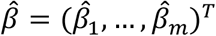 and the associated covariance matrix ∑_*B*._ One of the objectives of the DRLPC technique is to reduce the dimension of data before conducting regression analysis. To achieve this, the DRLPC employs two strategies for dimension reduction: local PCs and filtering on VIF. These strategies are employed to reduce the number of regression variables and remove multi-collinearity.

Figure 1 illustrates the steps in the DRLPC processing with an example region “*SLC35E4*” in chromosome 22 (bp position of 31,031,643 ∼ 31,064,736), which includes 57 SNPs in 1000 Genomes Projects (phase 3), EUR super-population.

**FIGURE 1.**
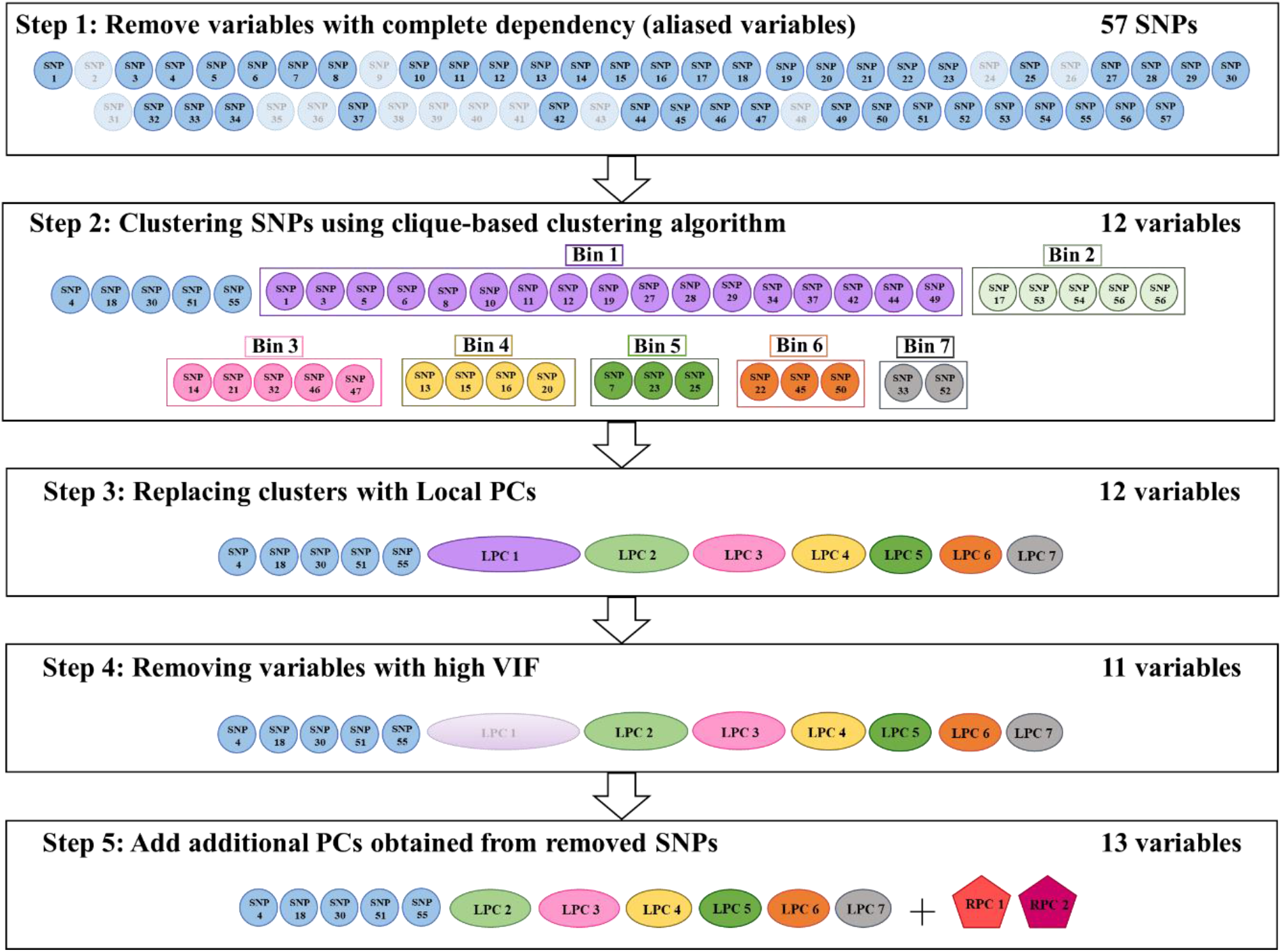
Illustration of DRLPC algorithm applied for “*SLC35E4”* region (chromosome 22) data from 1000 Genomes Project, EUR population data. Blurred variables at Step 1 and Step 4 denote those removed in that step.

The DRLPC algorithm requires three thresholds: a threshold (CLQcut) for pairwise *r*^2^ to construct clusters of highly correlated SNPs by the clique-based graph partitioning method (Yoo et al., 2015); a threshold (VIFcut) for variance inflation factor (VIF) values to remove variables with linear dependencies and reduce multi-collinearity; a threshold (PCcut) to select some PC variables that capture the variability of all the removed variables as the candidates for addition of variables at the last step of the algorithm. DRLPC also designates a value (Klim) as the partition limit for the alias^2^ removal step to apply this step separately for each subset of partitioned SNPs when the number of SNPs is greater than the sample size. The details of the DRLPC algorithm are explained as follows:

*Step 1*. ***Removal of linearly dependent SNPs:*** Suppose that *m* SNPs in a gene are indexed with *V*_1_ = {1,2, …, *m*}. Let *V* ≔ *V*_1_. First, SNPs with complete linear dependency (aliased SNPs) are detected, and one SNP per group of aliased SNPs forming each linear dependency relationship is removed. The set of SNPs removed by this process is denoted by *W*_1_. If the number of SNPs is greater than or close to the sample size, we partition SNPs into subsets, including fewer SNPs than the sample size. A partition limit (Klim) is set to partition the SNPs into sets with a size less than Klim in which the procedure of breaking the linear dependency is applied to each partition separately. Since the linear dependency may occur between partitioned parts, we apply this process repeatedly for the combined sets of the remaining SNPs until no completely dependent SNPs remain. In this way, the set of current SNPs *V* is updated by *V*_2_ = *V*\*W*_1_.

*Step 2*. ***Clustering of highly correlated SNPs:*** Using the clique-based clustering algorithm CLQ-D in Kim et al., (2018), find the groups of SNPs in *V*_2_ that have pairwise *r*^2^ greater than the CLQcut. Some SNPs do not form groups and remain as singletons. Denote the groups (bins) with multiple SNPs found in this step by *G* = {*G*_1_, ⋯, *G*_*j*_}. Also, denote the set of singleton SNPs by *S*.

*Step 3*. ***Replacing each group with a local PC:*** Replace the SNPs in each *G*_*j*_ by the first principal component obtained applying PCA only for the SNPs within the group, known as a local PC (LPC), which is denoted by *LPC*_*j*_. Now let *L* = {*LPC*_1_, ⋯, *LPC*_*j*_}. Then, update the current set of dimension-reduced variables *V*_2_ with *V*_3_ = *S* ∪ *L* = *S* ∪ {*LPC*_1_, ⋯, *LPC*_*j*_}. There is a possibility of no LPC replacing the SNPs when no cluster of multiple SNPs is found in Step 2.

*Step 4*. ***Removal of variables to reduce multi-collinearity:*** Suppose the variables in *V*_3_ are indexed such that *V*_3_ = {*v*_1_, *v*_2_, …, *v*_*K*_}, regardless they are the original SNPs or LPCs. First, calculate 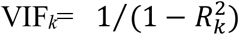 for each variable in *V*_3_ where 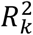 is the coefficient of determination obtained from the regression of *v*_*k*_ on the other variables in *V*_3_. Next, remove the variable with the highest VIF value from *V*_3_. Repeat this procedure iteratively until the highest value of VIF becomes under the threshold for VIF (VIFcut). The set of removed variables in this step is named *W*_2_. Update the current set of dimension-reduced variables *V* with *V*_4_ = *V*_3_\*W*_2_.

*Step 5*. ***Selective addition of PCs representing the removed SNPs:*** In the last step, another PCA step is applied to the set of SNPs removed in Steps 1 and 4. In Step 4, if an LPC is selected to be removed due to high VIF, the SNPs in the cluster corresponding to the LPC are all considered to be removed. Take the minimum number of PCs with their proportion of cumulative explained variance greater than the threshold value PCcut as the candidate for added variables. Then, examine if the variability of these PCs can be captured by the current set *V* = *V*_4_ by regression each of these PC on the variables in *V* and obtain the *R*^2^. If the *R*^2^ is less than 1/(1 − *VIFcut*) add the corresponding PC into the current set *V*. The set of PCs selected to be added in this step is denoted as *R*, and they are called RPCs. Update the current set *V* with *V*_5_ = *V*_4_ ∪ *R*.

*Final set of dimension-reduced variables*. In this way, the dimension of high-density SNP data is reduced iteratively, and the final variables after the dimension reduction process are the union of singleton SNPs and LPCs that are not removed by step 4 and RPCs added by step 5. The intersection of these three sets is empty, and each of the three sets can also be an empty set.

## 3 APPLICATIONS of DRLPC to SNP GENOTYPES

### 3.1 Datasets

To evaluate the performance of the DRLPC algorithm, we applied it in two datasets:

▹ the 1000 Genomes Project (phase 3) chromosome 22 data (1000 Genomes Project Consortium., (2015)) from three super-populations: European (EUR), East-Asian (EAS), and African (AFR), from https://ftp.1000genomes.ebi.ac.uk/vol1/ftp/release/20130502/.
▹ the HRC-imputed SNP genotype datasets of the Canadian Longitudinal Study on Aging (CLSA) (Raina et al., 2009). QC was applied as described in Forgetta et al., (2022) and summarized in supplemental methods.

We limited analysis to the data of European ancestry since other groups do not have large enough sample sizes. The datasets were preprocessed similarly. First, SNPs with missing values and multi-allelic SNPs were removed. Also, SNPs with minor allele frequency (MAF) less than 0.05 were excluded. In the CLSA, we required an INFO score ≥ 0.8 to select well-imputed SNPs. Table 1 reports the number of individuals in each population and the total number of remaining SNPs post-filtering for each of the 1000 Genomes and CLSA populations.

**TABLE 1.** Sample size, number of SNPs on chromosome 22, number of SNP sets of the 100 SNPs, number of SNPs in genes, and number of genes for each population dataset studied.

| Population | Sample size | Number of SNPs (total) | Number of SNPsets (100 SNPs per set) | Number of SNPs (total in genes) | Number of Genes |
| --- | --- | --- | --- | --- | --- |
| <sup>†</sup>EUR (1KG) | 503 | 85,718 | 857 | 50,705 | 698 |
| <sup>‡</sup>EAS (1KG) | 504 | 78,461 | 784 | 46,476 | 693 |
| <sup>§</sup>AFR(1KG) | 661 | 118,466 | 1,184 | 70,471 | 734 |
| European (CLSA) | 17,779 | 71,695 | 717 | 71,695 | 611 |

<sup>†</sup>EUR: European population; <sup>‡</sup>EAS; East Asian population; <sup>§</sup>AFR: African population

### 3.2 Methods

To examine the population-specific dimension reduction results, we created two types of SNP sets from each population dataset: 1) a set of 100 consecutive SNPs and 2) a set of SNPs in gene regions. For the 100 SNP sets, we partitioned all SNPs in chromosome 22 into sets of 100 consecutive SNPs. For gene-based datasets, we considered only the SNPs within gene regions, based on the gene information obtained from the Ensemble BioMart database for NCBI hg19 Build 37 (GRCh37.p13) (http://grch37.ensembl.org/biomart). Table 1 reports the number of SNPsets of 100 SNPs, the total number of SNPs in genes, and the number of genes with more than one SNP for each population. Table 2 also reports summary statistics of the number of SNPs in a gene for each population.

**TABLE 2.** Summary statistics of numbers of SNPs per gene on chromosome 22 for each population.

| Population | Number of genes | Summary of the number of SNPs in a gene |  |  |  |  |
| --- | --- | --- | --- | --- | --- | --- |
|  |  | Average | SD | Median | Min | Max |
| EUR | 698 | 72.6 | 156.7 | 26 | 2 | 1875 |
| EAS | 693 | 67.1 | 142.1 | 24 | 2 | 1846 |
| AFR | 734 | 96.0 | 209.5 | 36 | 2 | 2764 |
| European (CLSA) | 611 | 71.8 | 151.0 | 27 | 2 | 1759 |

To assess the performance of the proposed method, we applied DRLPC to each SNP set with several combinations of threshold values required for the algorithm. For CLQcut, which is the threshold value for the clique-based clustering algorithm CLQD (Yoo et al., 2015) used to find SNP clusters, we designated four values: 0.5, 0.8, 0.9, and 0.9. For VIFcut, which is the threshold value for variable removal based on the variance inflation factor (VIF) to reduce multi-collinearity, we assigned a fixed value of 20. For PCcut, which is the threshold value for selecting additional PCs representing the removed SNPs at the final step of the algorithm, we chose 0.8 and 0.9. Eight different combinations of threshold values were applied to obtain dimension reduction results by DRLPC.

### 3.3 Results

#### 3.3.1 Dimension reduction results of 1000 Genomes Project dataset

When the DRLPC algorithm is applied to the SNP sets in gene regions of chromosome 22, the number of variables in genotype data was reduced to 8∼16% in EUR and EAS and 18∼31% in AFR, depending on the threshold values used (Figure 2, and Supplementary Table S1). When the DRLPC algorithm is applied to each of the 100 consecutive SNP sets, the dimension of genotype data was reduced to 13-25% in EUR and EAS and 22-41% in AFR, depending on the threshold values used (Figure S1, and Supplementary Table S2). The dimension reduction in AFR by DRLPC was less than in EUR and EAS, which can be explained by weaker LD in the AFR population, fewer sites being in LD, and divergent patterns of LD between AFR and non-AFR (Tishkoff et al., 2002). In both types of SNP sets, it was observed that lower CLQ threshold values led to greater dimension reduction. The percentages of the final number of variables after the DRLPC process compared to the number of SNPs in the original dataset were very slightly lower with PCcut value of 0.8, compared to a PCcut value of 0.9.

**FIGURE 2.**
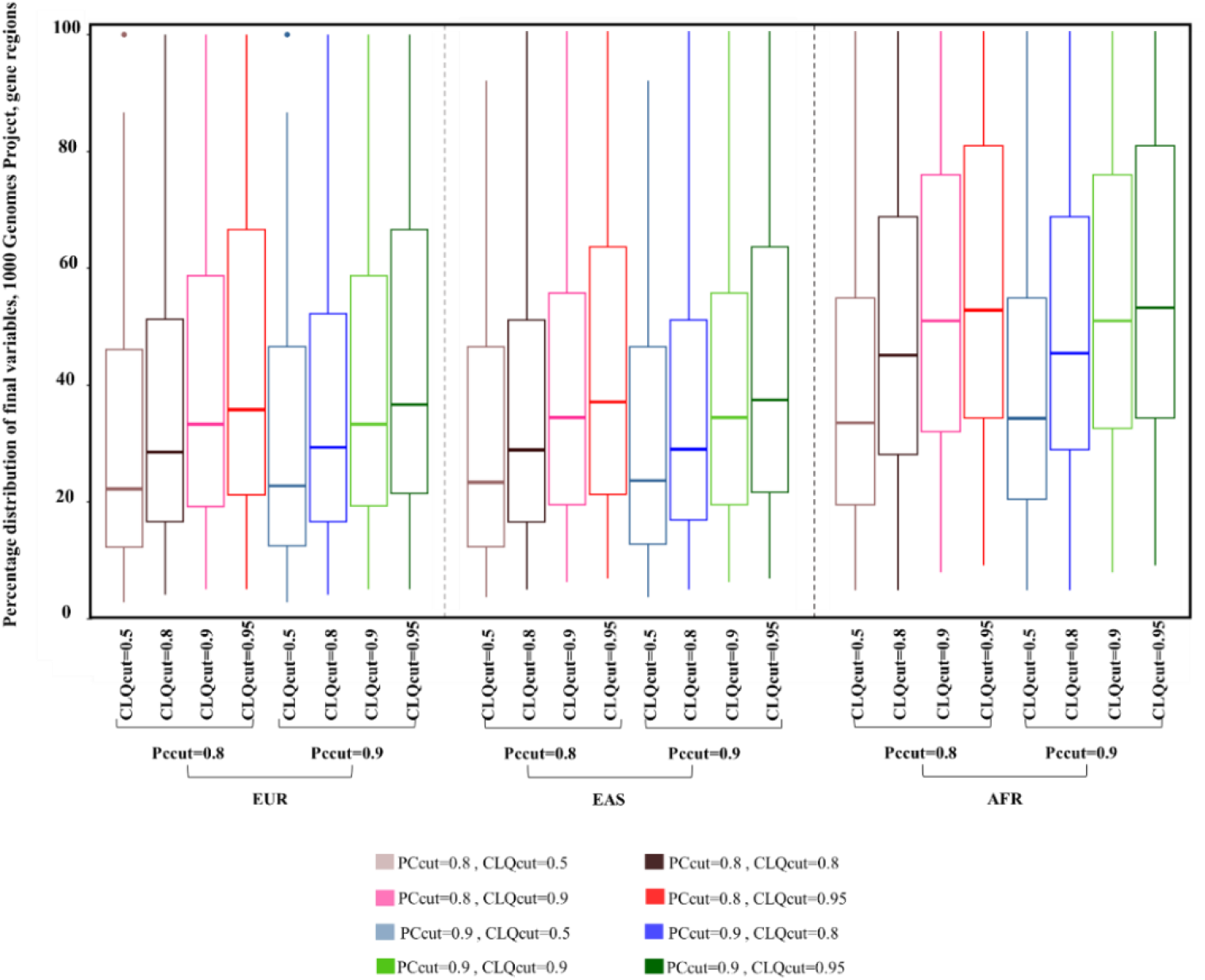
The percentage distribution of final variables after the DRLPC process compared to the number of SNPs in the original gene-based datasets of 1000 Genomes Project, chromosome 22, with four threshold values for CLQcut (0.5, 0.8, 0.9, 0.95), two threshold values (0.8, 0.9) for PCcut, and for three super-populations (EUR, EAS, AFR).

Figure 3 presents the relationship between gene size and the reduction rate across all 1000 Genomes super-populations and CLSA data European ancestry (considering a PCcut value of 0.8, a comparable plot for a PCcut value of 0.9 is provided in Supplementary Figure S2). The reduction rate is defined as one minus the proportion of final variables after the DRLPC process relative to the number of variables in the original data. An upward trend is observed, indicating that the reduction rate also tends to increase the number of SNPs in a gene increase. Similar trends are observed across all super-populations and different choices of threshold values. We also compared the reduction rates resulting from the DRLPC process for each super-population after dividing genes into two groups considering the number of SNPs: 1) genes with below-average numbers of SNPs and 2) genes with above-average numbers of SNPs. The average number of SNPs per gene for EUR, EAS, and AFR populations was 72.6, 67.1, and 96.0, respectively. As presented in Table 3, the differences in average reduction rates are about 28∼35% between the genes with a larger number of SNPs (above average group) and those below the average group. On average, the dimension reduction rates for genes with more SNPs were around 83% for EUR and EAS populations and 75% for the AFR population. In comparison, the corresponding rates for genes with fewer SNPs were around 53% for both the EUR and EAS populations and 42% for the AFR population. These results align with African populations exhibiting reduced levels of linkage disequilibrium (LD) compared to non-African populations (Campbell et al., 2008).

**TABLE 3.**
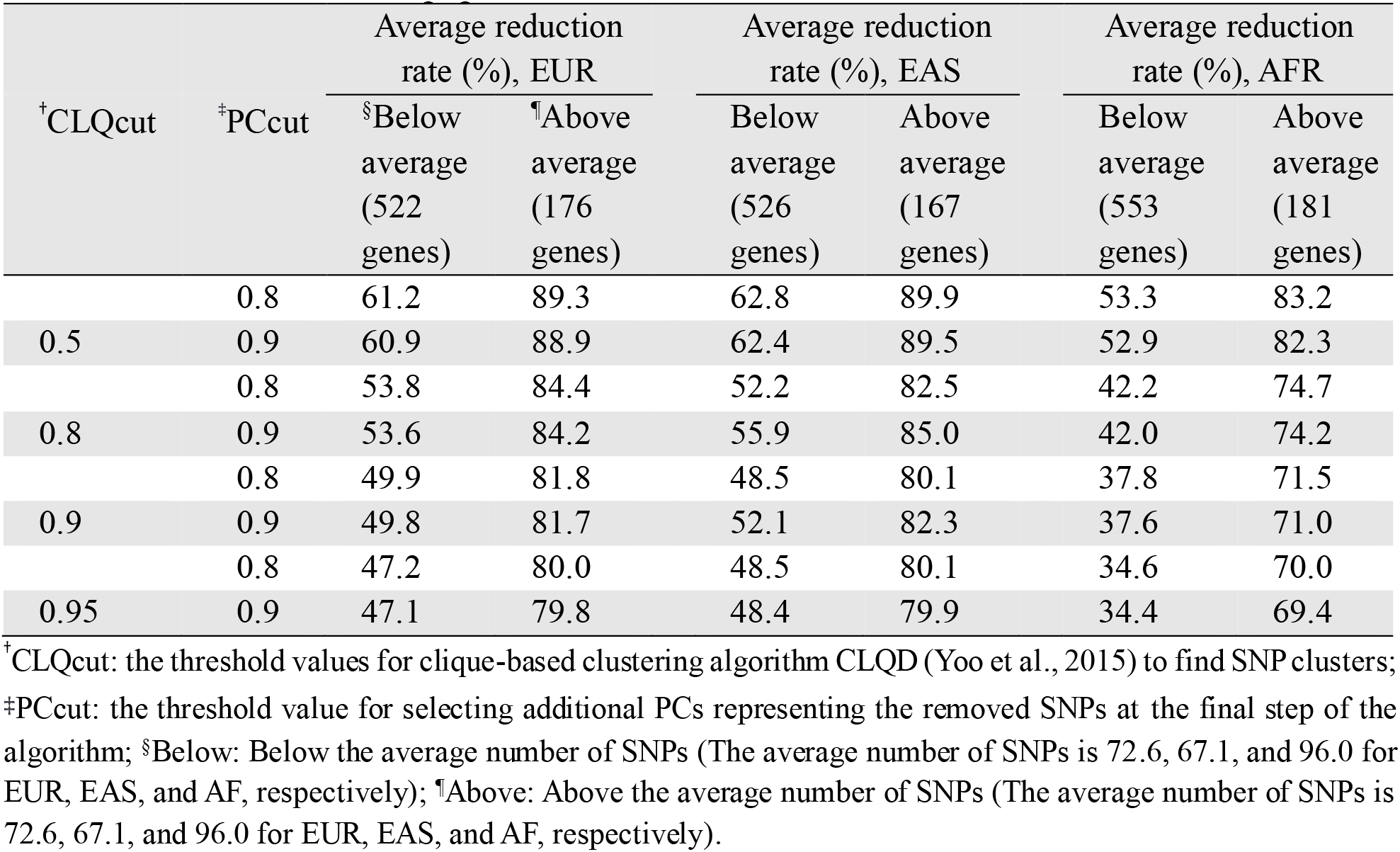
Average reduction rate after (percentage) in the number of variables after the DRLPC process compared to the original number of SNPs in the gene regions of chromosome 22 for EUR, EAS, and AFR populations.

**FIGURE 3.**
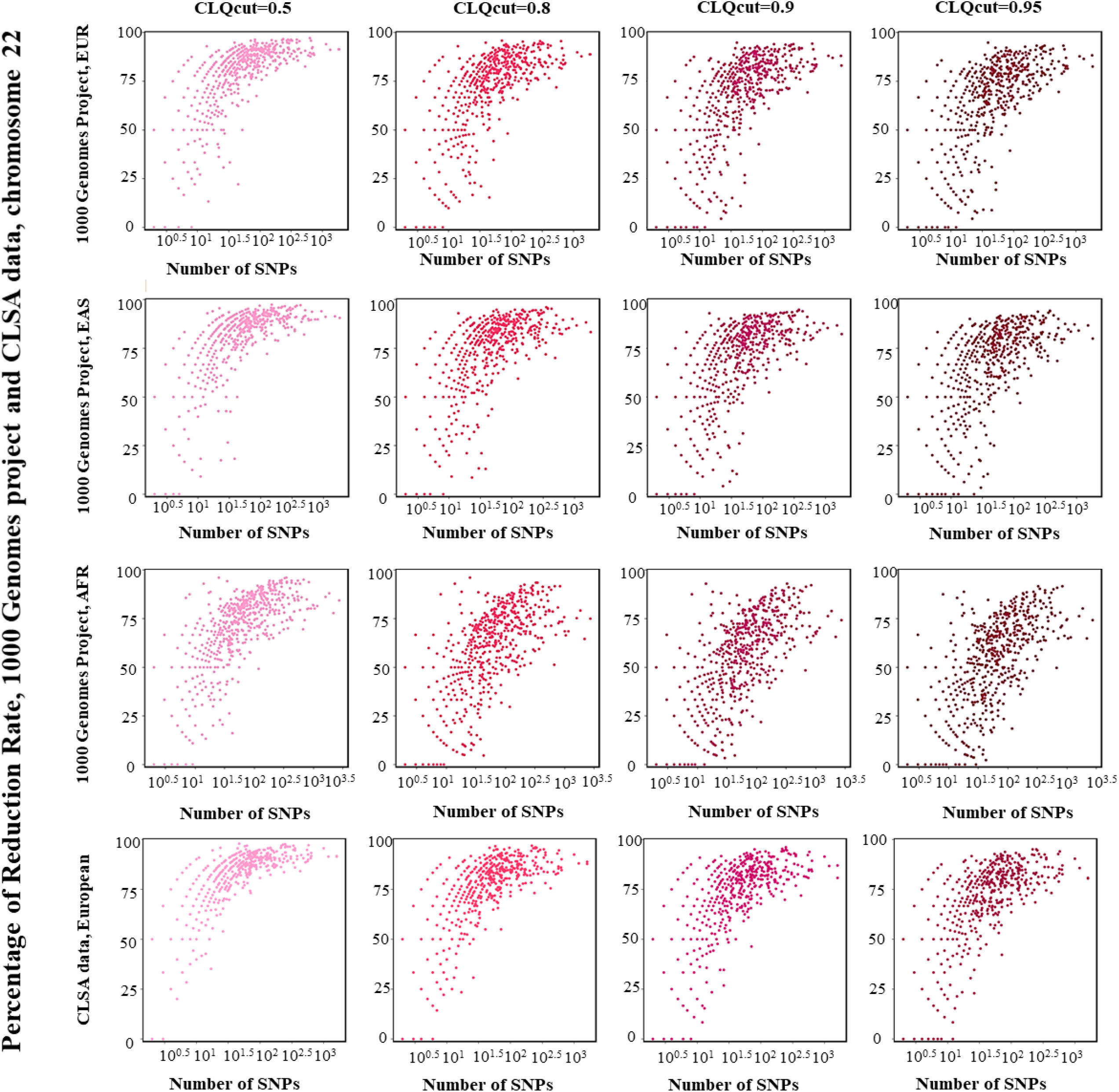
The reduction rates after the DRLPC process plotted with the number of SNPs in the original dataset (gene size) for gene-based datasets of 1000 Genomes Project and CLSA data, chromosome 22, with four threshold values for CLQcut (0.5, 0.8, 0.9, 0.95), a threshold value 0.8 for PCcut, and for three super-populations (EUR, EAS, AFR) and European ancestry.

In addition to dimension reduction, as previously noted, DRLPC aims to resolve multi-collinearity. To demonstrate the effectiveness of DRLPC, we compared the highest VIF values for the remaining variables at each step of DRLPC in gene regions. Table 4 illustrates the average and standard deviation of the highest VIF values of each step of DRLPC over gene regions of three super-populations. According to the DRLPC algorithm, at the first step, aliased variables (variables with complete dependency) were removed from the data; as shown in Table 3, the average of VIF values remained significantly high, which is a warning sign of extreme multi-collinearity. It is essential to highlight that the number of variables in steps 2 and 3 remains consistent; however, in step 3, bins(groups) are replaced by the first principal component. DRLPC can effectively address the multi-collinearity by replacing highly correlated SNPs with local PCs at step 3. As shown in Table 4, applying LPCA in step 3 partially resolves multi-collinearity. The average highest VIF values for CLQcut of 0.5 are reduced to the values below the VIF threshold (VIFcut=20) for all three super-populations. The VIF reduction^3^ of the highest VIF values from steps 1 to 3(2), ranging from 89.0% to 99.89%, suggests that incorporating the LPC has effectively mitigated issues related to multi-collinearity in the dataset. At step 4, the average of the highest VIF values is reduced to the value below the VIF threshold for all three super-populations for all CLQcut and PCcut values, underscoring DRLPC’s capability to resolve multi-collinearity effectively. As indicated in Table 4, the average of the highest VIF at step 5 with a PCcut value of 0.8 falls below the VIF threshold value for EUR and AFR. However, with a PCcut of 0.9, the average of the highest VIF increases and surpasses the threshold, particularly with CLQcut value of 0.5 for EUR and EAS. It may be ascribed to the higher average number of new variables (RPCs) using a larger PCcut value (Supplementary Table S6). For AFR, the average of the highest VIF values remains the VIF threshold when considering all CLQcut and PCcut values; however, similar to EUR and EAS, an increase in PCcut value corresponds to a rise in the average of the highest VIF. Elevating the average of the highest VIF by increasing the CLQcut values, can be attributed to the clique-based clustering algorithm’s tendency to form clusters with fewer SNPs, therefore less dimension reduction, using a larger CLQ threshold (Yoo et al., 2015). No significant difference was observed when using different PCcut values across populations for a specific CLQcut value except for step 5. The findings underscore the effectiveness of DRLPC in successfully mitigating multi-collinearity before conducting regression analysis (refer to Supplementary Excel file, Table S24 to S47 for details).

**TABLE 4.** The average and standard deviation of the highest VIF values after each DRLPC step obtained over gene regions in chromosome 22 using 1000 Genomes Project data of three super-populations. The VIF (average, SD) of aliased variables at step 1 for EUR, EAS, and AFR are (10572.5, 175545.0), (16114.1, 371104.1), (11300.5, 93899.4), respectively, and are same for using all CLQcut and PCcut value combinations.

|  | CLQcut | PCcut | Steps 2 & 3 |  | Step 4 |  | Step5 |  |
| --- | --- | --- | --- | --- | --- | --- | --- | --- |
|  |  |  | Average | SD | Average | SD | Average | SD |
| EUR | 0.5 | 0.8 | 11.3 | 21.1 | 5.6 | 5.6 | 8.6 | 13.2 |
|  |  | 0.9 | 11.3 | 21.1 | 5.6 | 5.6 | 25.3 | 374.6 |
|  | 0.8 | 0.8 | 60.0 | 117.8 | 8.7 | 6.6 | 9.4 | 8.0 |
|  |  | 0.9 | 60.0 | 117.8 | 8.7 | 6.6 | 20.3 | 258.5 |
|  | 0.9 | 0.8 | 214.2 | 2850.0 | 9.6 | 6.6 | 10.4 | 16.6 |
|  |  | 0.9 | 214.2 | 2850.0 | 9.6 | 6.6 | 11.1 | 17.5 |
|  | 0.95 | 0.8 | 1158.4 | 26297.8 | 10.9 | 6.7 | 11.8 | 23.2 |
|  |  | 0.9 | 1158.4 | 26297.8 | 10.9 | 6.7 | 13.1 | 24.9 |
| EAS | 0.5 | 0.8 | 9.5 | 17.5 | 4.6 | 5.1 | 21.3 | 322.2 |
|  |  | 0.9 | 9.5 | 17.5 | 4.6 | 5.1 | 24.7 | 323.0 |
|  | 0.8 | 0.8 | 53.0 | 88.0 | 8.5 | 17.1 | 9.1 | 17.4 |
|  |  | 0.9 | 53.0 | 88.0 | 8.5 | 17.1 | 10.4 | 18.6 |
|  | 0.9 | 0.8 | 89.2 | 126.9 | 9.2 | 6.6 | 9.8 | 12.7 |
|  |  | 0.9 | 89.2 | 126.9 | 9.2 | 6.6 | 10.6 | 13.7 |
|  | 0.95 | 0.8 | 133.4 | 249.4 | 11.5 | 22.9 | 11.8 | 23.2 |
|  |  | 0.9 | 133.4 | 249.4 | 11.5 | 22.9 | 12.7 | 24.0 |
| AFR | 0.5 | 0.8 | 18.9 | 40.3 | 8.3 | 15.8 | 12.7 | 27.5 |
|  |  | 0.9 | 18.9 | 41.1 | 8.3 | 19.9 | 17.1 | 39.4 |
|  | 0.8 | 0.8 | 112.0 | 539.5 | 12.4 | 22.1 | 14.5 | 31.0 |
|  |  | 0.9 | 112.1 | 539.9 | 12.4 | 22.1 | 16.9 | 32.0 |
|  | 0.9 | 0.8 | 200.1 | 850.8 | 14.2 | 23.7 | 16.3 | 36.6 |
|  |  | 0.9 | 200.1 | 850.8 | 14.2 | 23.7 | 17.8 | 37.9 |
|  | 0.95 | 0.8 | 354.6 | 1827.7 | 14.4 | 29.4 | 16.2 | 32.2 |
|  |  | 0.9 | 354.6 | 1827.7 | 14.4 | 29.4 | 18.0 | 34.4 |

#### 3.2.2 Dimension reduction results of CLSA dataset

Application of the DRLPC algorithm to SNP sets in gene regions, reduced the average dimension of genotype data to 7∼14%, depending on the threshold values (Figure 4 and Supplementary Table S3). We also observed that lower CLQ threshold values yield more dimension reduction than higher CLQ cut points, and larger genes had greater reduction than smaller genes (Figure 4, Table 5, Supplementary Table S4). The European ancestry results obtained from the CLSA data align consistently with the findings from the 1000 Genomes Project EUR population, revealing a comparable pattern. When the DRLPC algorithm is applied to each of the 100 consecutive SNP sets, the dimension of genotype data was reduced to 11∼21% depending on the threshold values used (Supplementary Table S4). With lower CLQcut threshold values, the dimension of 100 SNP sets was more reduced than with the higher CLQcut values. With the lower PCcut value (0.8), the percentages of final variables after the DRLPC process were applied were slightly lower than those with a PCcut of 0.9. Similar to previous results for 1000 Genomes Project data, a noticeable ascending pattern suggests a corresponding elevation in the reduction rate by increasing gene size (Figure S2). Additionally, the choice of different CLQcut values has a negligible effect on the identified pattern; moreover, regardless of CLQcut variations, the pattern remains steady.

**TABLE 5.** Average reduction rate (percentage) by DRLPC compared to the original number of SNPs in the gene regions (average number of SNPs per gene is 71.6) of European ancestry for CLSA population, chromosome 22.

|  |  | Average reduction rate (%), European, CLSA |  |
| --- | --- | --- | --- |
| CLQcut | PCcut | <sup>†</sup>Below average<br>(454 genes) | <sup>‡</sup>Above average<br>(157 genes) |
| 0.5 | 0.8 | 67.3 | 90.6 |
|  | 0.9 | 67.2 | 90.3 |
| 0.8 | 0.8 | 59.7 | 86.4 |
|  | 0.9 | 59.6 | 86.2 |
| 0.9 | 0.8 | 56.0 | 83.9 |
|  | 0.9 | 55.9 | 83.7 |
| 0.95 | 0.8 | 53.4 | 82.3 |
|  | 0.9 | 53.2 | 82.1 |

<sup>†</sup>Below: Below the average number of SNPs (The average number of SNPs is 71.6); <sup>‡</sup>Above: Above the average number of SNPs (The average number of SNPs is 71.6).

**FIGURE 4.**
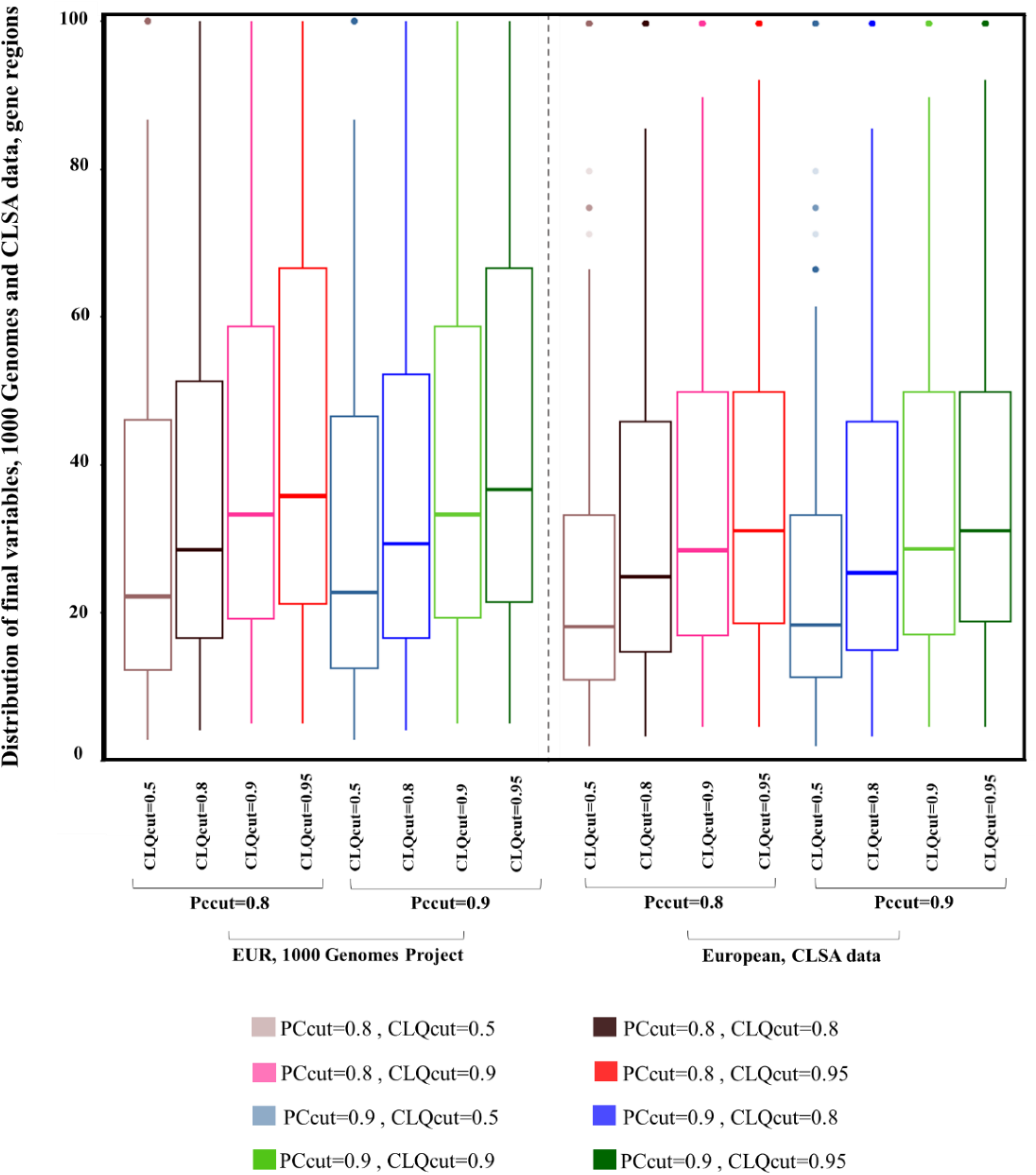
The percentage distribution of final variables after the DRLPC process compared with the original data of gene regions, with four threshold values for CLQcut (0.5, 0.8, 0.9, 0.95) and two threshold values (0.8, 0.9) for PCcut, 1000 Genomes project for EUR and CLSA data European ancestry.

Similar reduction rate patterns persist across various threshold values when examining both the 1000 Genomes Project EUR population and the CLSA data of European ancestry (Figure 3). Following the approach used in the 1000 Genomes Project data, we also compared the reduction rates resulting from the DRLPC process after dividing genes into two groups based on the number of SNPs: genes with the number of SNPs below the average and those above the average. As shown in Table 5, the average number of SNPs per gene was 71.6, the differences in average reduction rates are about 23∼28% between the bigger group (above the size average) and the smaller group (below the size average), as presented in Table 5. On average, the dimension reduction rates for genes with more SNPs were around 86%. The corresponding rates for genes with fewer SNPs were around 60%. Based on the results presented in Tables 2 and 5, it becomes evident that the EUR population in the 1000 Genomes Project and the European ancestry within the CLSA dataset exhibit a striking similarity in the observed reduction rates. Notably, this similarity holds across varying CLQcut and PCcut thresholds, with the highest reduction rates consistently occurring when both groups (below and above the size average) employ CLQcut 0.5 and PCcut 0.8. The values obtained for these thresholds are not only in the same range but also reflect the maximum reduction rates among the thresholds considered. The reduction rate for the smaller size group (below the size average) is 47.1∼61.2% and 53.2∼67.3% for EUR 1000 Genomes Project and European CLSA data, respectively. On the other hand, the reduction rate for the bigger group (above the size average) is 79.8∼89.3% and 82.1∼90.6% for EUR 1000 Genomes Project and European CLSA data, respectively. This close alignment of results underscores a similarity in the reduction rate trends observed across these two datasets.

In evaluating the efficacy of DRLPC to address multi-collinearity within the CLSA data, similar to our approach with the 1000 Genomes Project, we focused on the average of highest VIF values on gene regions of chromosome 22 (refer to Supplementary Table S5 and Supplementary Excel file Table S48 to S55). Our findings consistently correspond to those obtained from the 1000 Genomes Project, specifically within the EUR population and CLSA dataset. The VIF reduction from steps 1 to 3 (step 2) is around 99%, highlighting the effectiveness of the local Principal Component in revolving multi-collinearity in the dataset. Upon implementing LPCA at step 3, The average highest VIF values for CLQcut of 0.5 are reduced to the values below the VIF threshold (VIFcut=20). However, in harmony with the 1000 Genomes Project data observations, the average of the highest VIF values at step 4 consistently are reduced to the values below the VIF threshold across all CLQcut and PCcut values. In Step 5, the average of the highest VIF values remains below the threshold for both CLQcut values. The difference observed between the CLSA data and the 1000 Genomes Project data in Step 5 may be attributed to the lower average of adding new variables for the CLSA data in this step. However, considering the results in steps 3 and 4, the VIF findings substantiate the effectiveness of DRLPC in successfully mitigating multi-collinearity for both datasets, emphasizing the significance of this approach before conducting regression analyses, as detailed in earlier discussions. It should be mentioned that the average number of aliased variables in the CLSA data is notably lower compared to the results observed for the EUR population in the 1000 Genomes Project (Table S6 and S7). Several factors contribute to this disparity. The CLSA dataset encompasses imputed genetic data for European ancestry. Imputed genotype data undergoes rigorous quality control procedures to ensure high imputation accuracy. By selecting the high-quality imputed variants, poorly imputed variants that would have otherwise been aliased in the genotype data are effectively excluded from the analysis. Furthermore, it is essential to note that our study exclusively considered well-imputed SNPs. Another influential factor to consider is the discrepancy in the number of individuals between the two datasets. The 1000 Genomes Project dataset is limited to 503 individuals within the EUR population, while the CLSA dataset boasts a much larger cohort of 17,779 individuals.

The dimension reduction was higher for larger size genes across all three super-populations in this study, although there is a complex relationship between gene size and recombination rate since there are several factors that influence recombination rates, it could be attributed to the construction of robust LD blocks of highly correlated SNPs by the clique-based algorithm (Yoo et al., 2015). The structure of LD is influenced by various factors, including recombination, mutation, selection and population history, and genetic drift (Pritchard et al., 2001; McVean et al., 2004). Halldorsson et al., (2019) found that recombination rates are lower in genic regions (defined by the beginning of the first exon of a gene to the end of the last) than in noncoding regions. These results strongly contradict with earlier reports (Eyre-Walker, 1993; Kong et al., 2002) of a positive correlation between gene density and recombination rate. The apparent contradiction may be explained if recombination hotspots are more likely to occur near genes than within them. The relationship between recombination rate and LD is generally inverse; lower recombination rates are associated with higher LD.

## 4 SIMULATION STUDY

### 4.1 Methods

To investigate the impact of DRLPC on statistical performance in regression-based multi-SNP statistics, we designed simulation studies based on observed human genotypes from the 1000 Genomes Project (phase 3 chromosome 22 for three super-populations (EUR, EAS, and AFR), as in section 3). Quantitative traits were generated in three series of simulations of gene-level association in 100 genes. One series assumed a global null effect model (Model 1) to evaluate type I error, and two series assumed alternative trait models with one or two causal SNPs per gene (Models 2 and 3) to evaluate power gene-based tests, with and without DRLPC processing.

#### 4.1.1 Genotype data

To identify genes for multi-SNP association test evaluation, we first excluded rare and low frequency SNPs (MAF< 0.05), pruned SNPs in complete LD, and removed multi-allelic SNPs. This produced 775, 759, and 803 genes for EUR, EAS, and AFR, respectively, each with at least one SNP (https://www.ensembl.org/). From these, we selected genes with number of SNPs per gene ranging from *m*=11 to 500, resulting in 462, 454, and 498 genes for EUR, EAS, and AFR, respectively. Finally, we randomly selected 100 genes in common from 742 genes across all three super-populations, to compare the performance of gene-based tests under realistic gene structure by simulation studies (Supplementary Excel file Tables S21 to S23). Considering EUR, EAS, and AFR separately, including 503. 504, and 661 individuals respectively, the CLQD algorithm was applied with CLQcut = 0.5 threshold value to assign all *m* SNPs in a gene into mutually-exclusive clusters of varying size and number, according to the within-gene LD structure.

#### 4.1.2 Simulation model

To better understand the data and model characteristics that influence the performance of DRLPC method, we conducted a simulation study based on observed human genotypes. Quantitative trait *Y* values were generated for each gene under null and alternative hypotheses using genotype data from the 1000 Genomes Project phase 3 for three super-populations assuming an additive genetic model with *t* causal SNPs, as described below:

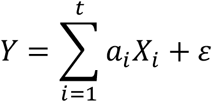

where *t* is the number of causal SNPs per gene, *a*_*i*_ is the effect of the *i*^th^ causal SNP, *X*_*i*_ is the number of minor alleles at the *i*^th^ SNP, and *ε* is the error term considered to follow a normal distribution with mean 0 and variance *σ*^2^. We considered three different quantitative trait models for each gene, including 0, 1, or 2 causal SNPs per gene (Table 6). Under the null hypothesis of no gene effect (Model 1), all *a*_*i*_ were specified to be null in the trait generation model. Under the alternative hypothesis (Models 2 and 3), non-null SNP effects were specified as: a) 1causal model with one causal SNP per gene has effect *a*_1_ = 1 (*t*=1), or b) 2causal model with two causal SNPs per gene has effects *a*_1_ = 1 and *a*_2_ = 1 (*t*=2). Under Model 2, one SNP in a gene was randomly selected to be causal. Under Model 3, a second SNP was also selected to be causal. If there was only one cluster in a gene, the second SNP was randomly chosen from the same cluster, and if there was more than one cluster in a gene, the second SNP was randomly selected from a different cluster.

**TABLE 6.** Quantitative trait models used to generate phenotypes for type I error and power comparisons of multi-SNP tests.

| Model | Explanation | <sup>†</sup>Trait model parameters |
| --- | --- | --- |
| 1 | No SNP association | All zero |
| 2 | One causal SNP within a gene (1causal) | $a_1 = 1$ |
| 3 | Two causal SNPs, both deleterious (2causal) | $a_1 = 1, a_2 = 1$ |

<sup>†</sup>The trait model is $Y = \sum_{i=1}^t a_i X_i + \varepsilon$ where $\varepsilon = N(0, \sigma^2)$ , $t$ is the number of causal SNPs, $a_i$ is the effect of the $i^{\text{th}}$ causal SNP, and $X_i$ is the number of risk alleles for the $i^{\text{th}}$ causal SNPs. The variance $\sigma^2$ is selected to maintain the power of the Wald test at 60% for each set of causal SNPs for 1causal and 2causal models.

To estimate empirical type I error, we generated 1000 replicated datasets for each gene under Model 1, and applied the analysis methods described in section 4.1.3 in each replicate, including all the SNPs and their cluster information in the regression analyses. The proportion of replicates in which the null hypothesis was rejected was then averaged over all genes, and then over subsets of genes stratified according to the original number of SNPs in the gene, and for each gene, all SNPs were included in the regression analysis. For power estimation under the alternative, we generated 1000 replicated datasets under Models 2 and 3, and similarly tabulated and averaged the per gene rejection rates in analyses that considered all SNPs. Assuming CLQcut 0.5 and PCcut 0.8, two causal SNPs were selected from different LPCs if possible. In this study, the error variance *σ*^2^ was adjusted separately for each gene and trait model to achieve a 60% power in the Wald test. The error variance was estimated using the original genotype variables in a set of 1,000 replicates under the alternative model, and regressions that include all causal and non-causal SNPs.

#### 4.1.3 Multi-marker test statistics

Whenever a gene includes several SNPs, multi-SNP analysis can be applied by multiple regression with multi-parameter hypotheses or by incorporating single-SNP marginal regression analysis results. Both approaches demand coded genotype data. In order to evaluate the impact of DRLPC on regression-based multi-SNP statistics, we selected several multi-marker statistics based on joint or marginal regression to compare the power using original data and dimension-reduced data by DRLPC. Among joint regression tests, Wald (Wald, 1943), Multiple linear combination (MLC) (Yoo et al., 2017), and PC80 tests (Gauderman et al., 2007) are evaluated in this study. Furthermore, the sequence kernel association (SKAT) and SKAT-O tests (Ionita-Laza et al., 2013; Lee et al., 2012), are included as well-known gene-based tests of SNP sets for gene-based association analysis. The MLC test is derived from the joint regression Wald statistics by applying a set of linear contrasts to the multi-SNP regression parameters that reduce the dimension (df) of the test statistic. The contrasts reflect the cluster membership determined using the CLQ algorithm prior to regression estimation (Yoo et al., 2015, Yoo et al., 2017). CLQ optimizes within-cluster correlation using pairwise correlation of additively coded SNPs, with SNP recoding as necessary for positive within-cluster correlation. The weights in the multiple linear combinations are derived from the regression variance-covariance matrix which depends on MAFs and LD among the SNPs in the gene. We considered two types of MLC tests: MLC-B (based on the beta coefficients) and MLC-Z (comparable Z statistics test, *Z* = (*Z*_1_, *Z*_2_, …, *Z*_*m*_)^*T*^), considering two CLQcut threshold values, 0.5 and 0.8, respectively. The details of all tests are available in Supplementary Methods.

### 4.2 Results

#### 4.2.1 Evaluation of type I error rate

We report the type I error estimates of each statistic using 10,000 replications considering two nominal critical values for *α* = 0.05 and 0.01 and averaging across 100 genes (Table 7, Supplementary Table sS8 and S9, Figures S3 and S4). For the simulation study, two CLQcut (0.5 and 0.8) and one PCcut (0.8) threshold values are selected to obtain type I error (and empirical power) for all gene-based tests using the DRLPC method. As shown in Table 7, the average empirical type I error for the Wald test was elevated in the original data and declined from 0.07 by 0.02 in the DRLPC processed data, and close to the nominal 0.05 level for all three super-populations under the 1causal model. The average standard deviation and average df for the Wald test also decreased considerably with application of DRLPC for both CLQcut points. On average, all other test statistics exhibited type I error control in original and DRLPC analysis. We also observed greater type I error inflation with larger genes for some tests, particularly for the Wald test (Supplementary Tables S10 to S12 and Figures S5 to S7). However, applying the DRLPC decreased the inflation, resulting in values close to the nominal 0.05 level for three super-populations under the 1causal model. The average and SD of MLC-B tests vary little across the CLQ threshold values, suggesting that clustering and dimension reduction do not affect standard error estimates. Comparing the obtained empirical type I error values between populations demonstrates a high similarity between the results. It can be inferred that the implementation of DRLPC has reduced the type I error values for all three super-populations.

**TABLE 7.** Empirical type I error of gene-based statistics (N=10,000 replicates) at the nominal level α= 0.05, averaged over 100 genes, using original data and two DRLPC processed data for three super-populations.

| Population | Statistics | Original data |  |  | CLQcut 0.5 |  |  | CLQcut 0.8 |  |  |
| --- | --- | --- | --- | --- | --- | --- | --- | --- | --- | --- |
|  |  | Average | SD | Average df | Average | SD | Average df | Average | SD | Average df |
| EUR | Wald | 0.071 | 0.010 | 35.7 | 0.052 | 0.003 | 9.0 | 0.053 | 0.003 | 12.6 |
|  | PC80 | 0.051 | 0.002 | 4.2 | 0.051 | 0.002 | 4.8 | 0.052 | 0.002 | 5.3 |
|  | MLC-B5 | 0.053 | 0.003 | 8.9 | 0.052 | 0.003 | 7.3 | 0.052 | 0.003 | 8.1 |
|  | MLC-B8 | 0.055 | 0.004 | 14.5 | 0.052 | 0.003 | 8.7 | 0.053 | 0.003 | 11.9 |
|  | SKAT | 0.049 | 0.003 | - | 0.049 | 0.003 | - | 0.050 | 0.002 | - |
|  | SKAT-O | 0.052 | 0.003 | - | 0.052 | 0.003 | - | 0.052 | 0.003 | - |
| EAS | Wald | 0.069 | 0.012 | 33.3 | 0.053 | 0.002 | 7.9 | 0.053 | 0.003 | 11.2 |
|  | PC80 | 0.051 | 0.002 | 3.7 | 0.052 | 0.002 | 4.6 | 0.052 | 0.002 | 4.9 |
|  | MLC-B5 | 0.053 | 0.003 | 7.9 | 0.052 | 0.003 | 6.4 | 0.052 | 0.003 | 7.1 |
|  | MLC-B8 | 0.055 | 0.004 | 7.9 | 0.053 | 0.003 | 7.6 | 0.053 | 0.003 | 10.3 |
|  | SKAT | 0.049 | 0.002 | - | 0.049 | 0.002 | - | 0.050 | 0.003 | - |
|  | SKAT-O | 0.052 | 0.003 | - | 0.052 | 0.003 | - | 0.052 | 0.003 | - |
| AFR | Wald | 0.076 | 0.015 | 67.8 | 0.054 | 0.003 | 18.1 | 0.055 | 0.004 | 26.4 |
|  | PC80 | 0.052 | 0.002 | 7.4 | 0.052 | 0.002 | 7.9 | 0.052 | 0.002 | 8.8 |
|  | MLC-B5 | 0.054 | 0.003 | 18.6 | 0.053 | 0.003 | 14.0 | 0.053 | 0.003 | 16.2 |
|  | MLC-B8 | 0.057 | 0.005 | 31.5 | 0.054 | 0.003 | 17.8 | 0.055 | 0.004 | 25.4 |
|  | PC80 | 0.052 | 0.002 | 7.4 | 0.052 | 0.002 | 7.9 | 0.052 | 0.002 | 8.8 |
|  | SKAT | 0.050 | 0.002 | - | 0.050 | 0.002 | - | 0.050 | 0.002 | - |
|  | SKAT-O | 0.052 | 0.003 | - | 0.052 | 0.003 | - | 0.052 | 0.003 | - |

<sup>†</sup>Used data: Original data; <sup>‡</sup>CLQcut 0.5: DRLPC processed data using CLQ threshold value 0.5; <sup>§</sup>CLQcut 0.8: DRLPC processed data using CLQ threshold value 0.8. <sup>¶</sup>List of test statistics: Wald: generalized Wald test (Wald, 1943); PC80: global test on regression using the minimum number of principal components capturing 80% of variance (Gauderman et al, 2007); MLC test: Multi Linear combination test (Yoo et al, 2017); MLC-B5: MLC tests using beta coefficient by considering CLQcut equal to 0.5, MLC-B8: MLC tests using beta coefficient by considering CLQcut equal to 0.8; SKAT: sequence kernel associated tests for the common variants (Ionita-Laza et al, 2013); SKAT-O: a linear combination of SKAT and burden test with optimized mixing proportion (Lee et al, 2012).

#### 4.2.2 Comparison of empirical power values for original data versus DRLPC processed data

To evaluate the efficiency of the DRLPC method, empirical power values of each of the multi-SNP statistics were estimated using 1000 replications for each of 100 genes using the original data and two DRLPC processed data for two trait models (Table 8, Supplementary Table S13, Supplementary Figures S8 to S10). We computed power as the proportion of replicates with p-values below the threshold corresponding to a nominal critical value for *α* = 0.05 and obtained average and standard deviation of empirical power across all gens under two trait models in three super-populations (Supplementary Table S14 provides similar results for 2causal model, Supplementary Tables S56 to S61 provide results for each gene for three super-populations).

**TABLE 8.** Empirical power (percentage) of gene-based statistics (N=1,000 replicates) at the 0.05 level for three populations, 1causal model, averaged over 100 genes.

| Population | Statistics | Original data |  |  | CLQcut 0.5 |  |  | CLQcut 0.8 |  |  |
| --- | --- | --- | --- | --- | --- | --- | --- | --- | --- | --- |
|  |  | Average | SD | Average df | Average | SD | Average df | Average | SD | Average df |
| EUR | Wald | 60.3 | 4.6 | 35.7 | 81.3 | 9.5 | 9.0 | 78.7 | 8.2 | 12.6 |
|  | PC80 | 82.1 | 18.4 | 4.2 | 81.4 | 12.7 | 4.8 | 81.9 | 11.3 | 5.3 |
|  | MLC-B5 | 81.3 | 8.6 | 8.9 | 82.2 | 11.2 | 7.3 | 82.7 | 9.5 | 8.1 |
|  | MLC-B8 | 77.0 | 7.8 | 14.5 | 81.8 | 9.5 | 8.7 | 79.4 | 8.6 | 11.9 |
|  | SKAT | 76.8 | 20.5 | - | 80.5 | 18.0 | - | 76.1 | 25.0 | - |
|  | SKAT-O | 74.6 | 22.0 | - | 77.6 | 19.5 | - | 74.0 | 25.0 | - |
| EAS | Wald | 60.5 | 4.0 | 33.3 | 82.3 | 8.9 | 7.9 | 79.6 | 7.6 | 11.2 |
|  | PC80 | 84.5 | 14.0 | 3.7 | 81.5 | 13.4 | 4.6 | 82.9 | 12.2 | 4.9 |
|  | MLC-B5 | 81.3 | 11.5 | 7.9 | 81.1 | 14.8 | 6.4 | 81.0 | 13.3 | 7.1 |
|  | MLC-B8 | 78.2 | 7.7 | 7.9 | 82.8 | 8.6 | 7.6 | 80.4 | 7.9 | 10.3 |
|  | SKAT | 79.5 | 18.5 | - | 80.0 | 18.2 | - | 74.6 | 22.2 | - |
|  | SKAT-O | 76.6 | 20.6 | - | 78.1 | 18.8 | - | 73.0 | 22.8 | - |
| AFR | Wald | 60.5 | 3.9 | 67.8 | 82.8 | 8.7 | 18.1 | 79.3 | 8.0 | 26.4 |
|  | PC80 | 85.8 | 11.9 | 7.4 | 82.7 | 12.7 | 7.9 | 85.5 | 9.3 | 8.8 |
|  | MLC-B5 | 82.8 | 8.2 | 18.6 | 83.6 | 9.4 | 14.0 | 83.3 | 10.2 | 16.2 |
|  | MLC-B8 | 77.0 | 7.0 | 31.5 | 82.9 | 8.7 | 17.8 | 80.0 | 8.2 | 25.4 |
|  | SKAT | 79.6 | 17.2 | - | 81.5 | 16.2 | - | 72.5 | 27.5 | - |
|  | SKAT-O | 74.7 | 19.3 | - | 77.8 | 17.8 | - | 69.7 | 27.9 | - |

As shown in Table 8, it is noteworthy that Wald test average power remarkably improves by 20% using DRLPC processed data compared to using original data for all three super-populations. As expected, CLQcut 0.5 produces larger clusters and smaller df than CLQcut 0.8. The PC80 achieves similar power using the original data compared to DRLPC for both CLQcut values in all three super-populations. The PC80 DRLPC processed data and original data results are remarkably similar on average. Nevertheless, it is worth mentioning that not all (global) principal components are readily interpretable in a biological context, and choosing a subset of principal components might lead to the exclusion of meaningful information; hence, the results obtained with the DRLLPC may offer a more dependable basis for interpretation.

MLC-B performs dimension reduction by constructing a weighted linear combination for each cluster using the original multiple regression coefficients, while DRLPC reduces dimension by constructing a new variable which is a weighted linear combination of the genotypes within each cluster and then performs, multiple regression with the new variables, but the weights differ. Nevertheless, the average power of MLC-B obtained using the original data is similar to the power of Wald using DRLPC processed data for corresponding CLQcut values. The DRLPC process slightly enhanced the power of MLC-B compared to using the original data in every population when the CLQ threshold was 0.8.

For the 1 causal model, SKAT usually has higher average power than SKAT-O, especially using CLQ cut-point 0.5 across three super-populations. While analyzing the 2causal model (Supplementary Table S14), we observed that SKAT and SKAT-O had usually higher power using the original data for all three populations. Since the effects of both causal variants for the 2causal model are in the same direction, SKAT usually has a higher average power than SKAT-O for all populations.

The average power of each test by applying DRLPC was higher than 70% for three populations under both trait models. Based on the result in Table 8, we can infer that the power of DRLPC processed data tends to increase when the degrees of freedom (df) of each test decrease, implying that DRLPC can enhance power by reducing the dimension of the data. The average power obtained by DRLPC is higher considering CLQcut 0.5 than CLQcut 0.8. under each trait model for all tests (Table 8 and Supplementary Tables S13 and S14).

For each population, we also compared the median and interquartile range (IQR) of empirical power for 100 genes, stratified into three groups based on the number of SNPs: genes with less than 100 SNPs, 101∼200 SNPs, and more than 200 SNPs (Figure S11 reports distributions of the number of SNPs per gene). As shown in Figure 5 (Supplementary Tables S15 and S16), DRLPC exhibited a strong enhancement in the power of the Wald test for larger genes, resulting in an approximately 30% increase in power with type I error control (Table 7). Based on the data application in section 3, in which the dimension reduction is greater in bigger-size genes, such results are expected. Furthermore, in contrast to the Wald test, the average power of the PC80 and MLC-B demonstrated a modest increase for larger gene sizes for CLQcut value 0.8. The results of the DRLPC processed data using the CLQcut value 0.5 of the third group for SKAT and SKAT-O have lower median the power than original data but with high variability due to the small number of genes in that group, which is 7. The average power for SKAT and SKAT-O tests using DRLPC processed data is higher than the original data using CLQcut 0.8 for the bigger-size genes; while for genes with less than 200 SNPs (groups 1 and 2), the average power is higher for DRLPC processed data than original data using CLQcut 0.5. It is worth noting that our finding remained consistent for another trait model and other populations (Supplementary Tables S17 to S20 and Figures S12 to S17 for results in EAS and AFR populations and the 2causal model), further substantiating the robustness and utility of DRLPC in various genetic association scenarios.

**FIGURE 5.**
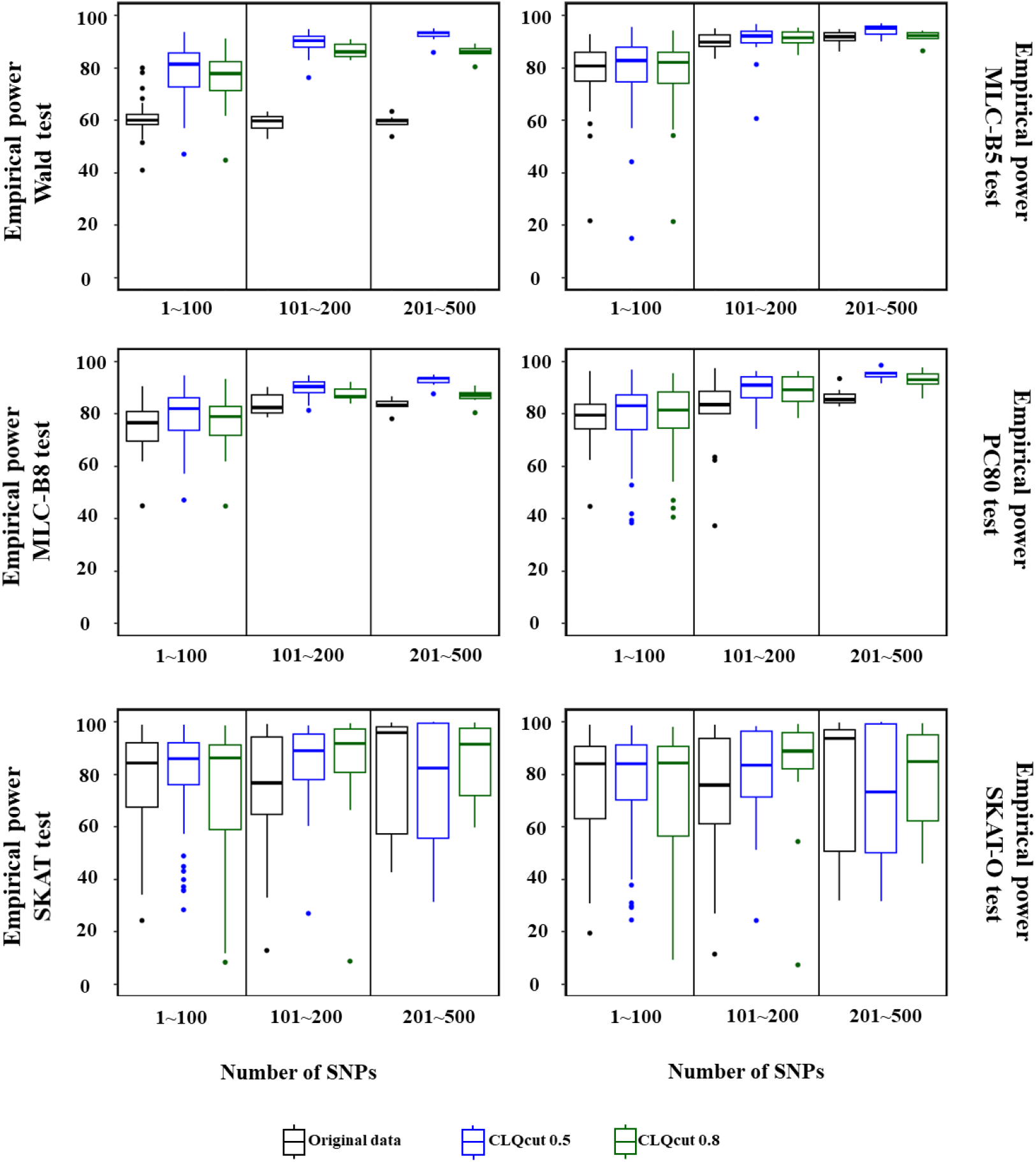
Percentage of empirical power for gene-based statistics (N=1,000 replicate) at the 0.05 level for EUR population, 1causal model, averaged across three groups of 100 genes based on the number of SNPs in the gene.

### 4.3 Runtime evaluations for DRLPC

Figure 6 presents the run-time of DRLPC when applied to gene-based SNP-set genotype data from the three super-populations in 1000 Genomes Project and the CLSA, after sorting genes based on their size using four CLQcut thresholds of (0.5, 0.8, 0.9, 0.95), as well as PCcut threshold of 0.8 (Supplementary Figure for same CLQcut values and PCcut of 0.9). It is noteworthy that computational time for different CLQcut values shows little difference between thresholds. We summarized the average run-time of DRLPC for several genes with different sizes (refer to Supplementary Excel file, Table S62 to S65 for more information), which demonstrated that the average computational time for genes with less than 500 SNPs, is around 0.06 seconds while the average computational time for genes with more than 500 SNPs is 3.41 seconds. Furthermore, the maximum run-time for larger genes (genes with more than 1000 SNPs) is less than 100 seconds for all three super-populations of 1000 Genomes Project (sample size 503-661) and 1,000 seconds for CLSA European ancestry (sample size 17,965) (Figure S18), underscoring the effectiveness of DRLPC in reducing the computational time.

**FIGURE 6.**
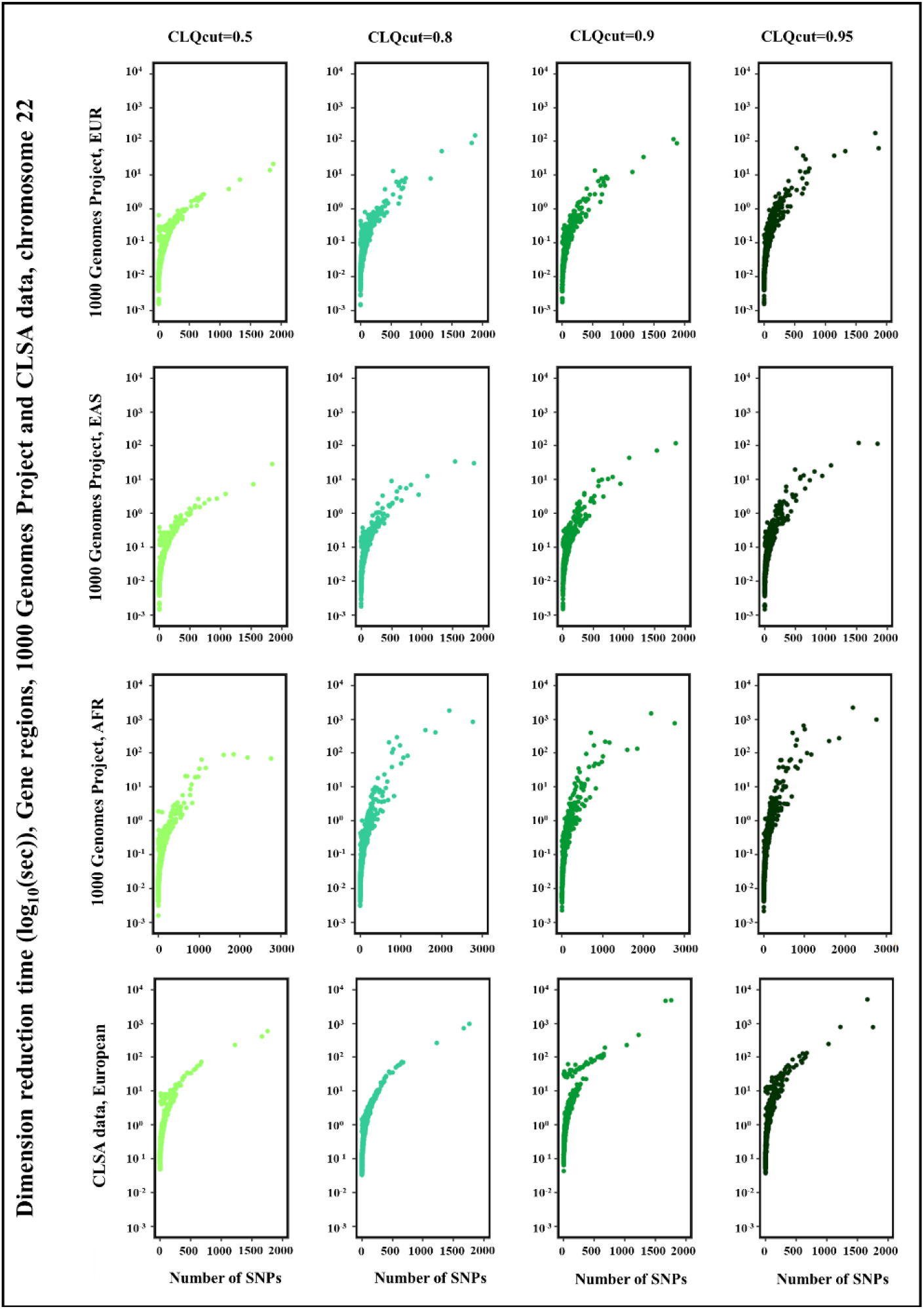
Computational time for dimension reduction using the DRLPC for gene regions (chromosome 22), with four threshold values for CLQcut (0.5, 0.8, 0.9, 0.95) and a threshold values 0.8 for PCcut, 1000 Genomes Project, three super-populations (sample size are 503-601) and CLSA data European ancestry (sample size is 17,779).

As previously discussed, our simulation study applied the DRLPC to the 1000 Genomes Project across three super-populations. Figure 7 illustrates the computational time for test statistics in a single replication using the original data and two sets of DRLPC processed data for 1causal model EUR population (Supplementary Figures S19 to S23 for 2causal model and other populations). The genes were sorted based on their size, and two CLQcut thresholds (0.5, 0.8) and a PCcut threshold of 0.8 were employed. Notably, the computational time using DRLPC decreased as gene sizes increased, particularly for genes with more than 200 SNPs, since the multiple regression dimension (df) has already been reduced by the DRLPC reduction within the clusters. This trend is consistent across various tests and populations. Additionally, using a CLQcut value of 0.5 demonstrated better computational performance, requiring approximately 10∼30% less time for larger-sized genes, to the 0.8 threshold. The Intel(R) Core(TM) i5-4200U CPU with 1.60GHz and a memory of 8.00 Gb DDR3 RAM and 238Gb local hard disk was used for the calculation.

**FIGURE 7.**
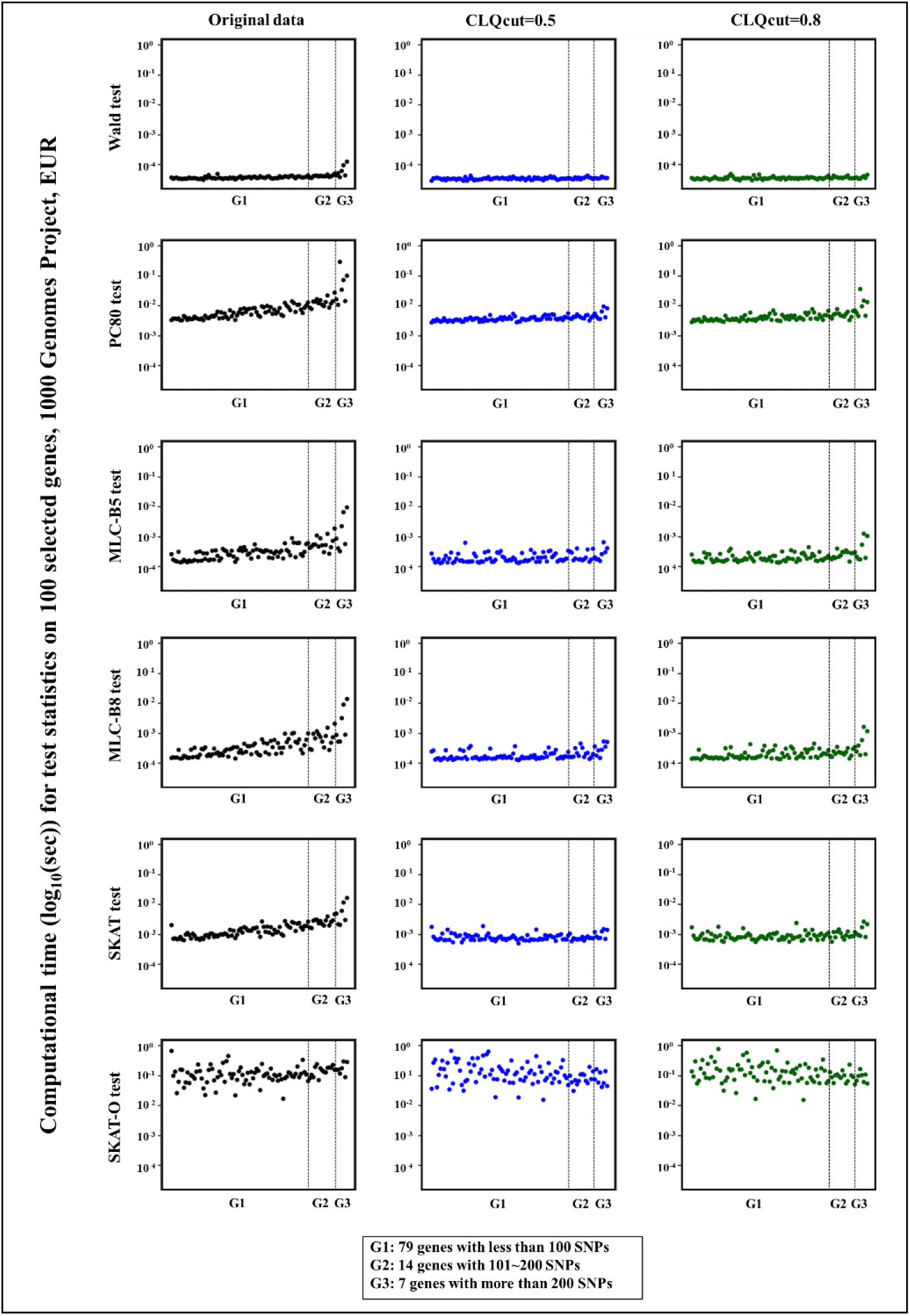
The computational time for test statistics in a single replication of 1causal model on 100 selected genes, chromosome 22: 79 genes with less than 101 SNPs, 14 genes with 101∼200 SNPs, and seven genes with more than 200 SNPs, from 1000 Genomes Project, EUR. The X-axis represents the original gene size, sorted by the number of SNPs. The original data and the DRLPC processed data were examined at two threshold values for CLQcut (0.5, 0.8) and a threshold value of 0.8 for PCcut.

## 5 DISCUSSION

By jointly analyzing multiple variants within a gene, instead of one variant at a time, gene-based multiple regression can improve power, robustness, and interpretation in genetic association analysis. Yoo et al., (2017) proposed multiple linear regression with multi-dimension Wald and reduced dimension multiple linear combination (MLC) test statistics and demonstrated multi-SNP regression-based analysis can be a well-powered and robust choice among the existing methods across a range of complex genetic architectures. Using the same LD clique-based clustering implemented to define sets of related SNPs for MLC tests (Yoo et al., 2015) and incorporating dimension reduction through LPCA in each cluster, we have proposed the DRLPC algorithm to enhance statistical validity and power of multi-SNP tests among multiple correlated genetic variants.

Dimension reduction is an approach to reduce the number of variables in a dataset while retaining as much variation in the original dataset as possible. Kambhatla et al. (1997) demonstrated that applying LPCA effectively reduces dimension in high-dimension data and relieves concerns related to multi-collinearity. Multi-collinearity occurs when there is a high level of linear dependency among regression variables. Methods proposed to resolve multi-collinearity include ridge regression (Hoerl & Kennard, 1970), partial least squares (Wold et al, 1984), lasso method (Tibshirani, 1997), principal component analysis (Pearson 1901, Hotteling 1933). While acknowledging the potential of PCA and LPCA in addressing multi-collinearity at least partially, it is important to note that these methods may not guarantee a complete solution due to their limited effectiveness in providing a comprehensive diagnosis of multi-collinearity. In this study, we introduced the DRLPC algorithm which reduces the dimension of dense sequencing data by selecting clusters with high within-cluster correlation and replacing each cluster with local principal components constructed locally among the SNP in the cluster before the regression analysis. Dimension reduction is a crucial strength of DRLPC, as it allows researchers to manage the difficulties of working with complex and highly interrelated genomic data. Incorporating the Local Principal Component (Kambhatla et al. 1997) in DRLPC facilitates the identification of the underlying genetic structure and improves the accuracy and stability of regression models. Moreover, DRLPC directly addresses the issue of multi-collinearity through a sequential two-step procedure. Initially, employing LPCA offers a degree of relief from multi-collinearity and enhances the power of regression-based multi-SNP genetic association analysis. This approach allows researchers to tackle two critical aspects simultaneously, resulting in a more efficient and comprehensive solution.

To investigate the performance of DRLPC in dimension reduction, we applied it to genotypic data from the 1000 Genomes Project for three super-populations (EUR, EAS, and AFR) and the CLSA European ancestry subset. Considering results for nearly 200 SNP sets of varying number obtained in chromosome 22, DRLPC effectively reduced dimension in all datasets. The dimension reduction rate for larger genes was around 83% for EUR and EAS, and 74% for AFR 1000 Genomes samples, and 85% for European ancestry CLSA samples. We observed less dimension reduction in AFR compared to EUR and EAS due to weaker LD in AFR (Supplementary Figures S24).

For some genes, there was a strong dependency between some SNPs before applying DRLPC, and LPCA reduced the average of VIF for the remaining variables. However, in some instances, VIF values exceeding the predefined threshold remained after applying LPCA. Subsequently, removing variables with the highest VIF (step 4) ensures that the remaining variables maintain VIF values below the threshold. By systematically eliminating high VIF values via the DRLPC framework (step 4), the average VIF for all genes descended below the predetermined threshold. The outcomes indicate that applying DRLPC yielded consistent results across populations.

To investigate the performance of DRLPC pre-processing in hypothesis testing for genetic association, we conducted simulations based on the 1000 Genomes populations to assess validity and power of several gene-level test statistics. Based on the simulation results, we conclude that the multi-SNP Wald regression test applied to the DRLPC processed data performs better than in the original data for genes with larger numbers of highly correlated SNPs. On average over 100 genes, all test statistics based on DRLPC effectively control type I errors near the nominal 0.05 level in all three super-populations. Moreover, DRLPC processing removed type I error inflation for the Wald test. This finding underscores the validity of the method.

Furthermore, the empirical power across 1,000 replications obtained for each of 100 genes in three super-populations under two trait models, indicated that the genotypic dimension reduction and the impact of DRLPC was almost identical in the two trait models for all tests. In both trait models, the Wald test with DRLPC showed the most robust efficiency, with power improved by around 20%, particularly for larger size genes. Use of the same clique-based algorithm and the same CLQcut value to create SNPs clusters for Local PCs in DRLPC and linear combination of SNPs within clusters in MLC, produces similar empirical power for the DRLPC Wald test and original MLC test.

The effect of DRLPC on PC80 was not remarkable since PC80 already achieves an acceptable power without DRLPC. Although constructing principal components from all SNP variables in a region is a common approach, interpreting them as biological entities may be challenging. It is possible that information may be lost by analyzing only a subset of principal components. On the other hand, clusters of the highly correlated SNPs produced by the clique-based algorithm and used by DRLPC and MLC retain their biological meaning.

The SKAT test is based on marginal beta coefficients and does not consider SNP covariance directly in the test statistic. Moreover, SNP LD is not considered in the linear burden test component of SKAT-O. Based on the results obtained in this study, the power for SKAT is often greater than for SKAT-O. In general, the positive impact of the DRLPC on SKAT was greater than that on SKAT-O. For SKAT and SKAT-O larger genes processed by DRLPC have lower median power than the original data, with variability attributed to the limited number of genes in this group, totaling 7. Although substantial differences were not observed between the three super-populations for the SKAT test, the power of the SKAT test was higher using the DRLPC processed data under the 1causal model compared to the 2causal model, particularly when using a CLQcut point 0.5 across all three super-populations.

We also conducted a stratified analysis by grouping genes based on their size and computing the type I error and average empirical power for each group using original data, and two DRLPC processed datasets. Notably, the number of SNPs in the gene did not substantially influence the type I error in the latter, resulting in the limited impact of this variation. Remarkably, the Wald test exhibited the most improvement. The enhancement in power through increased gene size was more conspicuous, especially with the CLQcut value set at 0.5. Therefore, we recommend the threshold value of 0.5 for DRLPC. In addition to reducing dimensionality while maintaining the interpretability of localized effects, pre-processing with DRPC offers the advantage of decreased computational time required for regression analysis. The results demonstrate that applying DRLPC elevates the statistical power of Wald tests and effectively reduces computational time.

Given mounting evidence for the role of LD structure in the effectiveness of gene-based tests, it is prudent to consider approaches like DRLPC, explicitly tailored to leverage LD information, as a viable alternative for genetic association analysis of dense genotyping data characterized by correlated SNPs and intricate LD structure.

## 6 CONCLUSIONS

In conclusion, our study has demonstrated that dimension reduction by local principal components (DRLPC) effectively reduces the dimension of high-density DNA sequencing or imputed array data and. Our results indicate that DRLPC significantly resolves multi-collinearity prior to regression analysis and improves the power obtained for the Wald test, making it an equivalent approach to the MLC test. By reducing the data dimension, DRLPC has been shown to enhance the accuracy and efficiency of multi-marker methods such as the Wald test. The simulation results strongly suggest that DRLPC has excellent potential for improving the power of SNP-based association studies. Applying DRLPC improves type I error control and enhances the statistical power of the Wald test especially (and potentially also for the MLC test) when the number of SNPs per gene is large and the sample size is relatively modest (i.e. low n/p ratio). Additionally, it reduces computational time. Our findings provide valuable insights into the use of DRLPC as a promising tool for the analysis of complex genetic data, and we hope that our study will inspire further research in this critical area.

## Supporting information

This file includes Supplementary methods, Tables S1 to S20, Figures S1 to S24.

This file includes Supplementary Tables S21 to S65.

## ACKNOWLEDGMENT

This work was supported by the National Research Foundation of Korea (NRF) [NRF-2018R1A2B6008016]. This research was made possible using the data collected by the Canadian Longitudinal Study on Aging (CLSA). Funding for the Canadian Longitudinal Study on Aging (CLSA) is provided by the Government of Canada through the Canadian Institutes of Health Research (CIHR) under grant reference: LSA 94473 and the Canada Foundation for Innovation, as well as the following provinces, Newfoundland, Nova Scotia, Quebec, Ontario, Manitoba, Alberta, and British Columbia. This research has been conducted using the CLSA dataset [Comprehensive Dataset version 4.0], under Application Number [1909028]. The CLSA is led by Drs. Parminder Raina, Christina Wolfson and Susan Kirkland. This research was partially funded by the CIHR Project Grant (#PJT-159463).

## DATA AVAILABILITY

Data are available from the Canadian Longitudinal Study on Aging (www.clsa-elcv.ca) for researchers who meet the criteria for access to de-identified CLSA data.

## FOOTNOTES

1 Dimension reduction refers to approaches to summarizing massive data such that most of the information in the data is preserved even with a smaller number of variables.

2 In regression studies, alias variables refer to variables that are highly correlated or redundant with each other.

3 The reduction in VIF is calculated as the percentage of one minus the ratio of the highest VIF at a specific step to the highest VIF at the preceding step.

