## Supplementary material for "Dimension Reduction using Local Principal Components for Regression-based Multi-SNP Analysis in 1000 Genomes and the Canadian Longitudinal Study on Aging (CLSA)": This file includes Supplementary methods, Tables S1 to S20, Figures S1 to S24.

Supplementary Methods

Supplementary Tables S1 to S20

Supplementary Figures S1 to S24

**FIGURE S18** Computational time for dimension reduction using DRLPC, PCcut value 0.9.......... **Error! Bookmark not defined.**
**FIGURE S19** Computational time for test statistics using DRLPC, EUR, 2cusal model**Error! Bookmark not defined.**
**FIGURE S20** Computational time for test statistics using DRLPC, EAS, 1cusal model**Error! Bookmark not defined.**
**FIGURE S21** Computational time for test statistics using DRLPC, EAS, 2cusal model .......... 50
**FIGURE S22** Computational time for test statistics using DRLPC, AFR, 1cusal model .......... 51
**FIGURE S23** Computational time for test statistics using DRLPC, AFR, 2cusal model .......... 52
**FIGURE S24** LD($r^2$) heatmap for three super-populations .......... 53

**REFERENCES** .......... 54

### 1 | SUPPLEMENTARY METHODS

#### 1.1 | Quality control process of the CLSA data

The quality control process of the CLSA data involved two main components: 1) the marker-based quality control analysis consisted of four tests aimed at ensuring marker consistency across various experimental factors, including genotyping batch, participant sex, Hardy-Weinberg equilibrium (HWE), and discordance of genotyping across control replicates (with a multiple-testing corrected p-value threshold for quality control tests as 3.93×10-10); 2) the sample-based quality control was conducted to identify genotyped samples of low quality, detect related individuals, and provide a genetic-based description of ancestry. A total of 794,409 genetic markers were initially selected. After passing all four tests in the marker-based quality control (removed 35,589 markers), removing markers that were not insertions/deletions, had an MAF > 0.01, and had marker-wise missingness < 0.01, the final dataset consisted of 570,275 genetic markers. Phasing and imputation[1] process were performed utilizing the Sanger imputation Service (2018). The Haplotype Reference Consortium (HRC) reference panel, which includes 64,940 haplotypes across 40,359,612 genetic markers, is employed for this process. The genotype imputation required the marker reference alleles to match the human genome GRCh37/hg19 reference sequence. To perform the imputation, a subset of the study participants (19,669 individuals) that passed sample-based quality control and a set of 719,696 markers meeting specific criteria (passing all four marker-based QC tests, SNP-wise missingness < 0.05, and MAF > 0.0001) is used as input. Applying the bcftools +fixref plugin removed 63,043 markers, yielding a total marker count of 656,653 for the imputation process markers.

We obtained the dimension reduction results of DRLPC for chromosome 22 SNPs, in consistency with the 1000 Genomes Project data used in this study, and the number of SNPs in chromosome 22 after imputation was 524,544. A set of quality-control procedures, including marker-based and sample-based quality control, was performed on CLSA data. Marker-based quality control analysis consisted of 4 tests intended to check for consistency of markers across various experimental factors, such as genotyping batch, participant sex, Hardy-Weinberg equilibrium (HWE), and discordance of genotyping across control replicates. All tests were conducted on a subset of ancestrally homogeneous participants. The Sample-based quality control aimed to identify low-quality genotyped samples, identify related individuals, and provide a genetic-based description of ancestry. Therefore, high-quality SNPs were used to ensure no bias was introduced due to the genotype batch effect or other genotyping artifacts. Same as the 1000 Genomes Project data, first, we removed SNPs with missing values and excluded multi-allelic SNPs. Moreover, we considered MAF ≥ 0.05 and INFO score ≥ 0.8 (to select the well-imputed SNPs) for European ancestry. After the filtering process, the total number of remaining SNPs was 71,695. The number of individuals in the final dataset was 17,779. In the CLSA data, m dosage variables are in the range 0~2.

---

[1] The genotype imputation is a computational method to predict marker genotypes not directly genotyped by an assay, such as a genotyping array.

#### 1.2 Gene-based tests used for Power comparison

Suppose that *m* SNPs in a gene, which were defined as $X = (X_1, X_2, \ldots, X_m)$, have been genotyped as an additive genotype model and coded as 0,1 or 2. The multi-SNP joint regression model of *m* SNPs by considering $E[Y]$ as the expected value of quantitative trait *Y* is formulated as follow:

$$E[Y] = \beta_0 + \beta_1 X_1 + \beta_2 X_2 + \cdots + \beta_m X_m$$

Global tests statistics based on the regression analysis are constructed from the beta estimates $\hat{\beta} = (\hat{\beta}_1, \ldots, \hat{\beta}_m)^T$ and associated covariance matrix $\sum_B$ .

##### *MLC-B and MLC-Z tests*

Numerous multi-marker methods based on marginal or joint association analysis have been developed to analyze rare or common SNPs. Yoo et al., (2014) proposed two multi-bin multi-marker regression tests; one is named MLC-B and the other MLC-Z. The MLC-B and MLC-Z are based on the beta coefficients and the comparable Z statistics respectively. The necessity of the MLC test is creation bins with a high correlation between SNP genotypes within a bin and a low correlation between SNP genotypes in different bins (Yoo et al., 2015). Assume that *L* bins of SNPs have been constructed in high LD (Linkage disequilibrium). The MLC-B test can construct by using beta coefficient $\hat{\beta} = (\hat{\beta}_1, \ldots, \hat{\beta}_m)^T$ and the covariance matrix $\Sigma_B$ with a weight matrix $W_s$. MLC-B is formulated as follow:

$$MLC - B = (\hat{\beta}^T W_s)(W_s^T \textstyle\sum_B W_s)^{-1}(W_s^T \hat{\beta}),$$

Where $W_s = (\sum_B^{-1} . J)(J^T . \sum_B^{-1} . J)^{-1}$ and *J* is a *m* by *L* matrix which $J_{ij}$ =1 if $i^{th}$ SNP belongs to the $j^{th}$ bin otherwise $J_{ij}$=0.

To construct MLC-Z, the standardized test statistic $Z_j = \frac{\hat{\beta}_j}{\sqrt{Var(\hat{\beta}_j)}} = \frac{\hat{\beta}_j}{\sqrt{\Sigma_{B_{ij}}^{-1}}}$ is used instead of the beta coefficient, and a correlation matrix $\sum_Z$ is used instead of the covariance matrix. MLC-Z can be formulated as:

$$MLC - Z = (Z^T W_o)(W_o^T \textstyle\sum_Z W_o)^{-1}(W_o^T Z),$$

where $W_o = (\sum_Z^{-1} . J)(J^T . \sum_Z^{-1} . J)^{-1}$ and *J* is defined as former. Under the null hypothesis, MLC-B and MLC-Z tests follow a chi-square distribution with *L* degree of freedom (df).

##### *Wald test*

Wald test (Wald, 1943), as a noteworthy joint regression analysis test, is defined with considering no dependency between SNPs and quantitative trait model *Y* as global null hypothesis versus at least there is a relationship between one SNP and trait *Y* as alternative hypothesis. In fact, Wald test could be considered as a special case of MLC test when each bin includes one SNP (Yoo et al., 2015). Wald test is formulated as:

$$Wald = \hat{\beta}^T \textstyle\sum_B^{-1} \hat{\beta}$$

Under the null hypothesis, Wald test follows a chi-square distribution with *m* df.

##### *LC-B and LC-Z test*

One of the special cases of the MLC test occurs when all SNPs are included in one bin; in this case, the Linear Combination (LC) test can be proposed (O'Brien, 1984; Pocock et al., 1987). LC-B and LC-Z tests are two multi-marker regression tests which LC-B is based on the beta coefficients and LC-Z is based on the corresponding *Z* statistics. LC-B and LC-Z can be formulated as follow:

$$LC - B = (\hat{\beta}^T w_s)(w_s^T \textstyle\sum_B w_s)^{-1}(w_s^T \hat{\beta})$$

Where $w_{sj} = (\sum_B^{-1}.J)_j (J^T.\sum_B^{-1}.J)^{-1}$ with $J = (1,1,...,1)^T$. Based on the definition of the MLC-Z test, the LC-Z test can be defined as below:

$$LC - B = (Z^T w_o)(w_o^T \textstyle\sum_Z w_o)^{-1}(w_o^T Z)$$

Where $w_{oj} = (\sum_Z^{-1}.J)_j (J^T.\sum_Z^{-1}.J)^{-1}$ and same as LC-B test with $J = (1,1,...,1)^T$.

Under the null hypothesis, LC-B and LC-Z tests follow a chi-square distribution with 1 df.

##### *PC80 test*

It is evident that, Principal Component analysis is one of the most famous dimension reduction methods. Principle Component analysis can create principle components of SNPs genotype as variables in multiple regression. For the simulation study based on the method of Gauderman et al. (2007), which considers 80% explained-variance criterion for determining how many *PC*s to include in the disease model, the PC80 test was selected as a gene-based test from the multiple regression analysis of a subset of principal components. Without loss of generality, *PCs* can be ordered as $PC_1$ as the maximum variance, $PC_2$ as the second-largest variance, etc. Among $PC_1, ..., PC_m$, which is ordered by the magnitude of explained variance, one can select a subset of *PC*s ($PC_1, ..., PC_S, S<m$) that explains more than 80% of the variance. The regression using S principal components is modeled as:

$$E[Y] = \beta_0^* + \beta_1^*(PC)_1 + \beta_2^*(PC)_2 + \cdots \beta_S^*(PC)_S$$

By considering $\hat{\beta}^* = (\beta_1^*, \beta_2^*, ..., \beta_S^*)^T$ as the estimated beta coefficient of principal components and their covariance matrix $\Sigma^*_B$, the PC80 is defined as:

$$PC80 = \hat{\beta}^{*T} \textstyle\sum_B^{*-1} \hat{\beta}^*$$

Under the null hypothesis, PC80 follows a mixture of chi-square distributions with S df.

By considering beta estimates $\hat{\beta} = (\hat{\beta}_1^M, \hat{\beta}_2^M, ..., \hat{\beta}_m^M)^T$ from the marginal single-SNP regression and covariance matrix $\sum_B^M$, a global statistic could be constructed by combining the marginal effects of individual SNPs.

##### *SSB and SSBw tests*

One of the sum tests' weaknesses is that power may have reduced with a small estimated global effect, $\hat{\beta}_c$, which is a linear combination of the marginal components of $\hat{\beta}_M$ with always positive coefficients, when the components $\hat{\beta}_M$ have different signs. Pan (2009) proposed a quadratic test statistic which replaces the components of $\hat{\beta}_M$ in the linear combination by their squares. The proposed test statistic defined as below:

$$SSB = \hat{\beta}_M^T \hat{\beta}_M = \sum_{j=1}^{m} \hat{\beta}_{M,j}^2$$

or its weighted version, which for each component of $\hat{\beta}_M$ based on its variance assigned a weight as follow:

$$SSBw = \hat{\beta}^T Diag(V_M)^{-1} \hat{\beta}_M = \sum_{j=1}^{m} \frac{\hat{\beta}_{M,j}^2}{v_{M,j}},$$

where$Diag(V_M) = diag(v_{M,1}, \dots, v_{M,m})$ is a diagonal matrix with elements $v_{M,j}$ as the estimated variance of $\hat{\beta}_{M,j}$ from marginal model.

Under the null hypothesis, SSB and SSBw follow a mixture of chi-square distributions with 1 df.

##### *SKAT and SKAT-O tests*

Ionita-Laza et al. (2013) proposed a quadratic supervised test, with a specific name SKAT (sequence kernel association test), which is a computationally efficient regression method to test for association between genetic variants in a region. Calculating a $p$-value directly without the requiring for permutation is one of the advantages of the SKAT test, and therefore it is very appropriate in exome and genome-wide sequencing studies because of the rapid estimation of $p$-values. SKAT is formulated as follow:

$$SKAT = Y'XWX'Y$$

where $Y$ is the $n$ by 1 vector of phenotypes, $X$ is the $n$ by $K$ matrix of genotypes and $W = diag(w_1, \dots, w_m)$ is a diagonal matrix which contains the weights of the $m$ variants and each $w_k$ is formulated as $\sqrt{w_k} = Beta(MAF_k; a_1, a_2)$ is the beta distribution density function with pre-specified parameters $a_1$ and $a_2$ evaluated at the sample minor-allele frequency (MAF) for the $k^{th}$ variant in the data.

The optimal unified test (SKAT-O) is an optimal test in the generalized SKAT family. SKAT-O is a linear combination of SKAT and a burden test (SKAT-B) (Lee et al., 2012). SKAT-O can be formulated as follow:

$$SKAT - O = \rho SKAT + (1 - \rho) SKAT - B$$

SKAT-O is a weighted average of SKAT and SKAT-B statistics.

SKAT and SKAT-O follow a mixture of chi-square distributions with 1 degree of freedom under the null hypothesis.

#### 1.2 | Quality control process of the CLSA data

The quality control process of the CLSA data involved two main components: 1) the marker-based quality control analysis consisted of four tests aimed at ensuring marker consistency across various experimental factors, including genotyping batch, participant sex, Hardy-Weinberg equilibrium (HWE), and discordance of genotyping across control replicates (with a multiple-testing corrected p-value threshold for quality control tests as 3.93×10-10); 2) the sample-based quality control was conducted to identify genotyped samples of low quality, detect related individuals, and provide a genetic-based description of ancestry. A total of 794,409 genetic markers were initially selected. After passing all four tests in the marker-based quality control (removed 35,589 markers), removing markers that were not insertions/deletions, had an MAF > 0.01, and had marker-wise missingness < 0.01, the final dataset consisted of 570,275 genetic markers. Phasing and imputation[2] process were performed utilizing the Sanger imputation Service (2018). The Haplotype Reference Consortium (HRC) reference panel, which includes 64,940 haplotypes across 40,359,612 genetic markers, is employed for this process. The genotype imputation required the marker reference alleles to match the human genome GRCh37/hg19 reference sequence. To perform the imputation, a subset of the study participants (19,669 individuals) that passed sample-based quality control and a set of 719,696 markers meeting specific criteria (passing all four marker-based QC tests, SNP-wise missingness < 0.05, and MAF > 0.0001) is used as input. Applying the bcftools +fixref plugin removed 63,043 markers, yielding a total marker count of 656,653 for the imputation process markers.

We obtained the dimension reduction results of DRLPC for chromosome 22 SNPs, in consistency with the 1000 Genomes Project data used in this study, and the number of SNPs in chromosome 22 after imputation was 524,544. A set of quality-control procedures, including marker-based and sample-based quality control, was performed on CLSA data. Marker-based quality control analysis consisted of 4 tests intended to check for consistency of markers across various experimental factors, such as genotyping batch, participant sex, Hardy-Weinberg equilibrium (HWE), and discordance of genotyping across control replicates. All tests were conducted on a subset of ancestrally homogeneous participants. The Sample-based quality control aimed to identify low-quality genotyped samples, identify related individuals, and provide a genetic-based description of ancestry. Therefore, high-quality SNPs were used to ensure no bias was introduced due to the genotype batch effect or other genotyping artifacts. Same as the 1000 Genomes Project data, first, we removed SNPs with missing values and excluded multi-allelic SNPs. Moreover, we considered MAF $\geq 0.05$ and INFO score $\geq 0.8$ (to select the well-imputed SNPs) for European ancestry. After the filtering process, the total number of remaining SNPs was 71,695. The number of individuals in the final dataset was 17,779. In the CLSA data, m dosage variables are in the range 0~2.

[2] The genotype imputation is a computational method to predict marker genotypes not directly genotyped by an assay, such as a genotyping array.

### 2 | SUPPLEMENTARY MATERIALS

#### 2.1 | Supplementary Tables

**TABLE S1** Average final variables (percentage) after the DRLPC process compared to the original data of SNPs in the gene regions of EUR, EAS, and AFR populations.

| | | EUR | | EAS | | AFR | |
|---|---|---|---|---|---|---|---|
| †CLQcut | ‡PCcut | Mean | SD | Mean | SD | Mean | SD |
| 0.5 | 0.8 | 9.3 | 14.1 | 8.3 | 13.1 | 18.5 | 35.9 |
| | 0.9 | 9.6 | 14.9 | 8.6 | 13.9 | 19.2 | 37.4 |
| 0.8 | 0.8 | 12.95 | 20.6 | 11.7 | 20.9 | 27.1 | 56.6 |
| | 0.9 | 13.2 | 21.5 | 11.8 | 21.1 | 27.6 | 57.4 |
| 0.9 | 0.8 | 14.9 | 24.3 | 13.5 | 23.4 | 30.0 | 58.5 |
| | 0.9 | 15.1 | 24.9 | 13.6 | 23.7 | 30.5 | 59.4 |
| 0.95 | 0.8 | 16.1 | 25.1 | 15.0 | 25.2 | 31.2 | 58.2 |
| | 0.9 | 16.3 | 25.9 | 15.1 | 25.8 | 31.8 | 59.4 |

†CLQcut: the threshold values for clique-based clustering algorithm CLQD (Yoo et al., 2015) to find SNP clusters.

‡PCcut: the threshold value for selecting additional PCs representing the removed SNPs at the final step of the algorithm.

**TABLE S2** Average final variables (percentage) after the DRLPC process compared to the original data of 100 SNP sets of EUR, EAS, and AFR populations.

| | | EUR | | EAS | | AFR | |
|---|---|---|---|---|---|---|---|
| CLQcut | PCcut | Mean | SD | Mean | SD | Mean | SD |
| 0.5 | 0.8 | 13.2 | 6.6 | 12.5 | 7.1 | 22.1 | 9.4 |
| | 0.9 | 13.8 | 6.9 | 13.0 | 7.2 | 23.0 | 9.5 |
| 0.8 | 0.8 | 19.1 | 9.3 | 18.2 | 9.8 | 32.8 | 13.2 |
| | 0.9 | 19.3 | 9.4 | 18.4 | 9.9 | 33.2 | 13.3 |
| 0.9 | 0.8 | 22.5 | 10.7 | 21.7 | 11.2 | 37.7 | 14.4 |
| | 0.9 | 22.6 | 10.8 | 21.9 | 11.3 | 38.0 | 14.4 |
| 0.95 | 0.8 | 25.0 | 11.4 | 24.6 | 12.2 | 40.7 | 14.7 |
| | 0.9 | 25.1 | 11.5 | 24.8 | 12.3 | 40.9 | 14.8 |

**TABLE S3** Average final variables (percentage) after the DRLPC process compared to the original data of SNPs in the gene regions of European ancestry for chromosome 22 CLSA data.

| CLQcut | PCcut | Mean | SD |
|---|---|---|---|
| 0.5 | 0.8 | 7.85 | 12.2 |
| | 0.9 | 8.1 | 12.8 |
| 0.8 | 0.8 | 11.1 | 18.2 |
| | 0.9 | 11.3 | 19.0 |
| 0.9 | 0.8 | 12.7 | 20.7 |
| | 0.9 | 12.9 | 21.6 |
| 0.95 | 0.8 | 14.1 | 24.9 |
| | 0.9 | 14.3 | 25.5 |

**TABLE S4** Average final variables (percentage) after the DRLPC process compared to the original data of 100 SNP sets of European ancestry for chromosome 22 CLSA data.

| CLQcut | PCcut | Mean | SD |
|---|---|---|---|
| 0.5 | 0.8 | 11.3 | 4.6 |
| | 0.9 | 11.5 | 4.7 |
| 0.8 | 0.8 | 16.3 | 7.1 |
| | 0.9 | 16.5 | 7.3 |
| 0.9 | 0.8 | 19.0 | 8.2 |
| | 0.9 | 19.2 | 8.3 |
| 0.95 | 0.8 | 21.1 | 8.8 |
| | 0.9 | 21.3 | 8.9 |

**TABLE S5** The average and standard deviation of the highest VIF values after each DRLPC step obtained over gene regions in chromosome 22 using CLSA data of European ancestry.

| | | Step 1 | | Step 2 & 3 | | Step 4 | | Step5 | |
|---|---|---|---|---|---|---|---|---|---|
| CLQcut | PCcut | Average | SD | Average | SD | Average | SD | Average | SD |
| 0.5 | 0.8 | 764564470658 | 1.9e+13 | 17.6 | 34.8 | 5.4 | 5.7 | 5.9 | 6.6 |
| | 0.9 | 764564470658 | 1.9e+13 | 17.6 | 34.8 | 5.4 | 5.7 | 6.8 | 8.8 |
| 0.8 | 0.8 | 764564470658 | 1.9e+13 | 142.1 | 246.9 | 8.2 | 6.6 | 8.5 | 7.3 |
| | 0.9 | 764564470658 | 1.9e+13 | 142.1 | 246.9 | 8.2 | 6.6 | 9.1 | 10.5 |
| 0.9 | 0.8 | 764564470658 | 1.9e+13 | 249.9 | 402.6 | 9.4 | 6.7 | 9.5 | 7.0 |
| | 0.9 | 764564470658 | 1.9e+13 | 249.9 | 402.6 | 9.4 | 6.7 | 10.1 | 8.4 |
| 0.95 | 0.8 | 764564470658 | 1.9e+13 | 362.1 | 567.8 | 12.2 | 26.2 | 12.4 | 26.2 |
| | 0.9 | 764564470658 | 1.9e+13 | 362.1 | 567.8 | 12.2 | 26.2 | 13.5 | 27.7 |

**TABLE S6** The average and standard deviation of variable proportion values after each DRLPC step obtained over gene regions in chromosome 22 using 1000 Genomes Project data of three super-populations.

| | CLQcut | PCcut | Step 1 | | Step2 & 3 | | Step 4 | | Step5 | |
|---|---|---|---|---|---|---|---|---|---|---|
| | | | Removed variables | | Remain variable | | Removed variables | | New variables | |
| | | | Average | SD | Average | SD | Average | SD | Average | SD |
| EUR | 0.5 | 0.8 | [†¶]-34.3 | -82.4 | 9.3 | 15.2 | -0.4 | -1.6 | [‡]+0.3 | +0.6 |
| | | 0.9 | -34.3 | -82.4 | 9.3 | 15.2 | -0.4 | -1.6 | +0.6 | +1.3 |
| | 0.8 | 0.8 | -34.3 | -82.4 | 15.6 | 29.7 | -2.8 | -10.4 | +0.1 | +0.6 |
| | | 0.9 | -34.3 | -82.4 | 15.6 | 29.7 | -2.8 | -10.4 | +0.3 | +1.6 |
| | 0.9 | 0.8 | -34.3 | -82.4 | 19.5 | 38.7 | -4.7 | -15.4 | +0.1 | +0.4 |
| | | 0.9 | -34.3 | -82.4 | 19.5 | 38.7 | -4.7 | -15.4 | +0.8 | +16.1 |
| | 0.95 | 0.8 | -34.3 | -82.4 | 23.1 | 47.4 | -7.1 | -23.5 | +0.1 | +0.4 |
| | | 0.9 | -34.3 | -82.4 | 23.1 | 47.4 | -7.1 | -23.5 | +0.3 | +1.5 |
| EAS | 0.5 | 0.8 | -30.9 | -71.0 | 8.3 | 13.8 | -0.3 | -1.2 | +0.3 | +0.6 |
| | | 0.9 | -30.9 | -71.0 | 8.3 | 13.8 | -0.3 | -1.2 | +0.6 | +1.3 |
| | 0.8 | 0.8 | -30.9 | -71.0 | 13.6 | 26.1 | -2.0 | -5.7 | +0.1 | +0.4 |
| | | 0.9 | -30.9 | -71.0 | 13.6 | 26.1 | -2.0 | -5.7 | +0.2 | +0.8 |
| | 0.9 | 0.8 | -30.9 | -71.0 | 17.1 | 34.1 | -3.6 | -11.4 | +0.0 | +0.2 |
| | | 0.9 | -30.9 | -71.0 | 17.1 | 34.1 | -3.6 | -11.4 | +0.2 | +0.7 |
| | 0.95 | 0.8 | -30.9 | -71.0 | 20.5 | 42.1 | -5.6 | -17.6 | +0.6 | +15.2 |
| | | 0.9 | -30.9 | -71.0 | 20.5 | 42.1 | -5.6 | -17.6 | +0.8 | +16.2 |
| AFR | 0.5 | 0.8 | -27.6 | -66.8 | 19.1 | 39.1 | -1.2 | -4.8 | +0.5 | +1.1 |
| | | 0.9 | -27.6 | -66.8 | 19.1 | 39.1 | -1.2 | -4.8 | +1.3 | +2.9 |
| | 0.8 | 0.8 | -27.6 | -66.8 | 33.6 | 74.2 | -6.7 | -19.7 | +0.2 | +0.8 |
| | | 0.9 | -27.6 | -66.8 | 33.6 | 74.2 | -6.7 | -19.7 | +0.7 | +2.6 |
| | 0.9 | 0.8 | -27.6 | -66.8 | 41.6 | 93.6 | -11.8 | -37.1 | +0.1 | +0.8 |
| | | 0.9 | -27.6 | -66.8 | 41.6 | 93.6 | -11.8 | -37.1 | +0.7 | +2.6 |
| | 0.95 | 0.8 | -27.6 | -66.8 | 47.9 | 108.0 | -16.9 | -52.2 | +0.2 | +1.0 |
| | | 0.9 | -27.6 | -66.8 | 47.9 | 108.0 | -16.9 | -52.2 | +0.8 | +3.2 |

[†]Sings (-) indicate that the average and standard deviation of removed variables; [‡]Signs (+) indicate the average and standard deviation of added variables.

**TABLE S7** The average and standard deviation of variable proportion values after each DRLPC step obtained over gene regions in chromosome 22 using the CLSA data of European ancestry.

| CLQcut | PCcut | Step 1 Removed variables | | Steps 2 & 3 Remain variable | | Step 4 Removed variables | | Step5 New variables | |
|---|---|---|---|---|---|---|---|---|---|
| | | Average | SD | Average | SD | Average | SD | Average | SD |
| 0.5 | 0.8 | -1.01 | -3.8 | 8.3 | 13.5 | -0.5 | -1.7 | +0.1 | +0.3 |
| | 0.9 | -1.01 | -3.8 | 8.3 | 13.5 | -0.5 | -1.7 | +0.3 | +0.9 |
| 0.8 | 0.8 | -1.01 | -3.8 | 14.3 | 27.5 | -3.3 | -10.5 | +0.1 | +0.8 |
| | 0.9 | -1.01 | -3.8 | 14.3 | 27.5 | -3.3 | -10.5 | +0.3 | +1.7 |
| 0.9 | 0.8 | -1.01 | -3.8 | 18.3 | 36.9 | -5.6 | -17.5 | +0.1 | +0.7 |
| | 0.9 | -1.01 | -3.8 | 18.3 | 36.9 | -5.6 | -17.5 | +0.3 | +1.7 |
| 0.95 | 0.8 | -1.01 | -3.8 | 22.0 | 46.2 | -8.0 | -22.5 | +0.1 | +0.4 |
| | 0.9 | -1.01 | -3.8 | 22.0 | 46.2 | -8.0 | -22.5 | +0.3 | +1.3 |

**TABLE S8** Empirical type I error of gene-based statistics (N=10,000 replicate) at the nominal level α=0.05, averaged over ten genes, using original data and two DRLPC processed data for three populations.

| | | †Original data | | | CLQcut 0.5 | | | CLQcut 0.8 | | |
|---|---|---|---|---|---|---|---|---|---|---|
| Population | ‡Statistics | Average | SD | Average of df | Average | SD | Average of df | Average | SD | Average of df |
| EUR | LC-Z | 0.050 | 0.003 | 1.0 | 0.051 | 0.002 | 1.0 | 0.051 | 0.003 | 1.0 |
| | MLC-Z5 | 0.053 | 0.003 | 8.3 | 0.052 | 0.003 | 6.9 | 0.052 | 0.003 | 7.4 |
| | MLC-Z8 | 0.055 | 0.004 | 13.3 | 0.052 | 0.003 | 8.1 | 0.053 | 0.003 | 10.8 |
| | SSB | 0.052 | 0.003 | - | 0.054 | 0.003 | - | 0.053 | 0.003 | - |
| | SSBw | 0.054 | 0.003 | - | 0.056 | 0.003 | - | 0.055 | 0.003 | - |
| EAS | LC-Z | 0.050 | 0.003 | 1.0 | 0.051 | 0.002 | 1.0 | 0.051 | 0.003 | 1.0 |
| | MLC-Z5 | 0.053 | 0.003 | 7.4 | 0.052 | 0.003 | 6.1 | 0.052 | 0.003 | 6.7 |
| | MLC-Z8 | 0.055 | 0.004 | 11.8 | 0.052 | 0.003 | 7.2 | 0.053 | 0.003 | 9.6 |
| | SSB | 0.052 | 0.003 | - | 0.054 | 0.003 | - | 0.053 | 0.003 | - |
| | SSBw | 0.054 | 0.003 | - | 0.056 | 0.003 | - | 0.056 | 0.003 | - |
| AFR | LC-Z | 0.050 | 0.002 | 1.0 | 0.051 | 0.002 | 1.0 | 0.050 | 0.002 | 1.0 |
| | MLC-Z5 | 0.054 | 0.003 | 16.4 | 0.053 | 0.003 | 12.8 | 0.053 | 0.003 | 14.8 |
| | MLC-Z8 | 0.058 | 0.005 | 27.6 | 0.054 | 0.003 | 16.2 | 0.055 | 0.004 | 22.7 |
| | SSB | 0.051 | 0.003 | - | 0.052 | 0.003 | - | 0.051 | 0.003 | - |
| | SSBw | 0.054 | 0.002 | - | 0.054 | 0.002 | - | 0.054 | 0.002 | - |

†Used data: Original data; CLQcut 0.5: DRLPC processed data suing CLQ threshold value 0.5; CLQcut 0.8: DRLPC processed data suing CLQ threshold value 0.8.

‡List of test statistics: LC-Z: linear combination test using Z statistics (O'Brien et al, 1984; Pocock et al, 1987); MLC test: Multi Linear combination test (Yoo et al, 2016) for MLC-Z5: MLC tests using Z statistics by considering CLQcut equal to 0.5, MLC-Z8: MLC tests using Z statistics by considering CLQcut equal to 0.8; SSB: sum of squared marginal beta coefficients (Pan et al, 2009); SSBw: sum of squared marginal beta coefficients with inverse variance weights (Pan et al, 2009).

**TABLE S9** Empirical type I error of gene-based statistics (N=10,000 replicate) at the nominal level α=0.01, averaged over ten genes, using original data and two DRLPC processed data for three populations.

| | | Original data | | | CLQcut 0.5 | | | CLQcut 0.8 | | |
|---|---|---|---|---|---|---|---|---|---|---|
| Population | ‡Statistics | Average | SD | Average of df | Average | SD | Average of df | Average | SD | Average of df |
| EUR | Wald | 0.021 | 0.005 | 32.5 | 0.011 | 0.001 | 8.4 | 0.012 | 0.001 | 11.4 |
| | PC80 | 0.011 | 0.001 | 4.0 | 0.011 | 0.001 | 4.6 | 0.011 | 0.001 | 5.0 |
| | MLC-B5 | 0.011 | 0.001 | 8.3 | 0.011 | 0.001 | 6.9 | 0.011 | 0.001 | 7.5 |
| | MLC-Z5 | 0.011 | 0.001 | 8.3 | 0.011 | 0.001 | 6.9 | 0.011 | 0.001 | 7.5 |
| | MLC-B8 | 0.012 | 0.002 | 13.3 | 0.011 | 0.001 | 8.1 | 0.012 | 0.002 | 10.8 |
| | MLC-Z8 | 0.012 | 0.002 | 13.3 | 0.011 | 0.001 | 8.1 | 0.011 | 0.001 | 10.8 |
| | SSB | 0.010 | 0.001 | - | 0.011 | 0.001 | - | 0.011 | 0.001 | - |
| | SSBw | 0.011 | 0.001 | - | 0.012 | 0.001 | - | 0.012 | 0.001 | - |
| | SKAT | 0.010 | 0.001 | - | 0.010 | 0.001 | - | 0.010 | 0.001 | - |
| | SKAT-O | 0.011 | 0.001 | - | 0.011 | 0.001 | - | 0.011 | 0.001 | - |
| | LC-B | 0.010 | 0.001 | 1.0 | 0.010 | 0.001 | 1.0 | 0.010 | 0.001 | 1.0 |
| | LC-Z | 0.010 | 0.001 | 1.0 | 0.010 | 0.001 | 1.0 | 0.010 | 0.001 | 1.0 |
| EAS | Wald | 0.019 | 0.006 | 30.8 | 0.011 | 0.001 | 7.6 | 0.012 | 0.001 | 10.3 |
| | PC80 | 0.011 | 0.001 | 3.6 | 0.011 | 0.001 | 4.4 | 0.011 | 0.001 | 4.7 |
| | MLC-B5 | 0.011 | 0.001 | 7.4 | 0.011 | 0.001 | 6.1 | 0.011 | 0.001 | 6.7 |
| | MLC-Z5 | 0.011 | 0.001 | 7.4 | 0.011 | 0.001 | 6.1 | 0.011 | 0.001 | 6.7 |
| | MLC-B8 | 0.012 | 0.002 | 11.8 | 0.011 | 0.001 | 7.3 | 0.011 | 0.001 | 9.7 |
| | MLC-Z8 | 0.012 | 0.002 | 11.8 | 0.011 | 0.001 | 7.3 | 0.011 | 0.001 | 9.7 |
| | SSB | 0.011 | 0.001 | - | 0.011 | 0.001 | - | 0.011 | 0.001 | - |
| | SSBw | 0.011 | 0.001 | - | 0.012 | 0.001 | - | 0.012 | 0.001 | - |
| | SKAT | 0.010 | 0.001 | - | 0.010 | 0.001 | - | 0.010 | 0.001 | - |
| | SKAT-O | 0.011 | 0.001 | - | 0.011 | 0.001 | - | 0.011 | 0.001 | - |
| | LC-B | 0.010 | 0.001 | 1.0 | 0.010 | 0.001 | 1.0 | 0.010 | 0.001 | 1.0 |
| | LC-Z | 0.010 | 0.001 | 1.0 | 0.010 | 0.001 | 1.0 | 0.010 | 0.001 | 1.0 |
| AFR | Wald | 0.024 | 0.008 | 58.8 | 0.012 | 0.001 | 16.5 | 0.012 | 0.002 | 22.2 |
| | PC80 | 0.011 | 0.001 | 7.0 | 0.011 | 0.001 | 7.5 | 0.011 | 0.001 | 8.1 |
| | MLC-B5 | 0.012 | 0.001 | 16.4 | 0.011 | 0.001 | 12.8 | 0.011 | 0.001 | 14.3 |
| | MLC-Z5 | 0.012 | 0.002 | 16.4 | 0.011 | 0.001 | 12.8 | 0.011 | 0.001 | 14.3 |

| | | | | | | | | | | |
|---|---|---|---|---|---|---|---|---|---|---|
| | MLC-B8 | 0.013 | 0.002 | 27.6 | 0.012 | 0.001 | 16.2 | 0.012 | 0.002 | 21.4 |
| | MLC-Z8 | 0.013 | 0.003 | 27.6 | 0.012 | 0.001 | 16.2 | 0.012 | 0.002 | 21.4 |
| | PC80 | 0.011 | 0.001 | 7.0 | 0.011 | 0.001 | 7.5 | 0.011 | 0.001 | 8.1 |
| | SSB | 0.010 | 0.001 | - | 0.010 | 0.001 | - | 0.010 | 0.001 | - |
| | SSBw | 0.011 | 0.001 | - | 0.011 | 0.001 | - | 0.011 | 0.001 | - |
| | SKAT | 0.010 | 0.001 | - | 0.010 | 0.001 | - | 0.010 | 0.001 | - |
| | SKAT-O | 0.011 | 0.001 | - | 0.011 | 0.001 | - | 0.011 | 0.001 | - |
| | LC-B | 0.010 | 0.001 | 1.0 | 0.010 | 0.001 | 1.0 | 0.010 | 0.001 | 1.0 |
| | LC-Z | 0.010 | 0.001 | 1.0 | 0.010 | 0.001 | 1.0 | 0.010 | 0.001 | 1.0 |

‡List of test statistics: Wald: generalized Wald test (Wald, 1943); LC-B: linear combination test using beta coefficients (O'Brien et al, 1984; Pocock et al, 1987; MLC test: Multi Linear combination test (Yoo et al, 2014) for ; MLC-B5: MLC tests using beta coefficient by considering CLQcut equal to 0.5, MLC-B8 MLC tests using beta coefficient by considering CLQcut equal to 0.8; PC80: global test on regression using the minimum number of principal components capturing 80% of variance (Gauderman et al, 2007); SKAT: sequence kernel associated tests for the common variants (Ionita-Laza et al, 2013); SKAT-O: a linear combination of SKAT and burden test with optimized mixing proportion (Lee et al, 2012).

**TABLE S10** Empirical type I error of gene-based tests using original data and two DRLPC processed data (CLQcut=0.5 and 0.8) for 100 genes stratified by gene size (number of SNPs), 1causal model, EUR population.

| | From 1 to 100 ($N_1$= 79) | | | From 101 to 200 ($N_2$= 14) | | | From 201 to 500 ($N_3$= 7) | | |
|---|---|---|---|---|---|---|---|---|---|
| Tests | †Org | ‡CLQ5 | §CLQ8 | Org | CLQ5 | CLQ8 | Org | CLQ5 | CLQ8 |
| Wald | 0.067 | 0.052 | 0.053 | 0.083 | 0.053 | 0.054 | 0.090 | 0.054 | 0.055 |
| PC80 | 0.051 | 0.052 | 0.052 | 0.051 | 0.051 | 0.052 | 0.051 | 0.052 | 0.052 |
| MLC-B5 | 0.052 | 0.052 | 0.052 | 0.054 | 0.052 | 0.052 | 0.057 | 0.053 | 0.054 |
| MLC-Z5 | 0.053 | 0.052 | 0.052 | 0.054 | 0.052 | 0.054 | 0.056 | 0.052 | 0.054 |
| MLC-B8 | 0.054 | 0.054 | 0.053 | 0.057 | 0.053 | 0.054 | 0.060 | 0.054 | 0.057 |
| MLC-Z8 | 0.054 | 0.052 | 0.053 | 0.057 | 0.053 | 0.054 | 0.060 | 0.053 | 0.058 |
| SSBw | 0.053 | 0.056 | 0.830 | 0.054 | 0.056 | 0.056 | 0.054 | 0.057 | 0.057 |
| SKAT | 0.049 | 0.049 | 0.049 | 0.050 | 0.050 | 0.050 | 0.050 | 0.050 | 0.051 |
| SKAT-O | 0.052 | 0.052 | 0.052 | 0.053 | 0.052 | 0.053 | 0.052 | 0.053 | 0.053 |

†Org: original data; ‡CLQ5; CLQcut value is 0.5; CLQ8; §CLQcut value is 0.8.

**TABLE S11** Empirical type I error of gene-based tests using original data and two DRLPC processed data (CLQcut=0.5 and 0.8) for 100 genes stratified by gene size (number of SNPs), 1causal model, EAS population.

| | From 1 to 100 ($N_1$=82) | | | From 101 to 200 ($N_2$=12 ) | | | From 201 to 500 ($N_3$=6 ) | | |
|---|---|---|---|---|---|---|---|---|---|
| Tests | Org | CLQ5 | CLQ8 | Org | CLQ5 | CLQ8 | Org | CLQ5 | CLQ8 |
| Wald | 0.066 | 0.053 | 0.053 | 0.081 | 0.053 | 0.055 | 0.088 | 0.053 | 0.053 |
| PC80 | 0.051 | 0.052 | 0.052 | 0.052 | 0.052 | 0.053 | 0.050 | 0.051 | 0.051 |
| MLC-B5 | 0.053 | 0.052 | 0.052 | 0.054 | 0.052 | 0.053 | 0.054 | 0.051 | 0.052 |
| MLC-Z5 | 0.052 | 0.053 | 0.052 | 0.055 | 0.052 | 0.053 | 0.056 | 0.053 | 0.052 |
| MLC-B8 | 0.053 | 0.052 | 0.053 | 0.057 | 0.053 | 0.055 | 0.057 | 0.052 | 0.053 |
| MLC-Z8 | 0.054 | 0.052 | 0.052 | 0.058 | 0.053 | 0.055 | 0.085 | 0.092 | 0.089 |
| SSBw | 0.054 | 0.056 | 0.055 | 0.055 | 0.055 | 0.056 | 0.061 | 0.054 | 0.056 |
| SKAT | 0.049 | 0.049 | 0.049 | 0.050 | 0.050 | 0.050 | 0.050 | 0.048 | 0.048 |
| SKAT-O | 0.052 | 0.052 | 0.052 | 0.051 | 0.052 | 0.053 | 0.053 | 0.050 | 0.050 |

**TABLE S12** Empirical type I error of gene-based tests using original data and two DRLPC processed data (CLQcut=0.5 and 0.8) for 100 genes stratified by gene size (number of SNPs), 1causal model, AFR population.

| | From 1 to 100 ($N_1$=68) | | | From 101 to 200 ($N_2$=18 ) | | | From 201 to 500 ($N_3$=14 ) | | |
|---|---|---|---|---|---|---|---|---|---|
| Tests | Org | CLQ5 | CLQ8 | Org | CLQ5 | CLQ8 | Org | CLQ5 | CLQ8 |
| Wald | 0.067 | 0.053 | 0.054 | 0.09 | 0.055 | 0.058 | 0.100 | 0.056 | 0.059 |
| PC80 | 0.051 | 0.052 | 0.052 | 0.052 | 0.052 | 0.053 | 0.052 | 0.053 | 0.053 |
| MLC-B5 | 0.053 | 0.052 | 0.053 | 0.056 | 0.054 | 0.055 | 0.057 | 0.055 | 0.057 |
| MLC-Z5 | 0.053 | 0.053 | 0.053 | 0.056 | 0.054 | 0.054 | 0.057 | 0.056 | 0.056 |
| MLC-B8 | 0.055 | 0.53 | 0.051 | 0.06 | 0.055 | 0.057 | 0.063 | 0.056 | 0.059 |
| MLC-Z8 | 0.055 | 0.053 | 0.054 | 0.055 | 0.055 | 0.056 | 0.067 | 0.056 | 0.059 |
| SSBw | 0.054 | 0.054 | 0.054 | 0.053 | 0.054 | 0.054 | 0.055 | 0.055 | 0.055 |
| SKAT | 0.050 | 0.050 | 0.050 | 0.050 | 0.050 | 0.050 | 0.051 | 0.050 | 0.05 |
| SKAT-O | 0.052 | 0.052 | 0.052 | 0.052 | 0.052 | 0.052 | 0.052 | 0.053 | 0.053 |

**TABLE S13** Empirical power (percentage) of gene-based statistics (N=1000 replicate) at the 0.05 level and average of degree of freedom for three populations, 1causal model, averaged over 100 genes.

| Population | Statistics | Original data | | | CLQcut 0.5 | | | CLQcut 0.8 | | |
|---|---|---|---|---|---|---|---|---|---|---|
| | | Power | | df | Power | | df | Power | | df |
| | | Average | SD | Average | Average | SD | Average | Average | SD | Average |
| EUR | MLC-Z5 | 80.2 | 8.9 | 8.9 | 80.2 | 15.1 | 7.3 | 80.7 | 9.5 | 8.1 |
| | MLC-Z8 | 79.5 | 17.7 | 7.8 | 81.3 | 11.7 | 8.7 | 78.9 | 8.5 | 11.9 |
| | SSB | 83.1 | 15.2 | - | 34.2 | 24.6 | - | 55.0 | 28.9 | - |
| | SSBw | 78.4 | 19.0 | - | 84.4 | 10.1 | - | 84.35 | 9.4 | - |
| | LC-B | 49.8 | 33.6 | 1.0 | 45.9 | 32.1 | 1.0 | 43.5 | 31.6 | 1.0 |
| | LC-Z | 49.8 | 34.9 | 1.0 | 37.0 | 30.1 | 1.0 | 39.6 | 30.1 | 1.0 |
| EAS | MLC-Z5 | 80.9 | 11.8 | 7.8 | 81.1 | 11.9 | 6.4 | 79.8 | 12.8 | 7.1 |
| | MLC-Z8 | 78.1 | 7.7 | 12.7 | 82.7 | 8.6 | 7.6 | 80.2 | 7.9 | 10.4 |
| | SSB | 82.8 | 14.4 | - | 37.4 | 26.6 | - | 56.1 | 29.1 | - |
| | SSBw | 80.9 | 17.0 | - | 84.9 | 10.6 | - | 85.1 | 9.1 | - |
| | LC-B | 48.6 | 34.1 | 1.0 | 52.0 | 31.2 | 1.0 | 47.5 | 31.9 | 1.0 |
| | LC-Z | 47.8 | 35.0 | 1.0 | 37.0 | 30.0 | 1.0 | 39.6 | 29.8 | 1.0 |
| AFR | MLC-Z5 | 82.5 | 9.0 | 18.6 | 82.2 | 12.1 | 14.0 | 82.8 | 9.9 | 16.2 |
| | MLC-Z8 | 81.3 | 11.3 | 21.3 | 82.9 | 8.6 | 17.8 | 80.0 | 8.1 | 25.4 |
| | SSB | 83.0 | 14.6 | - | 36.7 | 23.9 | - | 66.6 | 28.3 | - |
| | SSBw | 81.2 | 16.0 | - | 86.2 | 9.8 | - | 87.5 | 8.8 | - |
| | LC-B | 33.3 | 29.2 | 1.0 | 33.2 | 26.2 | 1.0 | 31.9 | 27.1 | 1.0 |
| | LC-Z | 33.0 | 28.8 | 1.0 | 29.1 | 26.4 | 1.0 | 31.5 | 25.3 | 1.0 |

**TABLE S14** Empirical power (percentage) of gene-based statistics (N=1000 replicate) at the 0.05 level and average of degree of freedom for three populations, 2causal model, averaged over 100 genes.

| Population | Statistics | Original data | | | CLQcut 0.5 | | | CLQcut 0.8 | | |
|---|---|---|---|---|---|---|---|---|---|---|
| | | Average power | SD | Average df | Average power | SD | Average df | Average power | SD | Average df |
| EUR | Wald | 60.9 | 4.8 | 35.7 | 82.0 | 9.6 | 9.0 | 79.6 | 7.7 | 12.7 |
| | PC80 | 84.7 | 13.4 | 4.2 | 80.3 | 14.5 | 4.8 | 80.9 | 14.5 | 5.3 |
| | MLC-B5 | 80.9 | 12.5 | 9.0 | 82.5 | 11.9 | 7.3 | 80.9 | 13.8 | 8.1 |
| | MLC-B8 | 77.8 | 12.5 | 9.0 | 82.4 | 11.9 | 7.3 | 80.4 | 13.8 | 8.1 |
| | MLC-Z5 | 80.9 | 12.2 | 8.9 | 80.1 | 14.7 | 8.7 | 80.7 | 80.7 | 12.9 |
| | MLC-Z8 | 82.0 | 12.9 | 7.8 | 82.3 | 9.5 | 8.7 | 80.3 | 7.8 | 11.9 |
| | SSB | 83.3 | 15.9 | - | 40.4 | 21.9 | - | 61.9 | 22.5 | - |
| | SSBw | 84.1 | 15.0 | - | 85.2 | 12.3 | - | 85.3 | 11.6 | - |
| | SKAT | 83.2 | 15.9 | - | 81.7 | 17.5 | - | 76.4 | 21.8 | - |
| | SKAT-O | 81.9 | 16.6 | - | 80.9 | 17.6 | - | 76.9 | 21.5 | - |
| | LC-B | 57.2 | 34.2 | 1.0 | 59.3 | 32.3 | 1.0 | 59.9 | 31.4 | 1.0 |
| | LC-Z | 56.9 | 34.7 | 1.0 | 47.6 | 33.0 | 1.0 | 55.2 | 31.1 | 1.0 |
| EAS | Wald | 60.3 | 5.5 | 33.3 | 81.5 | 9.4 | 7.9 | 79.3 | 8.4 | 11.2 |
| | PC80 | 84.6 | 12.1 | 3.7 | 78.9 | 14.1 | 4.6 | 79.8 | 13.4 | 4.9 |
| | MLC-B5 | 79.1 | 15.2 | 7.9 | 82.2 | 10.5 | 6.4 | 79.2 | 14.3 | 7.1 |
| | MLC-Z5 | 79.2 | 14.7 | 7.9 | 81.1 | 11.1 | 6.4 | 77.2 | 16.1 | 7.1 |
| | MLC-B8 | 77.8 | 8.4 | 12.6 | 81.9 | 10 | 7.6 | 79.9 | 8.6 | 10.4 |
| | MLC-Z8 | 77.7 | 8.4 | 12.6 | 81.8 | 10 | 7.6 | 79.5 | 8.6 | 10.4 |
| | SSB | 80.9 | 17.5 | - | 39.1 | 22.1 | - | 59.4 | 23.9 | - |
| | SSBw | 84.3 | 15.7 | - | 82.9 | 14 | - | 83.9 | 12.7 | - |
| | SKAT | 83.4 | 16.4 | - | 78.3 | 19.8 | - | 71.6 | 21.2 | - |
| | SKAT-O | 82.8 | 16.2 | - | 78.4 | 19.4 | - | 72.8 | 21.4 | - |
| | LC-B | 62.1 | 32.1 | 1.0 | 65.6 | 29.8 | 1.0 | 58.9 | 32.9 | 1.0 |
| | LC-Z | 62.4 | 31.5 | 1.0 | 52.0 | 31.1 | 1.0 | 51.7 | 31.8 | 1.0 |
| AFR | Wald | 60.8 | 3.9 | 67.8 | 82.7 | 9.2 | 18.1 | 79.5 | 7.7 | 26.4 |
| | PC80 | 82.3 | 8.8 | 7.4 | 81.7 | 13.0 | 7.9 | 80.8 | 10.1 | 8.8 |
| | MLC-B5 | 83.0 | 7.8 | 18.6 | 83.9 | 9.7 | 14.0 | 83.1 | 9.3 | 16.2 |
| | MLC-Z5 | 82.2 | 8.7 | 18.6 | 82.7 | 11.0 | 14.0 | 82.3 | 9.4 | 16.2 |

| | | | | | | | | | |
|---|---|---|---|---|---|---|---|---|---|
| MLC-B8 | 77.1 | 6.8 | 31.5 | 82.7 | 9.3 | 17.8 | 80.2 | 7.9 | 25.4 |
| MLC-Z8 | 81.1 | 10.8 | 19.7 | 82.7 | 9.4 | 17.8 | 80.1 | 7.9 | 25.4 |
| SSB | 81.2 | 15.5 | - | 43.7 | 24.1 | - | 67.7 | 23.0 | - |
| SSBw | 83.5 | 15.2 | - | 86.7 | 11.5 | - | 87.3 | 10.1 | - |
| SKAT | 82.1 | 16.5 | - | 79.6 | 17.1 | - | 74.2 | 21.4 | - |
| SKAT-O | 79.4 | 17.5 | - | 77.9 | 17.3 | - | 72.3 | 21.4 | - |
| LC-B | 43.1 | 32.1 | 1.0 | 44.2 | 30.7 | 1.0 | 38.6 | 29.3 | 1.0 |
| LC-Z | 41.8 | 32.4 | 1.0 | 36.9 | 28.6 | 1.0 | 37.6 | 28.7 | 1.0 |

**TABLE S15** Empirical power of gene-based tests using original data and two DRLPC processed data (CLQcut=0.5 and 0.8) for 100 genes stratified by gene size (number of SNPs), 1causal model, EUR population.

| | From 1 to 100 ($N_1$= 79) | | | From 101 to 200 ($N_2$= 14) | | | From 201 to 500 ($N_3$= 7) | | |
|---|---|---|---|---|---|---|---|---|---|
| Tests | Org | CLQ5 | CLQ8 | Org | CLQ5 | CLQ8 | Org | CLQ5 | CLQ8 |
| Wald | 0.6 | 0.8 | 0.8 | 0.6 | 0.9 | 0.87 | 0.6 | 0.9 | 0.9 |
| PC80 | 0.8 | 0.8 | 0.8 | 0.8 | 0.9 | 0.89 | 0.9 | 1.0 | 0.9 |
| MLC-B5 | 0.8 | 0.8 | 0.8 | 0.9 | 0.9 | 0.91 | 0.9 | 0.9 | 0.9 |
| MLC-Z5 | 0.8 | 0.8 | 0.8 | 0.9 | 0.9 | 0.90 | 0.9 | 0.9 | 0.9 |
| MLC-B8 | 0.8 | 0.8 | 0.8 | 0.8 | 0.9 | 0.88 | 0.8 | 0.9 | 0.9 |
| MLC-Z8 | 0.8 | 0.8 | 0.8 | 0.8 | 0.9 | 0.88 | 0.9 | 0.9 | 0.9 |
| SSBw | 0.8 | 0.8 | 0.8 | 0.7 | 0.9 | 0.89 | 0.8 | 0.9 | 0.9 |
| SKAT | 0.8 | 0.8 | 0.7 | 0.7 | 0.8 | 0.83 | 0.8 | 0.8 | 0.8 |
| SKAT-O | 0.8 | 0.8 | 0.7 | 0.7 | 0.8 | 0.82 | 0.7 | 0.7 | 0.8 |

**TABLE S16** Empirical power of gene-based tests using original data and two DRLPC processed data (CLQcut=0.5 and 0.8) for 100 genes stratified by gene size (number of SNPs), 2causal model, EUR population.

| | From 1 to 100 ($N_1$= 79) | | | From 101 to 200 ($N_2$= 14) | | | From 201 to 500 ($N_3$= 7) | | |
|---|---|---|---|---|---|---|---|---|---|
| Tests | Org | CLQ5 | CLQ8 | Org | CLQ5 | CLQ8 | Org | CLQ5 | CLQ8 |
| Wald | 0.6 | 0.8 | 0.8 | 0.6 | 0.9 | 0.9 | 0.6 | 0.9 | 0.9 |
| PC80 | 0.8 | 0.8 | 0.8 | 0.9 | 0.9 | 0.9 | 1.0 | 0.9 | 1.0 |
| MLC-B5 | 0.8 | 0.8 | 0.8 | 0.9 | 0.9 | 0.9 | 0.9 | 0.9 | 0.9 |
| MLC-Z5 | 0.8 | 0.8 | 0.8 | 0.9 | 0.9 | 0.9 | 0.9 | 0.9 | 0.9 |
| MLC-B8 | 0.7 | 0.8 | 0.8 | 0.9 | 0.9 | 0.9 | 0.8 | 0.9 | 0.9 |
| MLC-Z8 | 0.8 | 0.8 | 0.8 | 0.8 | 0.9 | 0.9 | 0.9 | 0.9 | 0.9 |
| SSBw | 0.8 | 0.8 | 0.8 | 0.9 | 0.9 | 0.9 | 0.9 | 1.0 | 1.0 |
| SKAT | 0.8 | 0.8 | 0.8 | 0.9 | 0.9 | 0.8 | 0.9 | 0.9 | 0.9 |
| SKAT-O | 0.8 | 0.8 | 0.8 | 0.8 | 0.9 | 0.8 | 0.9 | 0.9 | 0.9 |

**TABLE S17** Empirical power of gene-based tests using original data and two DRLPC processed data (CLQcut=0.5 and 0.8) for 100 genes stratified by gene size (number of SNPs), 1causal model, EAS population.

| | From 1 to 100 ($N_1$=82) | | | From 101 to 200 ($N_2$=12 ) | | | From 201 to 500 ($N_3$=6 ) | | |
|---|---|---|---|---|---|---|---|---|---|
| Tests | Org | CLQ5 | CLQ8 | Org | CLQ5 | CLQ8 | Org | CLQ5 | CLQ8 |
| Wald | 0.6 | 0.8 | 0.8 | 0.6 | 0.9 | 0.9 | 0.6 | 0.9 | 0.9 |
| PC80 | 0.8 | 0.8 | 0.8 | 0.8 | 0.9 | 0.9 | 1.0 | 0.9 | 0.9 |
| MLC-B5 | 0.8 | 0.8 | 0.8 | 0.9 | 0.9 | 0.9 | 0.9 | 0.9 | 0.9 |
| MLC-Z5 | 0.8 | 0.8 | 0.8 | 0.9 | 0.9 | 0.9 | 0.9 | 0.9 | 0.9 |
| MLC-B8 | 0.8 | 0.8 | 0.8 | 0.9 | 0.9 | 0.9 | 0.9 | 0.9 | 0.9 |
| MLC-Z8 | 0.8 | 0.8 | 0.8 | 0.9 | 0.9 | 0.9 | 0.9 | 0.9 | 0.9 |
| SSBw | 0.8 | 0.8 | 0.8 | 0.9 | 0.9 | 0.9 | 0.9 | 1.0 | 0.9 |
| SKAT | 0.8 | 0.8 | 0.8 | 0.9 | 0.8 | 0.8 | 0.9 | 0.9 | 0.7 |
| SKAT-O | 0.8 | 0.8 | 0.7 | 0.8 | 0.8 | 0.8 | 0.9 | 0.8 | 0.7 |

**TABLE S18** Empirical power of gene-based tests using original data and two DRLPC processed data (CLQcut=0.5 and 0.8) for 100 genes stratified by gene size (number of SNPs), 2causal model, EAS population.

| | From 1 to 100 ($N_1$=82) | | | From 101 to 200 ($N_2$=12 ) | | | From 201 to 500 ($N_3$=6 ) | | |
|---|---|---|---|---|---|---|---|---|---|
| Tests | Org | CLQ5 | CLQ8 | Org | CLQ5 | CLQ8 | Org | CLQ5 | CLQ8 |
| Wald | 0.6 | 0.8 | 0.8 | 0.6 | 0.9 | 0.9 | 0.6 | 0.9 | 0.9 |
| PC80 | 0.8 | 0.8 | 0.8 | 0.9 | 0.9 | 0.9 | 1.0 | 0.9 | 0.9 |
| MLC-B5 | 0.8 | 0.8 | 0.8 | 0.9 | 0.9 | 0.9 | 0.9 | 0.9 | 0.9 |
| MLC-Z5 | 0.8 | 0.8 | 0.7 | 0.9 | 0.9 | 0.9 | 0.9 | 0.9 | 0.9 |
| MLC-B8 | 0.8 | 0.8 | 0.8 | 0.9 | 0.9 | 0.9 | 0.8 | 0.9 | 0.9 |
| MLC-Z8 | 0.8 | 0.8 | 0.8 | 0.9 | 0.9 | 0.9 | 0.8 | 0.9 | 0.9 |
| SSBw | 0.8 | 0.8 | 0.8 | 0.9 | 0.9 | 0.9 | 1.0 | 1.0 | 1.0 |
| SKAT | 0.8 | 0.8 | 0.7 | 0.9 | 0.8 | 0.8 | 1.0 | 0.9 | 0.8 |
| SKAT-O | 0.8 | 0.8 | 0.7 | 0.9 | 0.8 | 0.8 | 1.0 | 0.9 | 0.8 |

**TABLE S19** Empirical power of gene-based tests using original data and two DRLPC processed data (CLQcut=0.5 and 0.8) for 100 genes stratified by gene size (number of SNPs), 1causal model, AFR population.

| | From 1 to 100 ($N_1$=68) | | | From 101 to 200 ($N_2$=18 ) | | | From 201 to 500 ($N_3$=14 ) | | |
|---|---|---|---|---|---|---|---|---|---|
| Tests | Org | CLQ5 | CLQ8 | Org | CLQ5 | CLQ8 | Org | CLQ5 | CLQ8 |
| Wald | 0.6 | 0.8 | 0.8 | 0.6 | 0.9 | 0.9 | 0.6 | 0.9 | 0.9 |
| PC80 | 0.8 | 0.8 | 0.8 | 0.9 | 0.9 | 0.9 | 0.9 | 0.9 | 0.9 |
| MLC-B5 | 0.8 | 0.8 | 0.8 | 0.9 | 0.9 | 0.9 | 0.9 | 0.9 | 0.9 |
| MLC-Z5 | 0.8 | 0.8 | 0.8 | 0.9 | 0.9 | 0.9 | 0.9 | 0.9 | 0.9 |
| MLC-B8 | 0.7 | 0.8 | 0.8 | 0.8 | 0.9 | 0.9 | 0.8 | 0.9 | 0.9 |
| MLC-Z8 | 0.8 | 0.8 | 0.8 | 0.9 | 0.9 | 0.9 | 0.9 | 0.9 | 0.9 |
| SSBw | 0.8 | 0.8 | 0.8 | 0.9 | 0.9 | 0.9 | 0.9 | 1.0 | 1.0 |
| SKAT | 0.8 | 0.8 | 0.7 | 0.9 | 0.9 | 0.9 | 0.9 | 0.9 | 0.8 |
| SKAT-O | 0.7 | 0.7 | 0.6 | 0.8 | 0.9 | 0.9 | 0.9 | 0.9 | 0.8 |

**TABLE S20** Empirical power of gene-based tests using original data and two DRLPC processed data (CLQcut=0.5 and 0.8) for 100 genes stratified by gene size (number of SNPs), 2causal model, AFR population.

| | From 1 to 100 ($N_1$=68) | | | From 101 to 200 ($N_2$=18 ) | | | From 201 to 500 ($N_3$=14 ) | | |
|---|---|---|---|---|---|---|---|---|---|
| Tests | Org | CLQ5 | CLQ8 | Org | CLQ5 | CLQ8 | Org | CLQ5 | CLQ8 |
| Wald | 0.6 | 0.8 | 0.8 | 0.6 | 0.9 | 0.9 | 0.6 | 0.9 | 0.8 |
| PC80 | 0.8 | 0.8 | 0.8 | 0.9 | 0.9 | 0.9 | 1.0 | 0.9 | 0.9 |
| MLC-B5 | 0.8 | 0.8 | 0.8 | 0.9 | 0.9 | 0.9 | 0.9 | 0.9 | 0.9 |
| MLC-Z5 | 0.8 | 0.8 | 0.8 | 0.9 | 0.9 | 0.9 | 0.9 | 0.9 | 0.9 |
| MLC-B8 | 0.7 | 0.8 | 0.8 | 0.8 | 0.9 | 0.9 | 0.8 | 0.9 | 0.9 |
| MLC-Z8 | 0.8 | 0.8 | 0.8 | 0.8 | 0.9 | 0.9 | 0.9 | 0.9 | 0.9 |
| SSBw | 0.8 | 0.8 | 0.8 | 0.8 | 0.9 | 0.9 | 0.9 | 1.0 | 1.0 |
| SKAT | 0.8 | 0.8 | 0.7 | 0.8 | 0.9 | 0.9 | 0.9 | 0.9 | 0.8 |
| SKAT-O | 0.7 | 0.7 | 0.6 | 0.8 | 0.8 | 0.9 | 0.9 | 0.9 | 0.8 |

#### 2.2 | Supplementary Figures

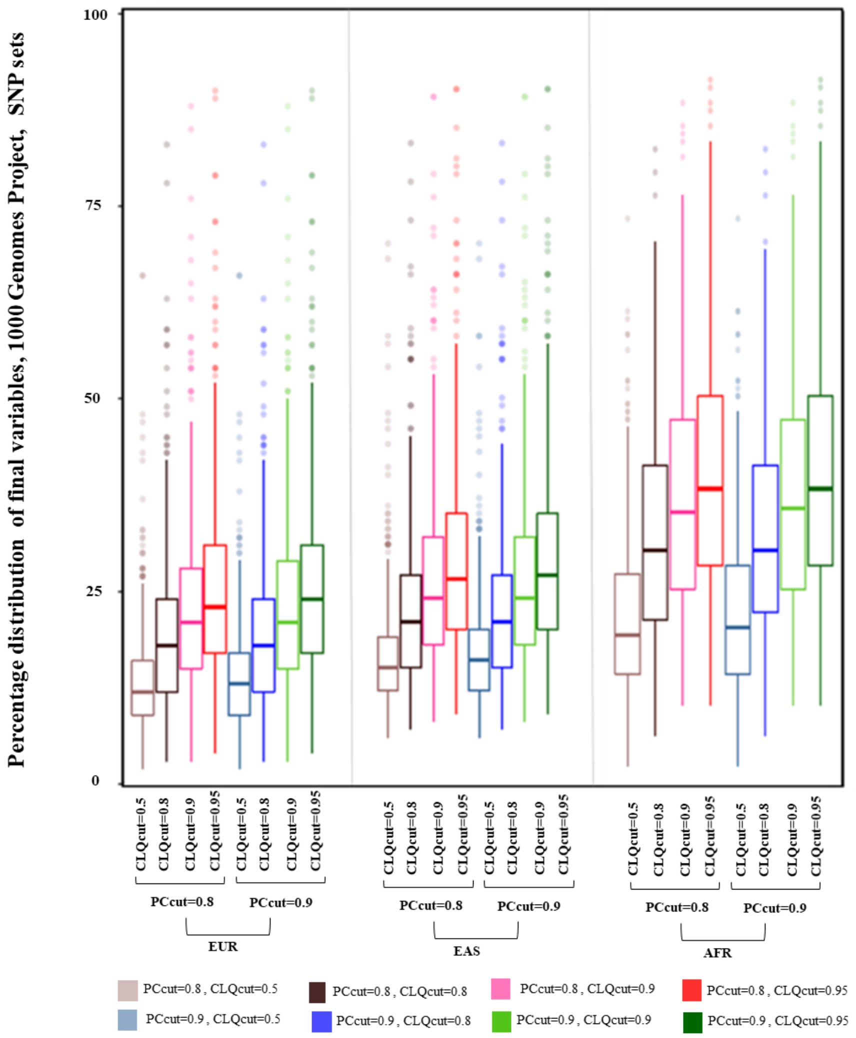

**FIGURE S1** The percentage distribution of final variables after the DRLPC process compared to the number of SNPs in 100 SNP sets of 1000 Genomes Project, chromosome 22, with four threshold values for CLQcut (0.5, 0.8, 0.9, 0.95), two threshold values (0.8, 0.9) for PCcut, and for three super-populations (EUR, EAS, AFR)

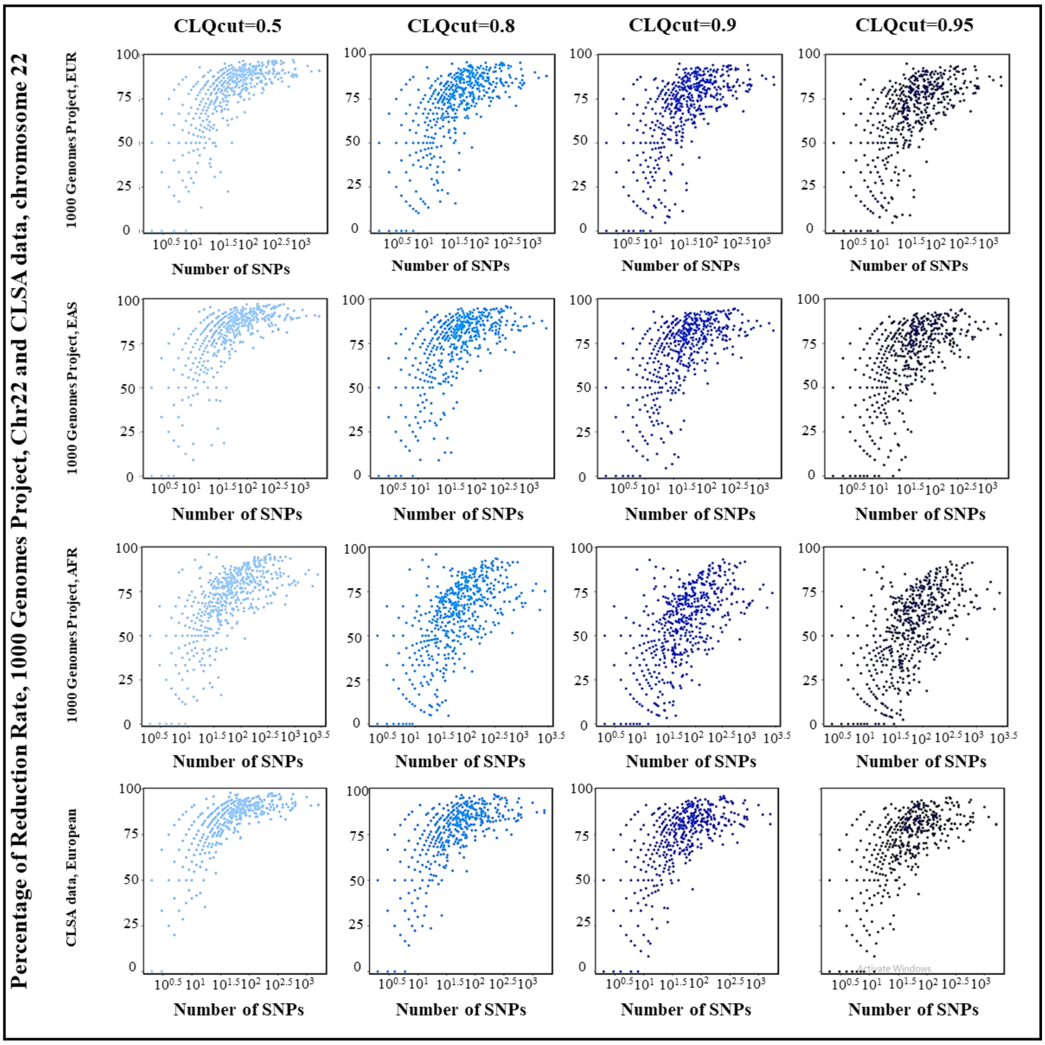

**FIGURE S2** The reduction rates after the DRLPC process plotted with the number of SNPs in the original dataset (gene size) for gene-based datasets of CLSA data, chromosome 22, with four threshold values for CLQcut (0.5, 0.8, 0.9, 0.95), a threshold value 0.8 for PCcut and for three super-populations (EUR, EAS, AFR) and European ancestry.

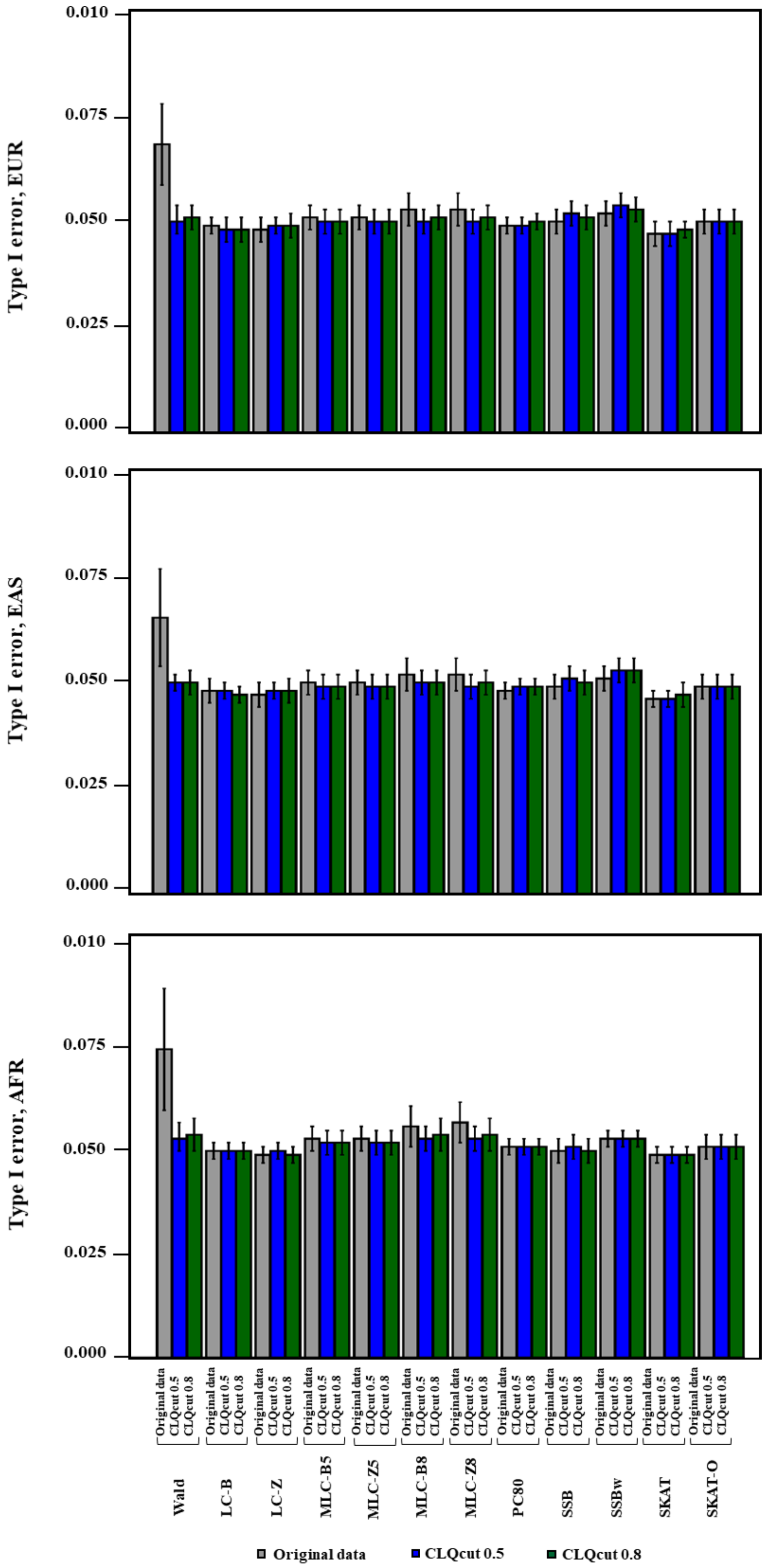

**FIGURE S3** Average empirical type I error of gene-based statistics (N=10,000 replicate) obtained at the 0.05 level averaged over 100 genes, comparison between original data and two processed data considering CLQcut point 0.5 and 0.8 with one PCcut value 0.8, for three populations.

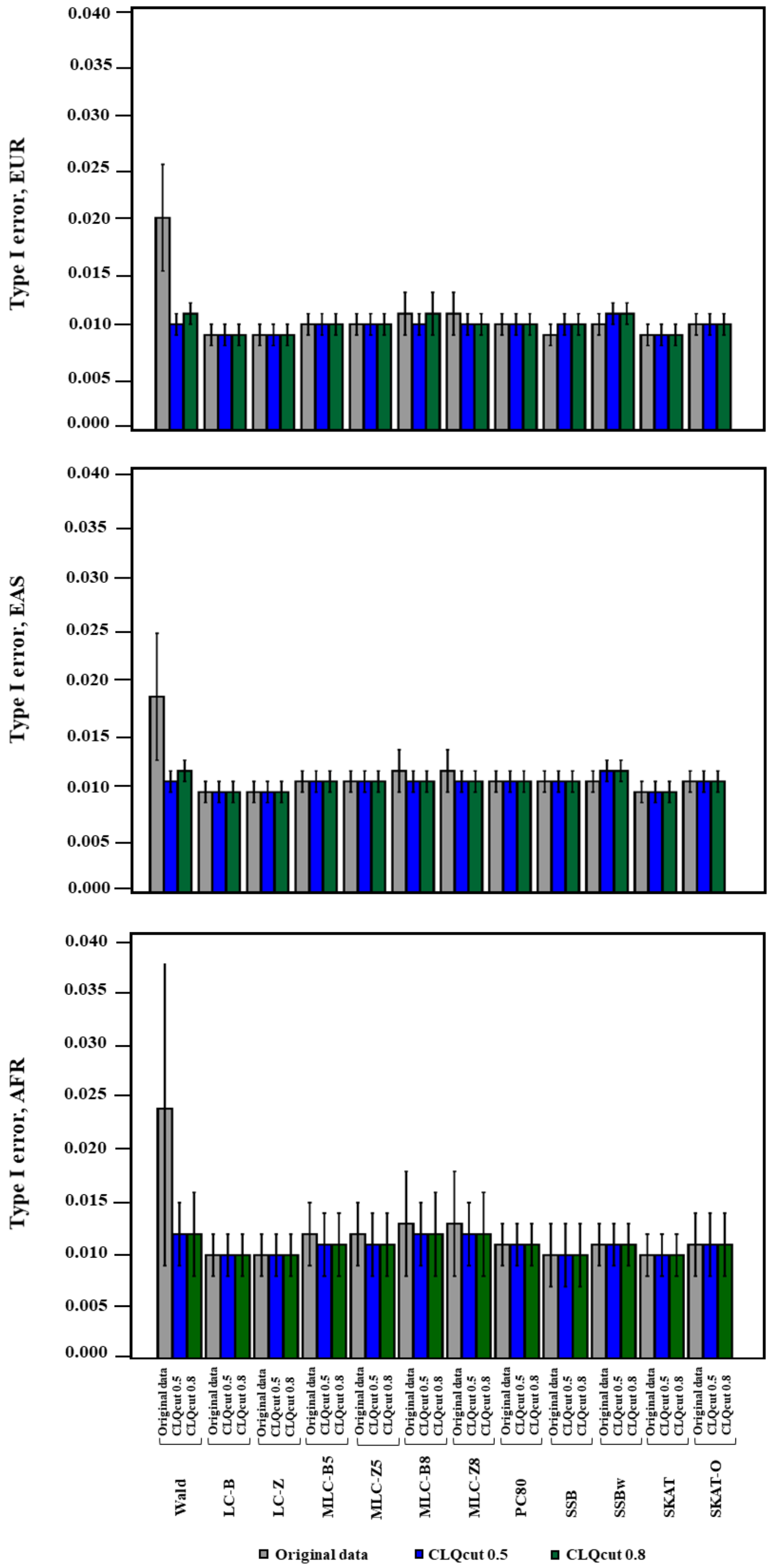

**FIGURE S4** Average empirical type I error of gene-based statistics (N=10,000 replicate) obtained at the 0.01 level averaged over 100 genes, comparison between three super-populations, with original data and two processed data considering CLQcut point 0.5 and 0.8 with one PCcut value 0.8.

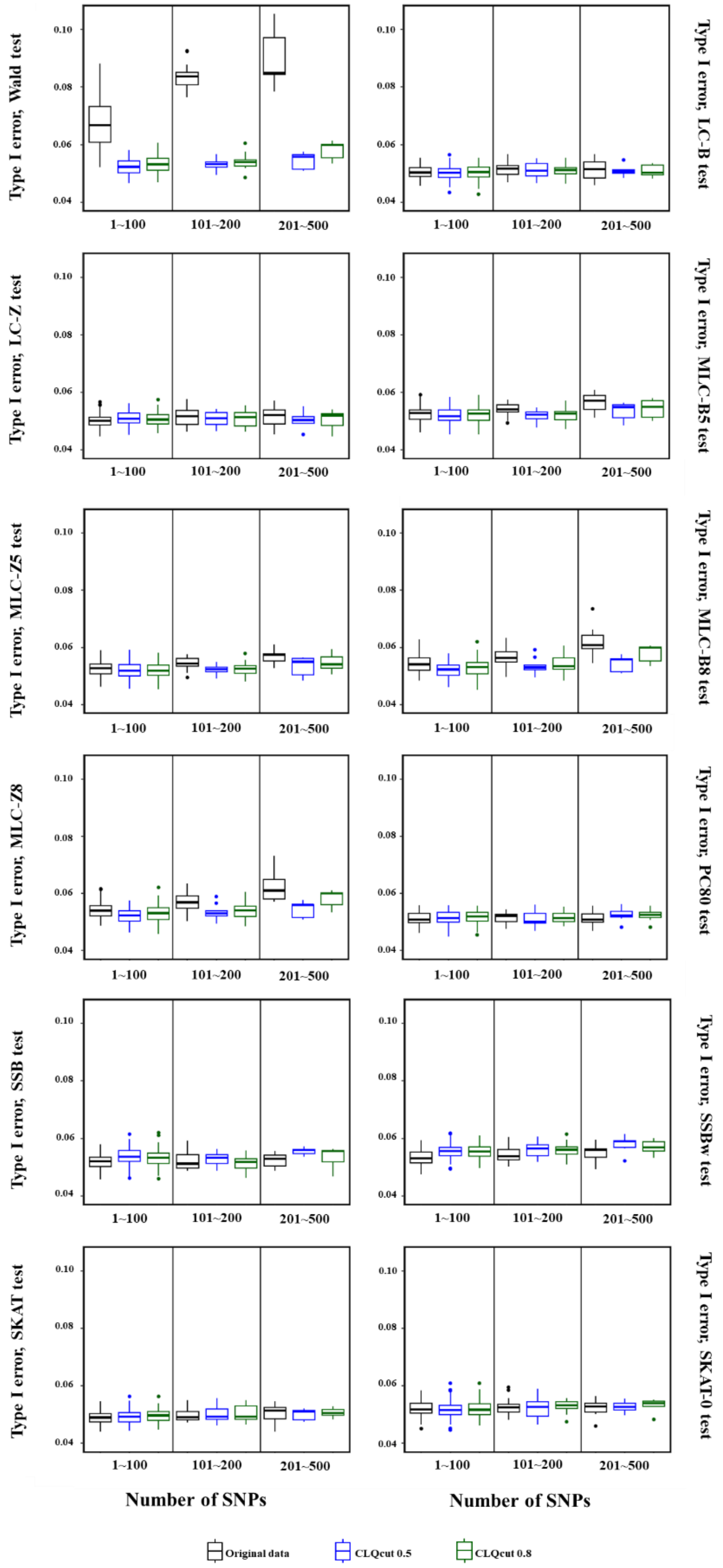

**FIGURE S5** Simulation study: Type I error of gene-based statistics (N=10,000 replicate) obtained at the nominal level $\alpha$ =0.05, based on the gene size over 100 genes, for original data and two processed data with CLQcut point 0.5 and 0.8 with one PCcut value 0.8, EUR population.

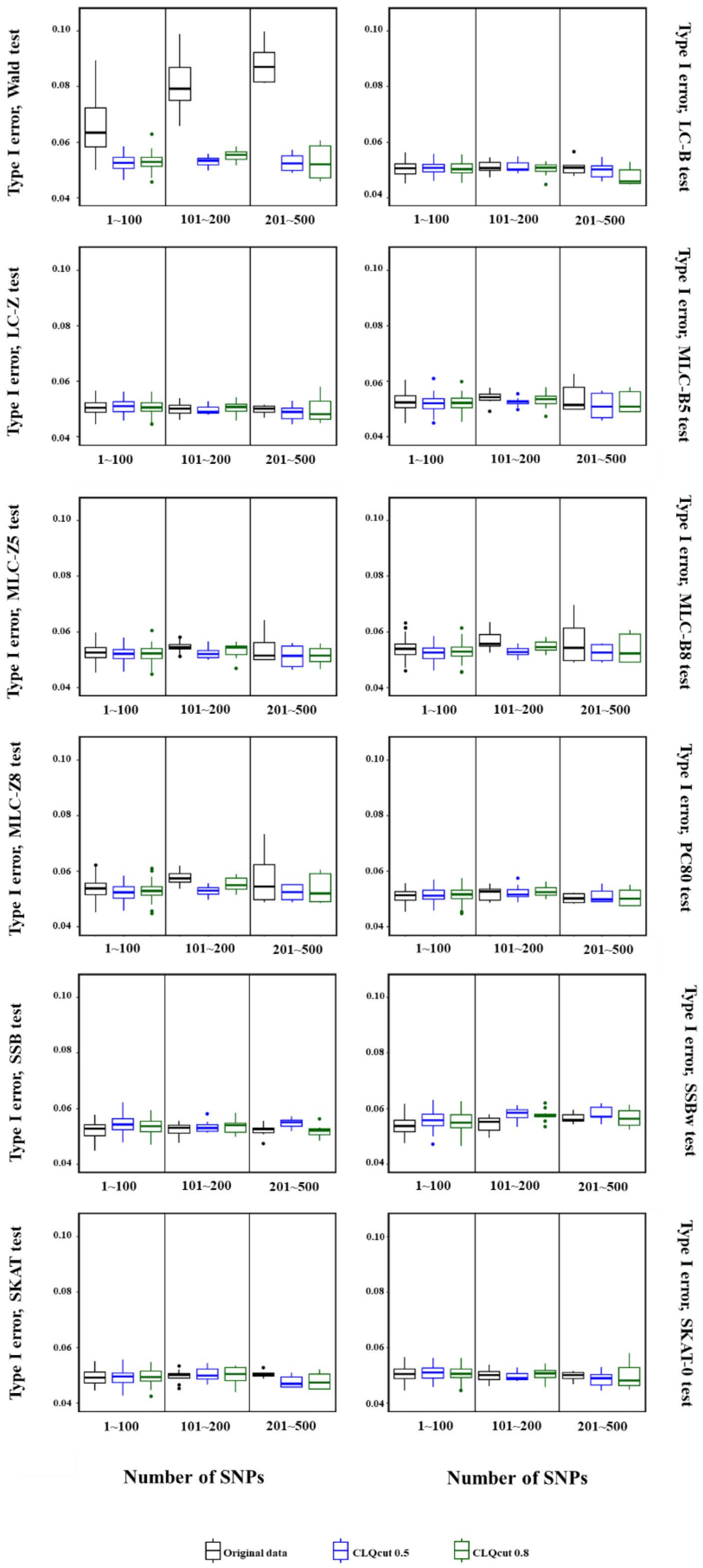

**FIGURE S6** Simulation study: Type I error of gene-based statistics (N=10,000 replicate) obtained at the nominal level $\alpha$ =0.05, based on the gene size over 100 genes, for original data and two processed data with CLQcut point 0.5 and 0.8 with one PCcut value 0.8, EAS population.

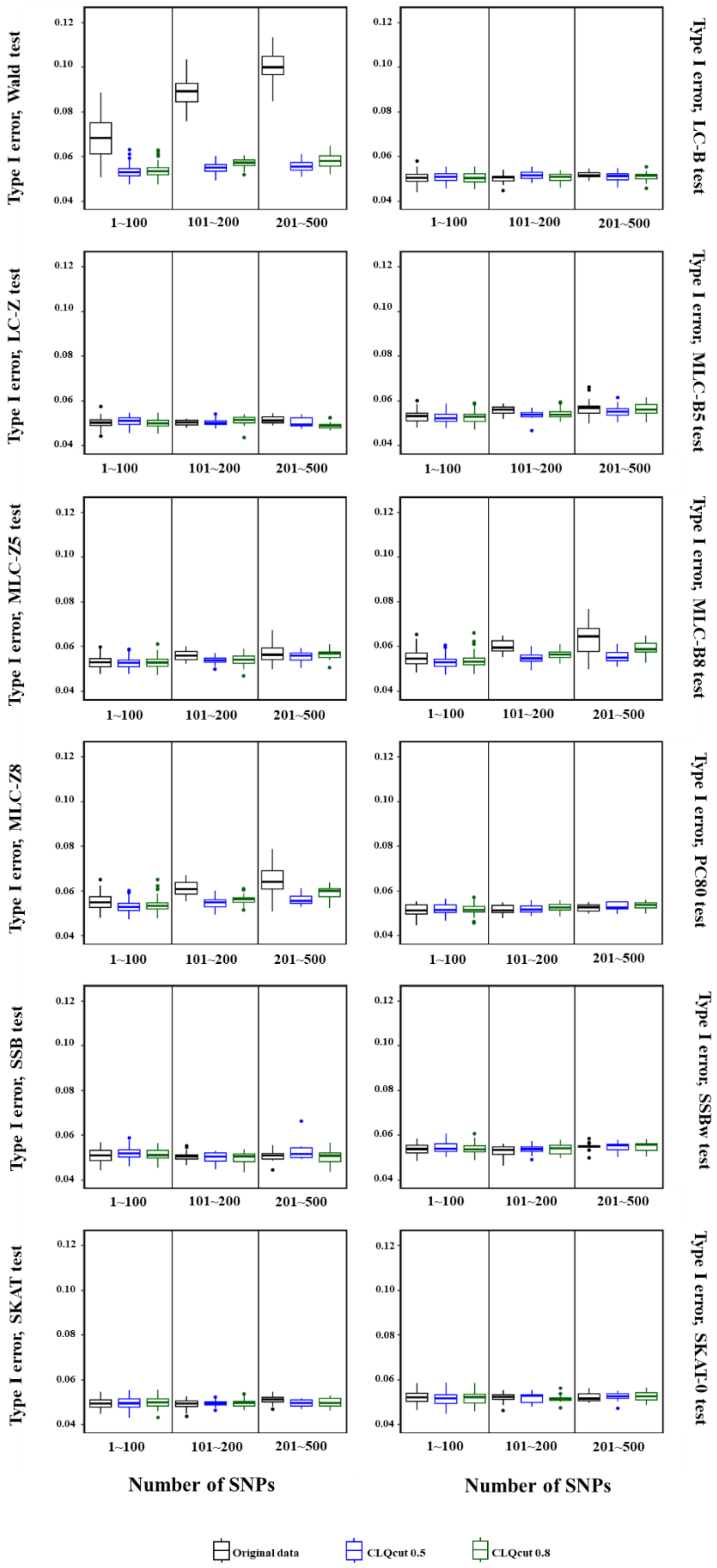

**FIGURE S7** Simulation study: Type I error of gene-based statistics (N=10,000 replicate) obtained at the nominal level $\alpha$ =0.05, based on the gene size over 100 genes, for original data and two processed data with CLQcut point 0.5 and 0.8 with one PCcut value 0.8, AFR population.

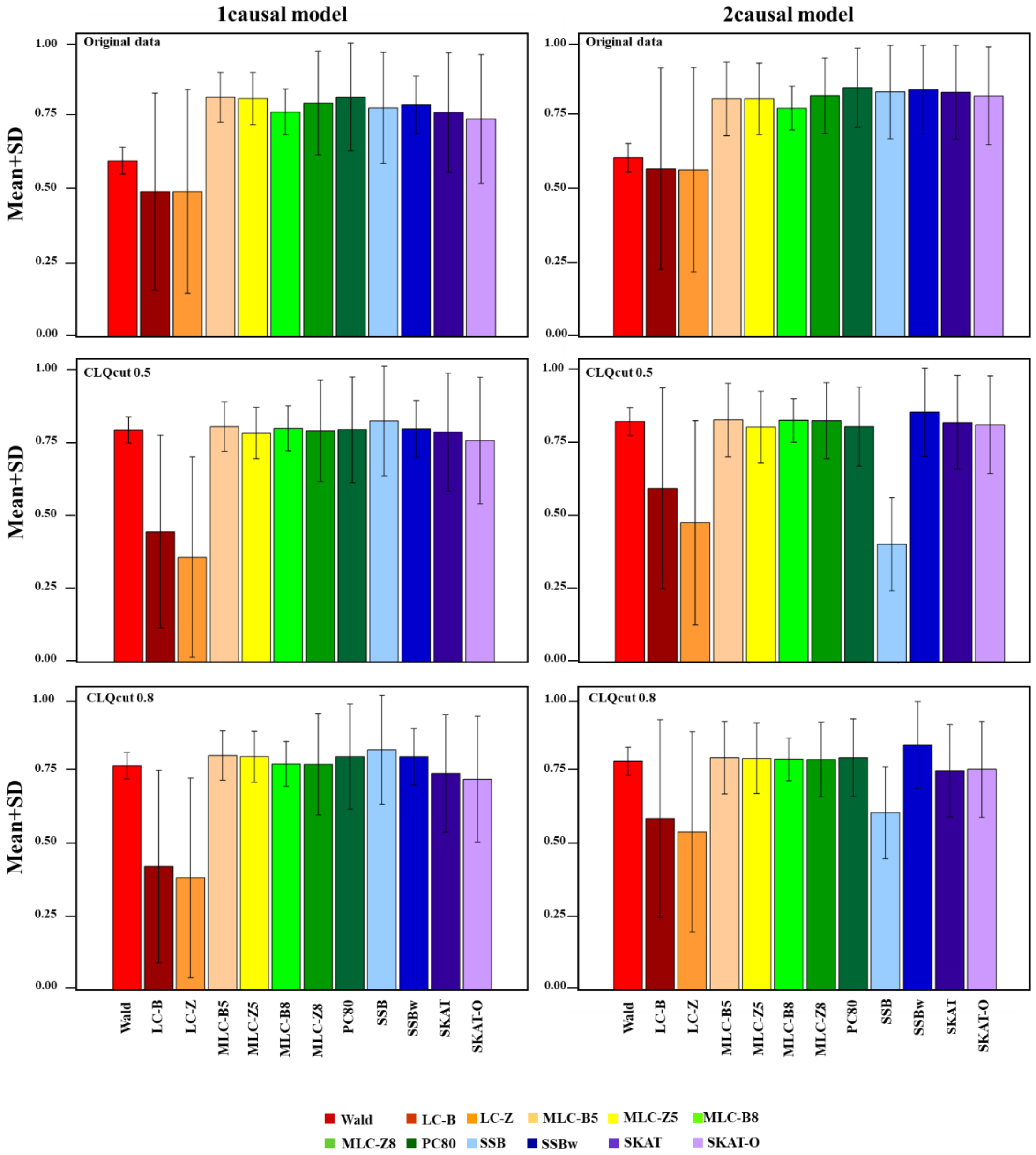

342 **FIGURE S8** Average empirical power of multi-marker statistics tests under two trait models (1causal and
343 2causal) for original data and two DRLPC processed data for the EUR population.
344

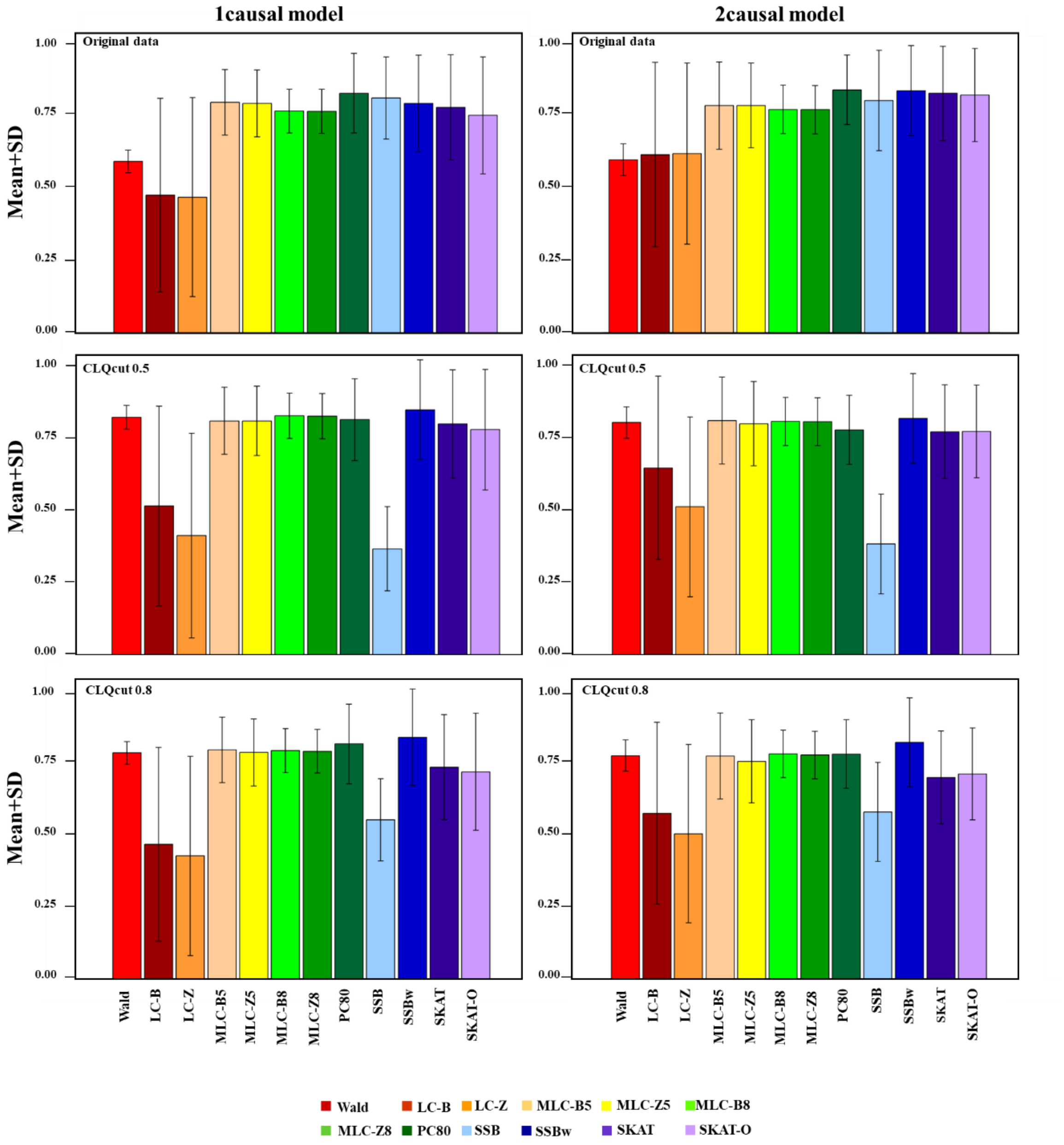

**FIGURE S9** Average empirical power of multi-marker statistics tests under two trait models (1causal and 2causal) for original data and two DRLPC processed data for the EAS population.

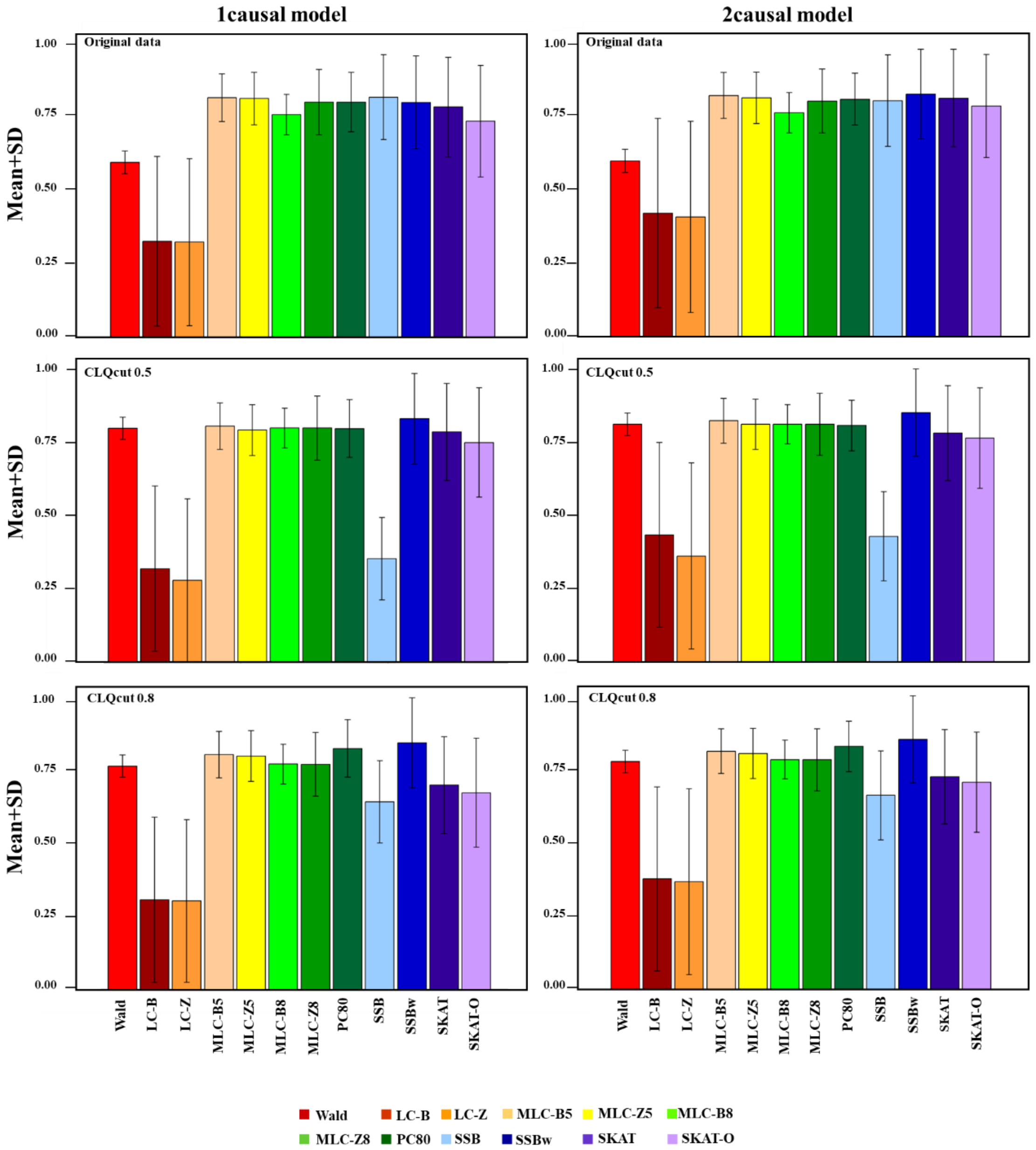

**FIGURE S10** Average empirical power of multi-marker statistics tests under two trait models (1causal and 2causal) for original data and two DRLPC processed data for the AFR population.

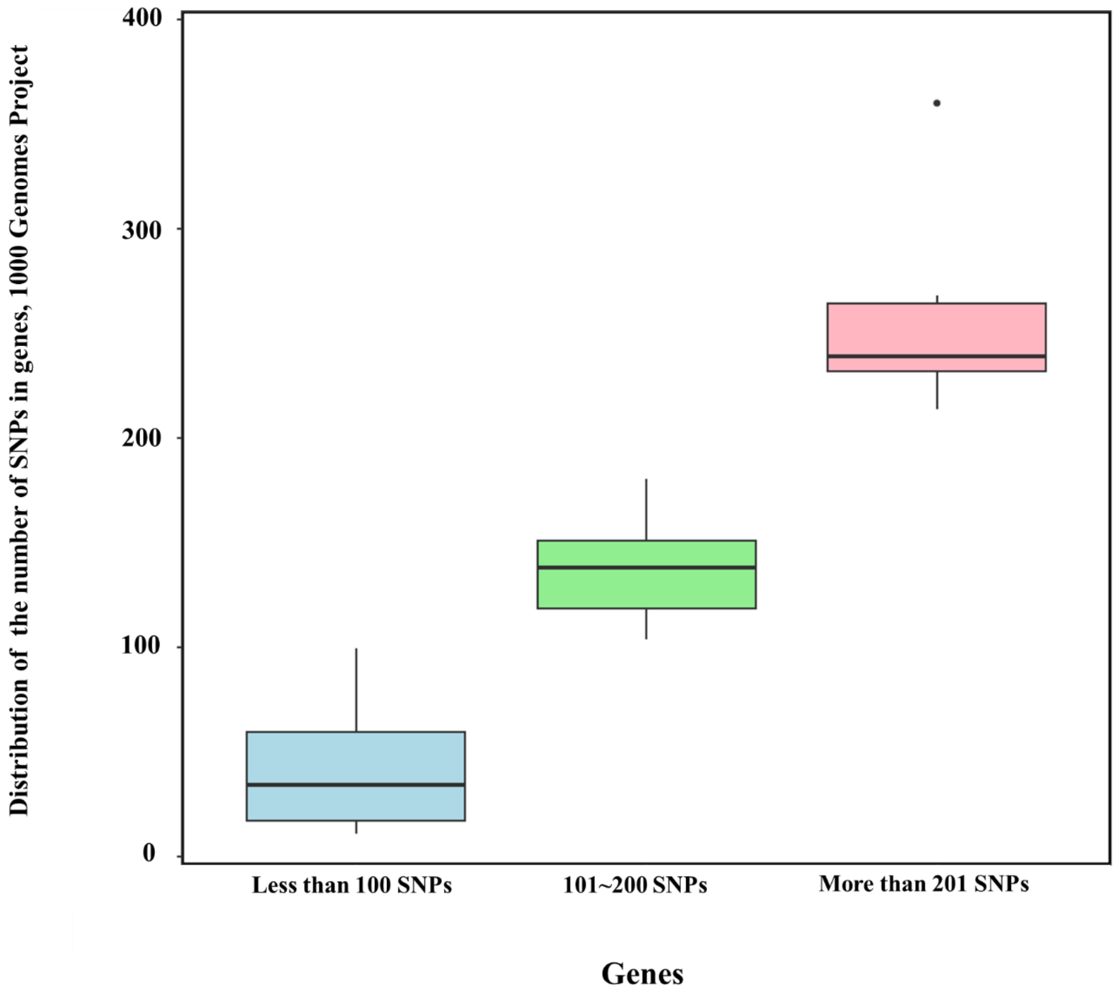

**FIGURE S11** Distribution of the number of SNPs for three gene's group (genes with 1~100, 101~200, 201~500 SNPs) for 1000 Genomes Project data, chromosome 22, EUR population.

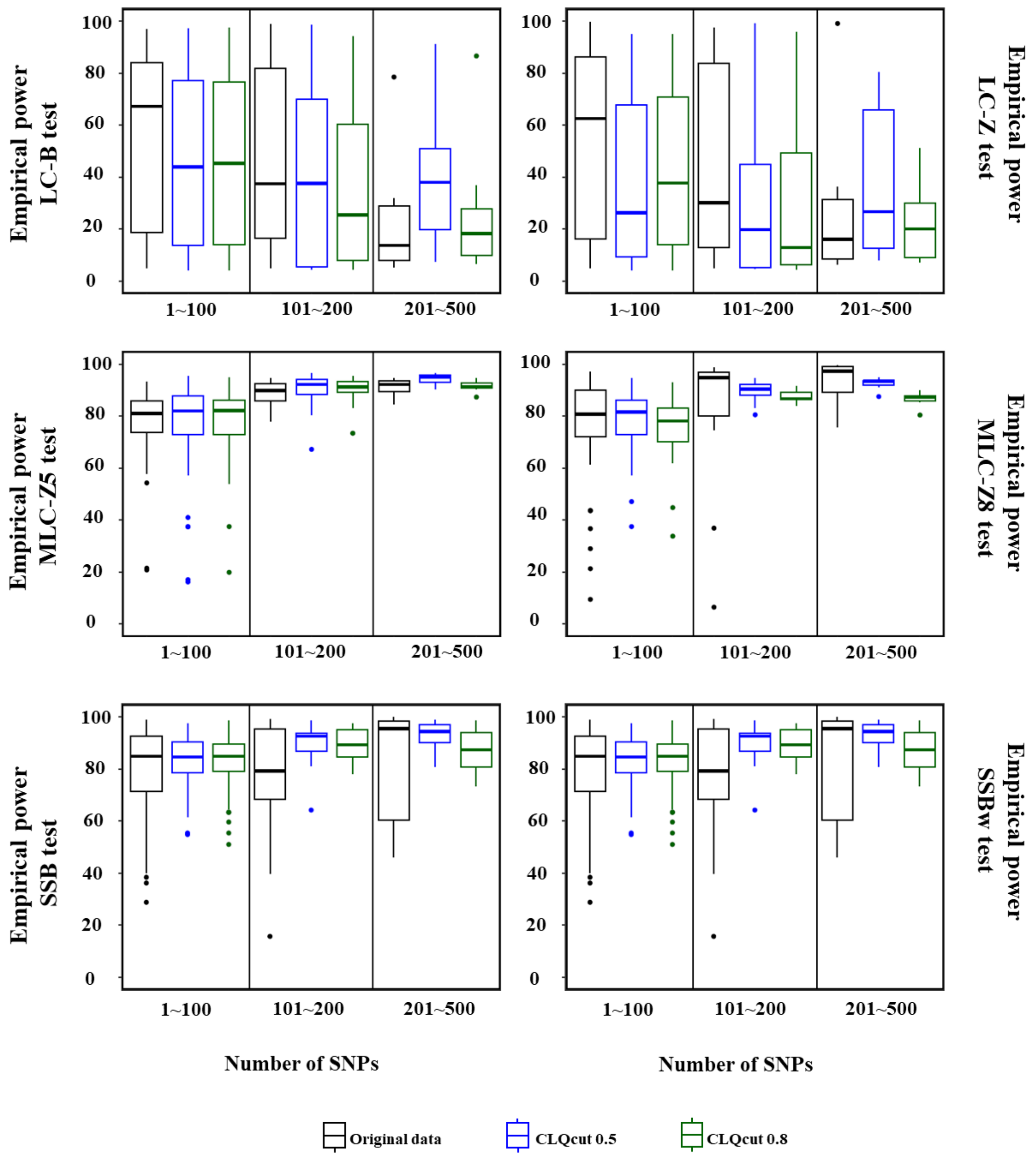

**FIGURE S12** Percentage of average empirical power for gene-based statistics (N=1000 replicate) at the 0.05 level for EUR population, 1causal model, averaged across three groups of 100 genes based on the number of SNPs in the gene.

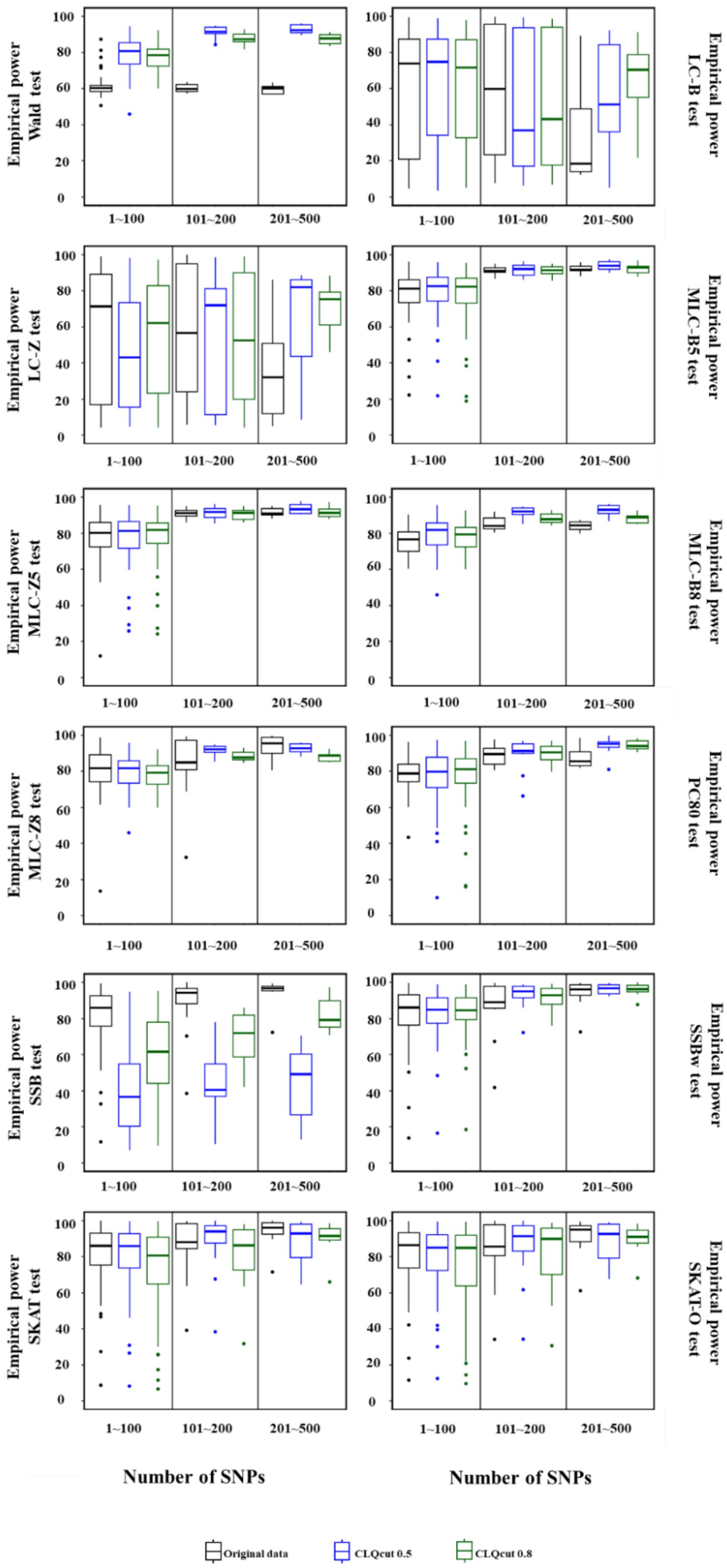

**FIGURE S13** Percentage of average empirical power for gene-based statistics (N=1000 replicate) at the 0.05 level for EUR population, 2causal model, averaged across three groups of 100 genes based on the number of SNPs in the gene.

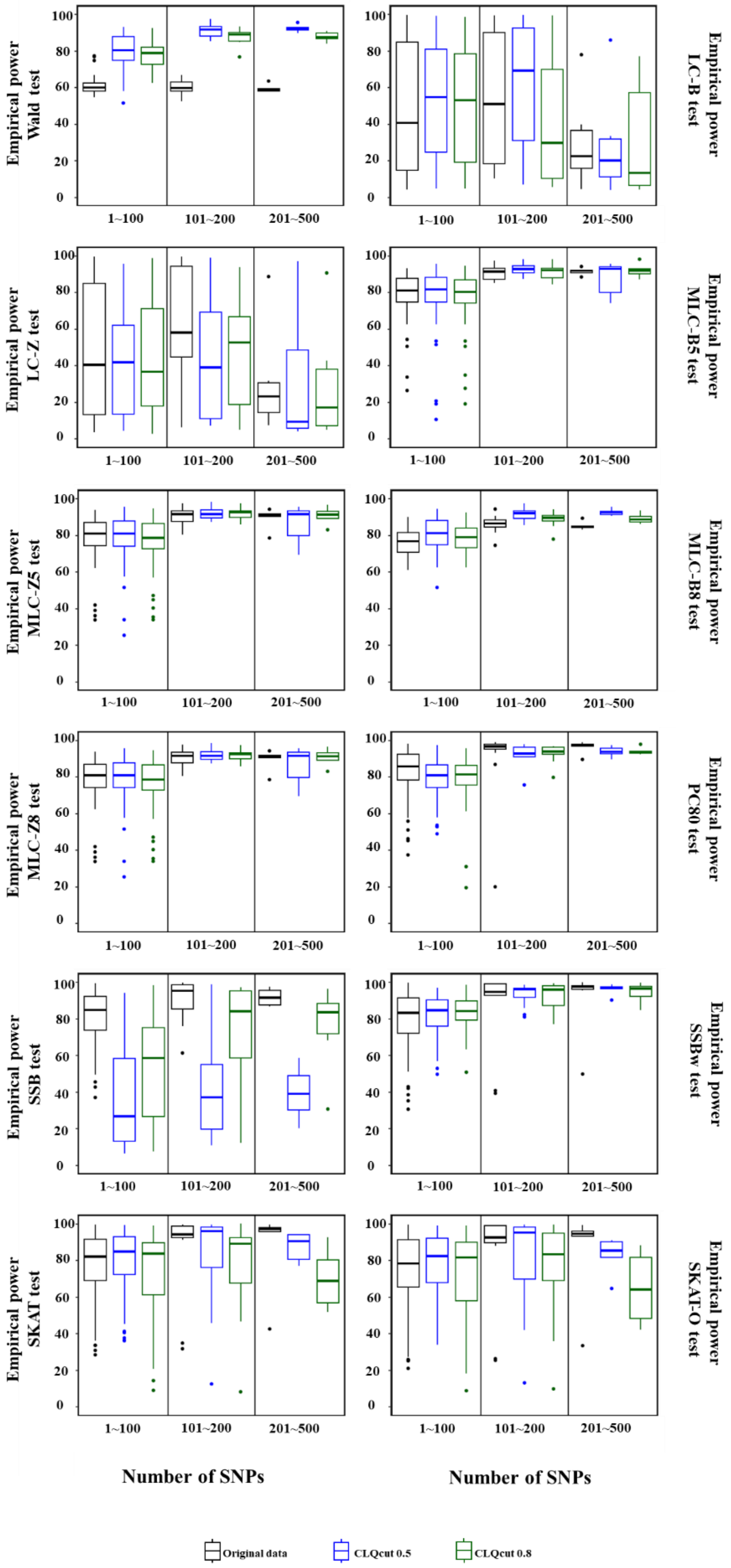

**FIGURE S14** Percentage average empirical power for gene-based statistics (N=1000 replicate) at the 0.05 level for EAS population, 1causal model, averaged across three groups of 100 genes based on the number of SNPs in the gene.

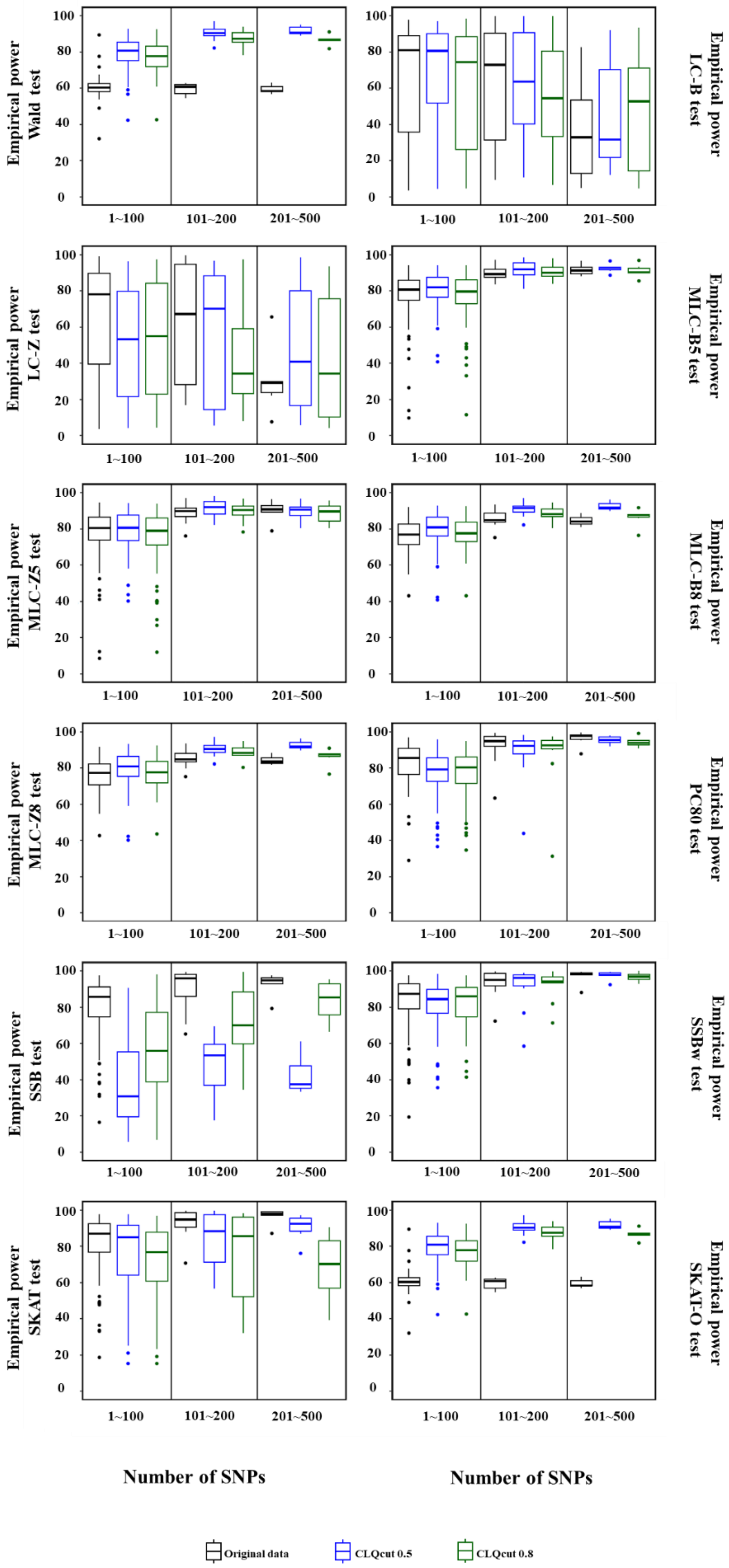

**FIGURE S15** Percentage of average empirical power for gene-based statistics (N=1000 replicate) at the 0.05 level for EAS population, 2causal model, averaged across three groups of 100 genes based on the number of SNPs in the gene.

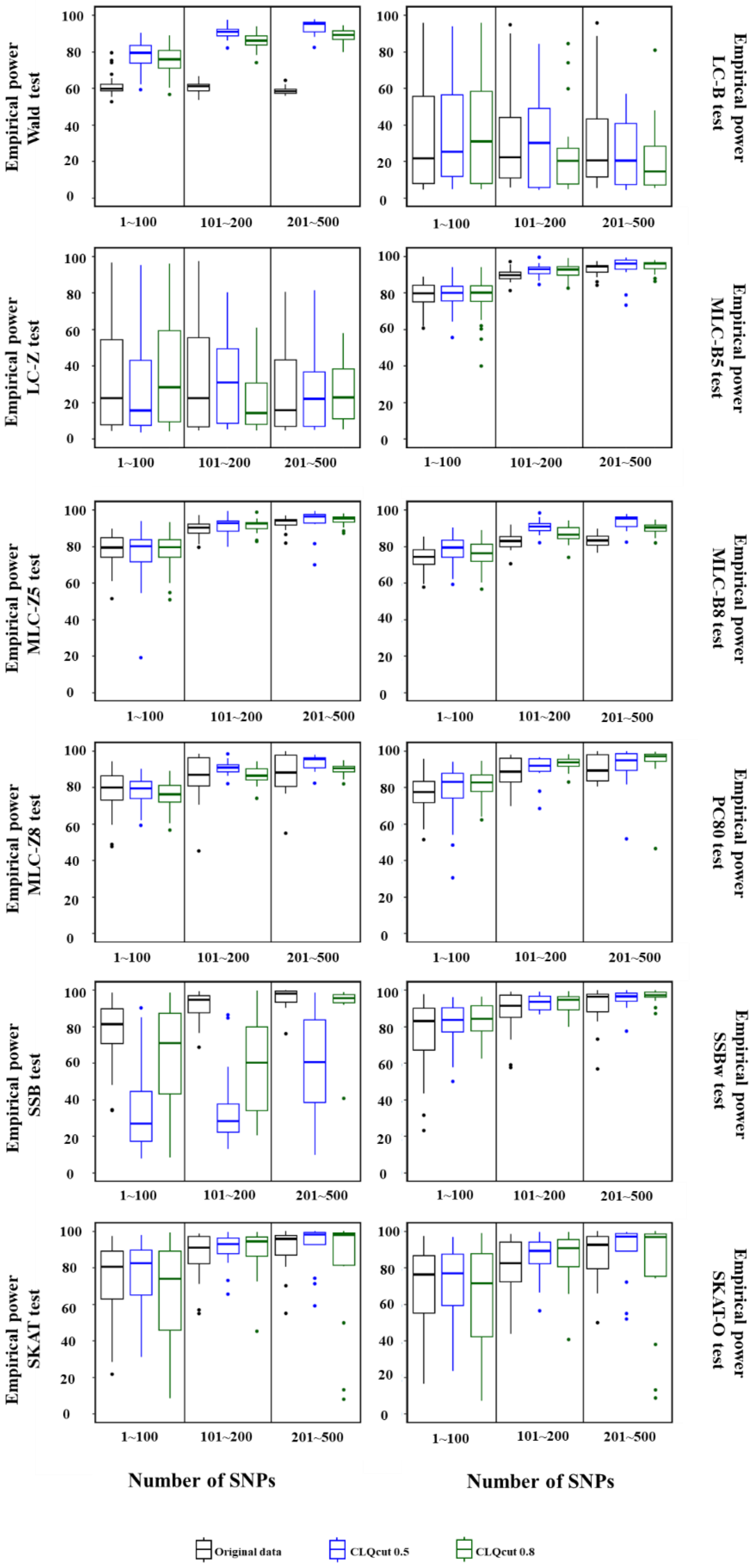

**FIGURE S16** Percentage of average empirical power for gene-based statistics (N=1000 replicate) at the 0.05 level for AFR population, 1causal model, averaged across three groups of 100 genes based on the number of SNPs in the gene.

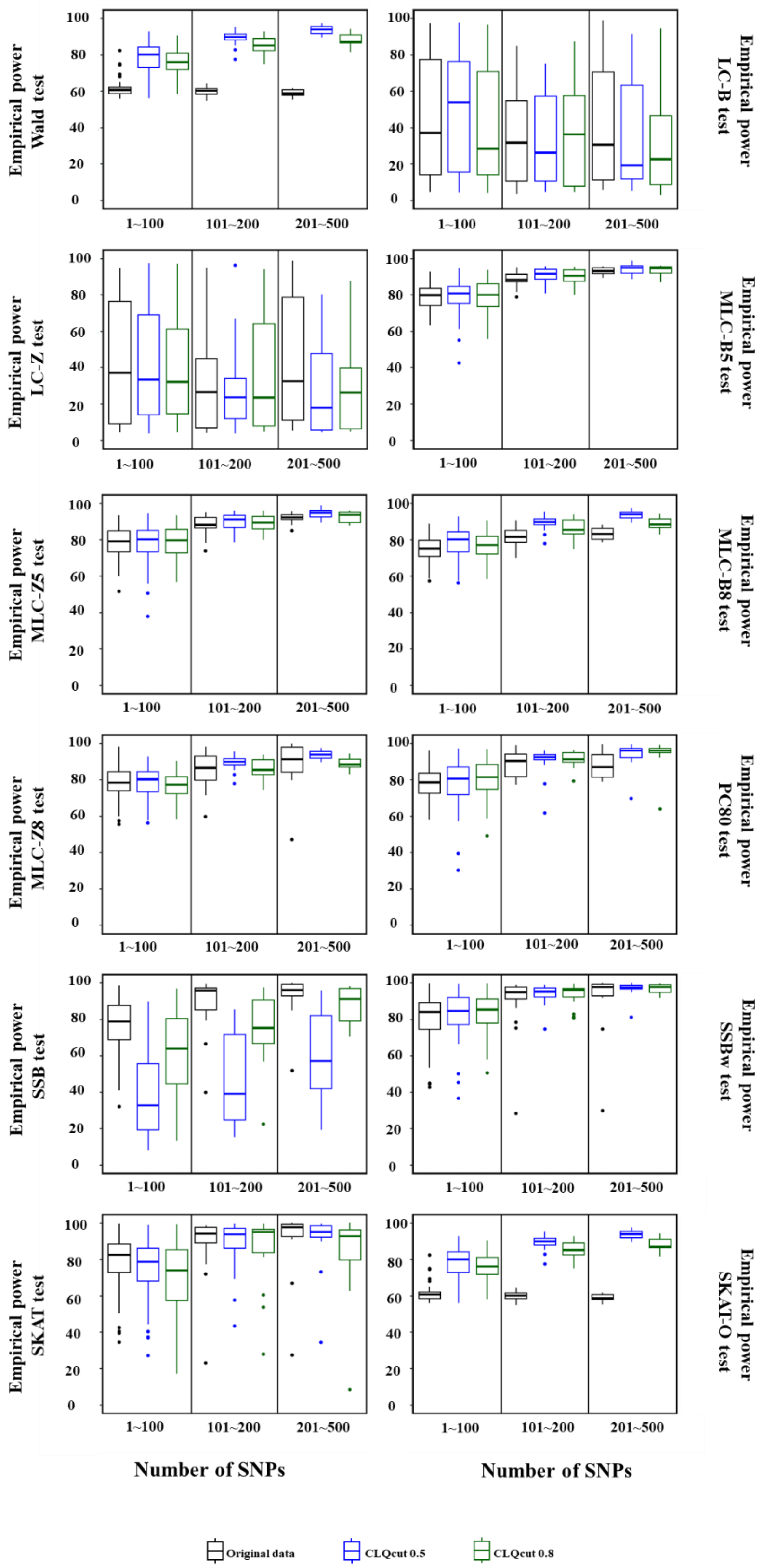

**FIGURE S17** Percentage of average empirical power for gene-based statistics (N=1000 replicate) at the 0.05 level for AFR population, 2causal model, averaged across three groups of 100 genes based on the number of SNPs in the gene.

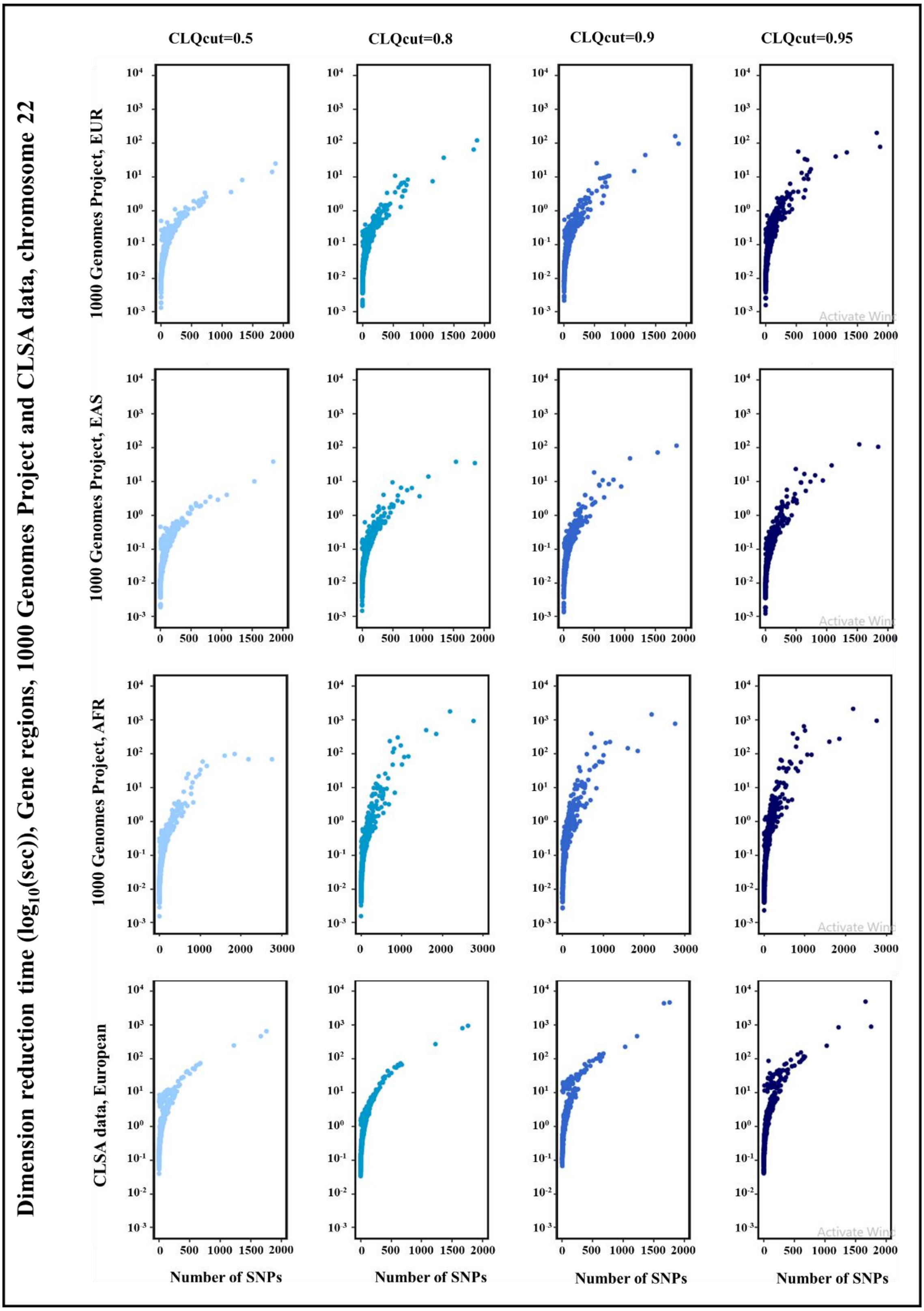

**FIGURE S18** Computational time for dimension reduction using the DRLPC for gene regions, with four threshold values for CLQcut (0.5, 0.8, 0.9, 0.95) and a threshold values 0.9 for PCcut, 1000 Genomes Project, three super-populations and CLSA data European ancestry, chromosome 22. The CLSA sample size is 17,779 whereas 1000 Genomes sample sizes are 503-661.

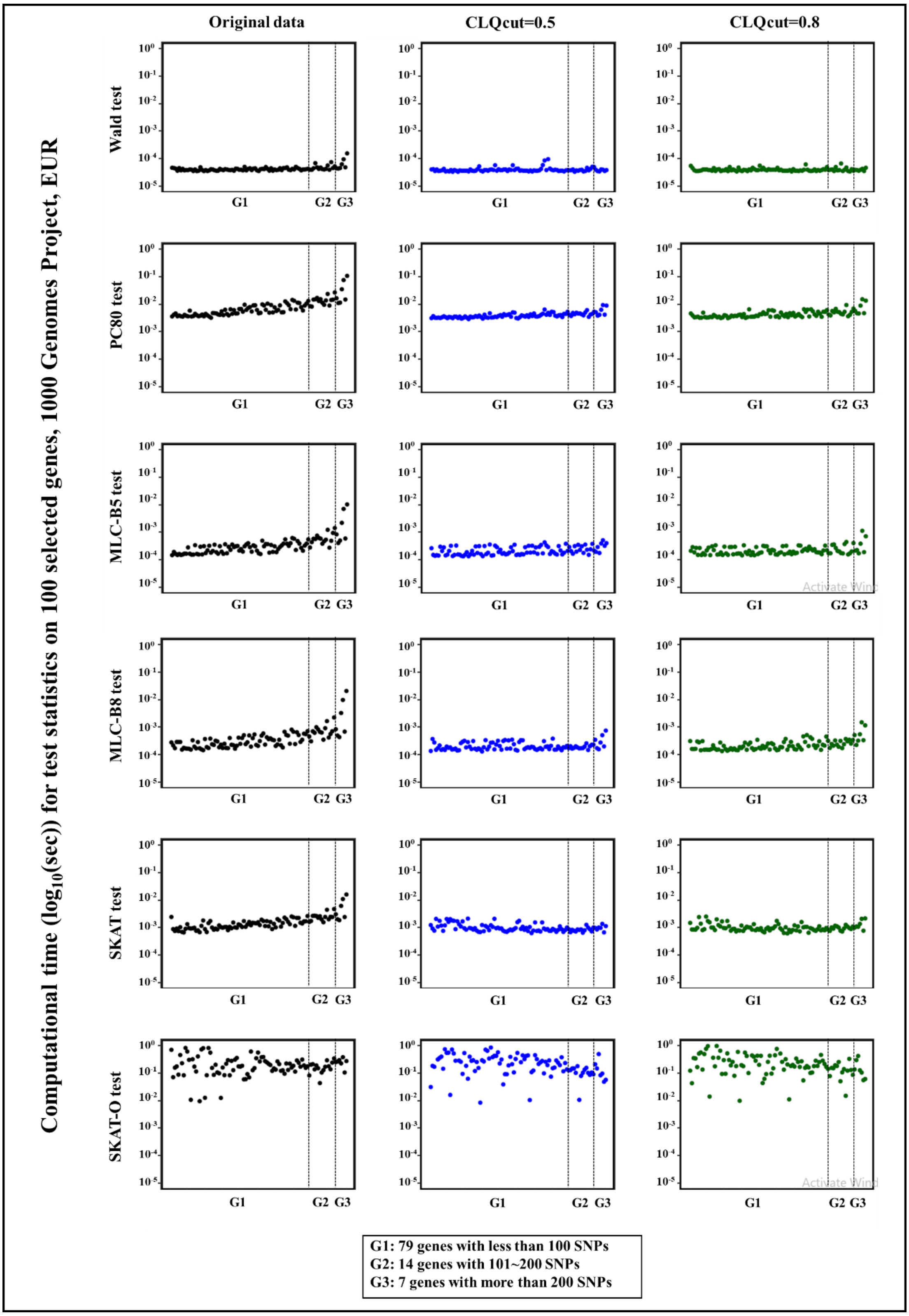

**FIGURE S19** The computational time for test statistics in a single replication of 2causal model on 100 selected genes, 79 genes with less than 101 SNPs, 14 genes with 101~200 SNPs, and seven genes with more than 200 SNPs, using the original data and the DRLPC processed data with two threshold values for CLQcut (0.5, 0.8) and a threshold values 0.8 for PCcut, 1000 Genomes Project, EUR, chromosome 22. The x-axix represents the original gene size, sorted based on the number of SNPs.

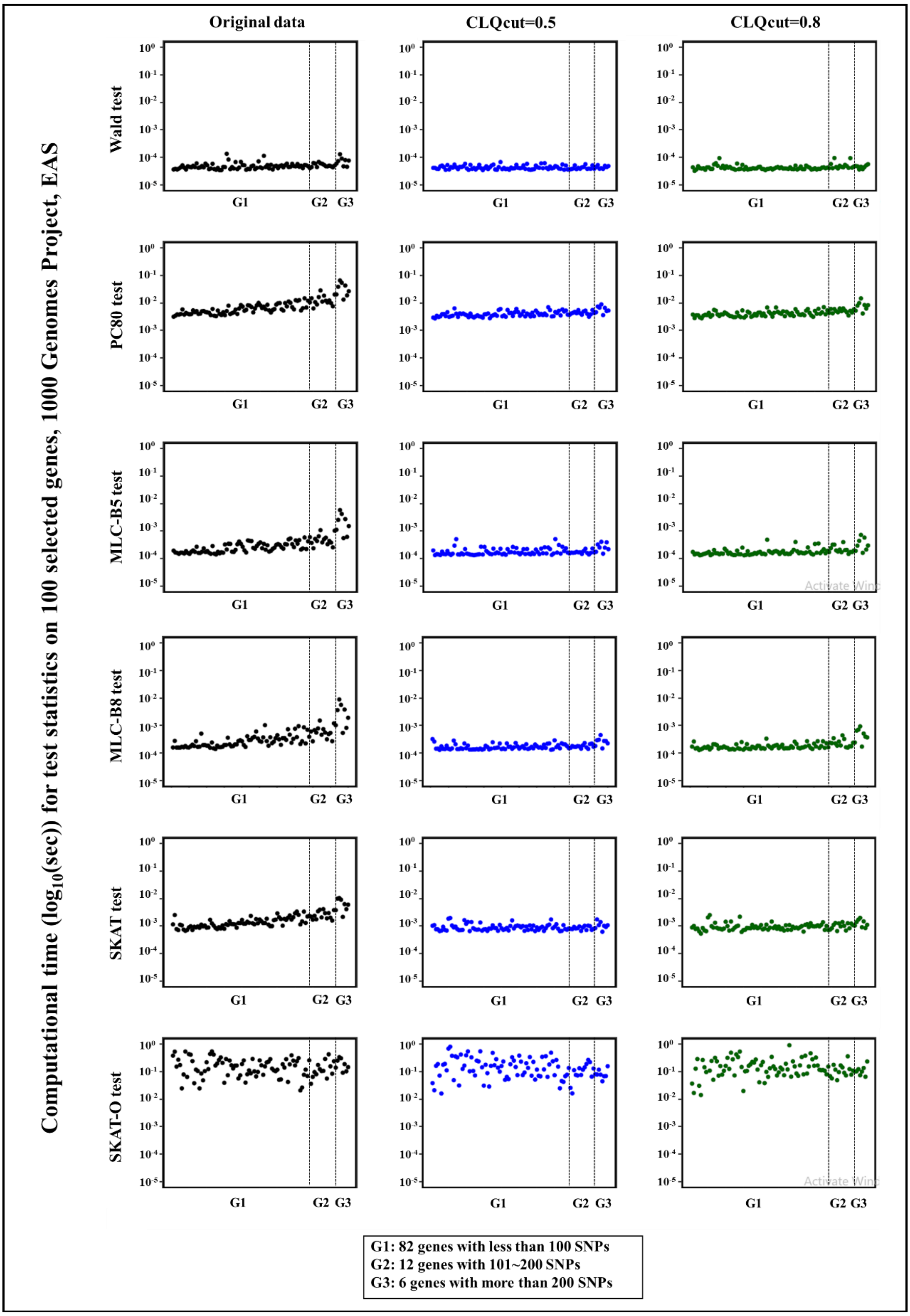

**FIGURE S20** The computational time for test statistics in a single replication of 1casual model on 100 selected genes, 82 genes with less than 101 SNPs, 12 genes with 101~200 SNPs, and six genes with more than 200 SNPs, using the original data and the DRLPC processed data with two threshold values for CLQcut (0.5, 0.8) and a threshold values 0.8 for PCcut, 1000 Genomes Project, EAS, chromosome 22. The x-axix represents the original gene size, sorted based on the number of SNPs.

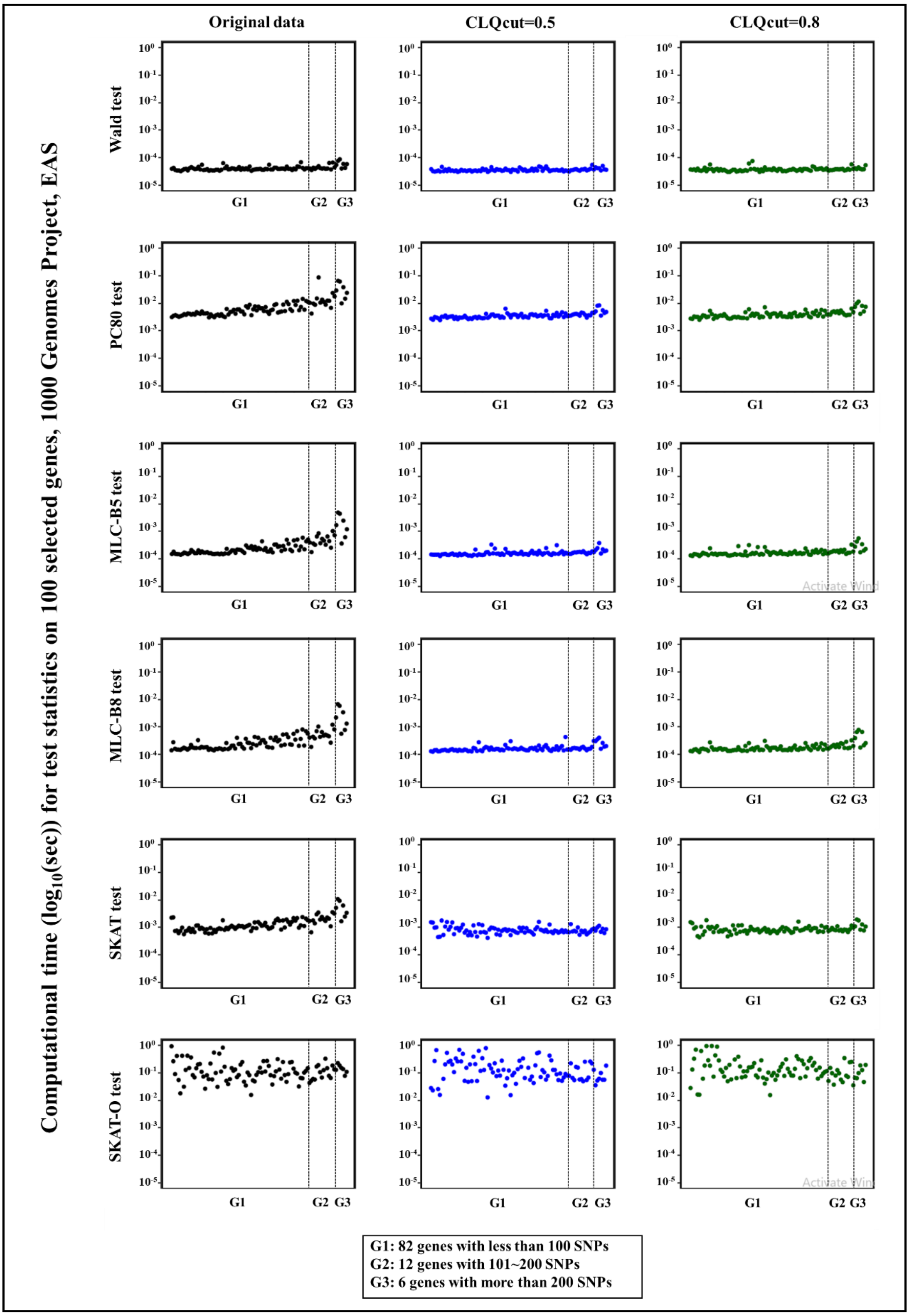

**FIGURE S21** The computational time for test statistics in a single replication of 2casual model on 100 selected genes, 82 genes with less than 101 SNPs, 12 genes with 101~200 SNPs, and six genes with more than 200 SNPs, using the original data and the DRLPC processed data with two threshold values for CLQcut (0.5, 0.8) and a threshold values 0.8 for PCcut, 1000 Genomes Project, EAS, chromosome 22. The x-axix represents the original gene size, sorted based on the number of SNPs.

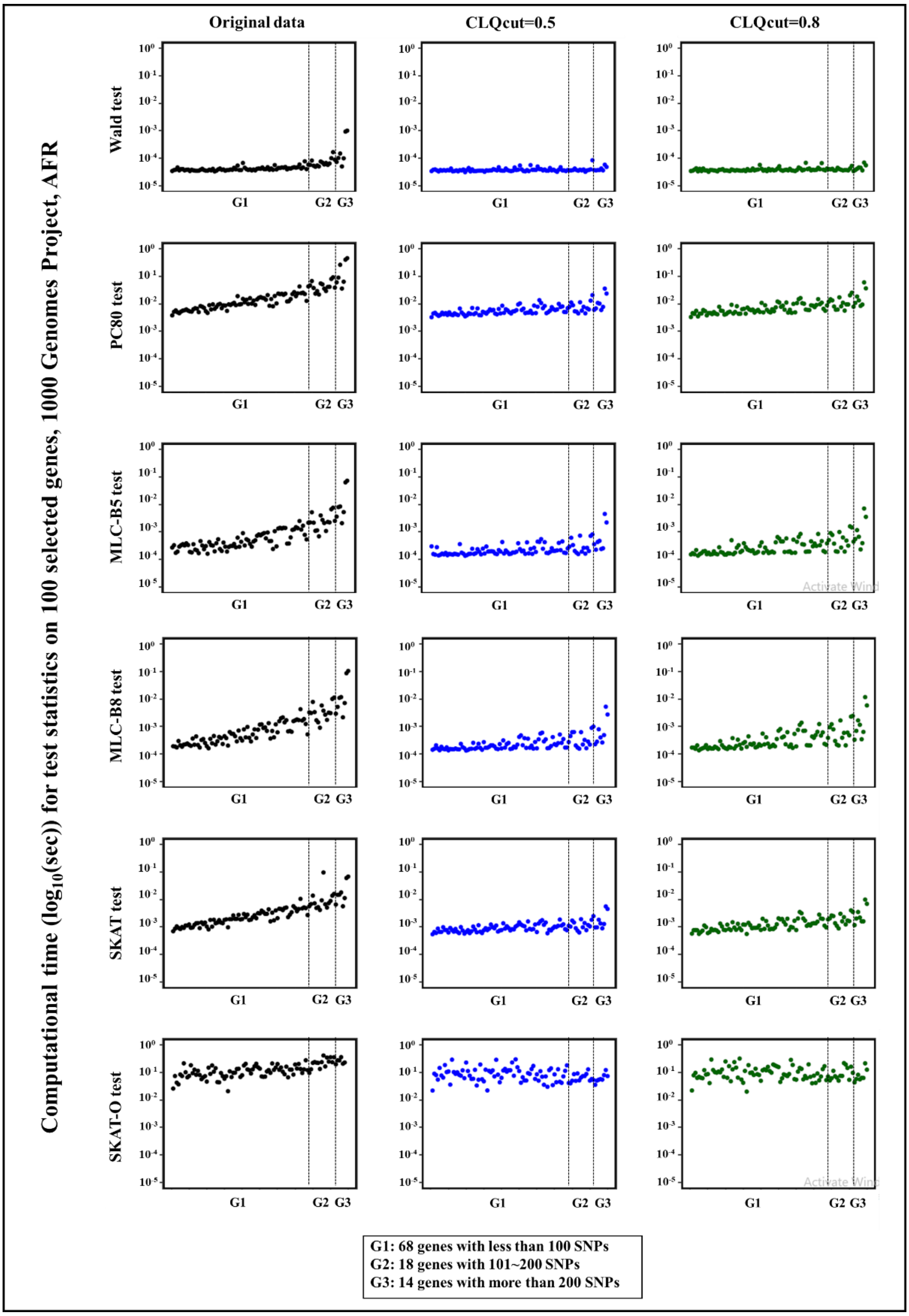

**FIGURE S22** The computational time for test statistics in a single replication of 1casual model on 100 selected genes, 68 genes with less than 101 SNPs, 18 genes with 101~200 SNPs, and 14 genes with more than 200 SNPs, using the original data and the DRLPC processed data with two threshold values for CLQcut (0.5, 0.8) and a threshold values 0.8 for PCcut, 1000 Genomes Project, AFR, chromosome 22. The x-axix represents the original gene size, sorted based on the number of SNPs.

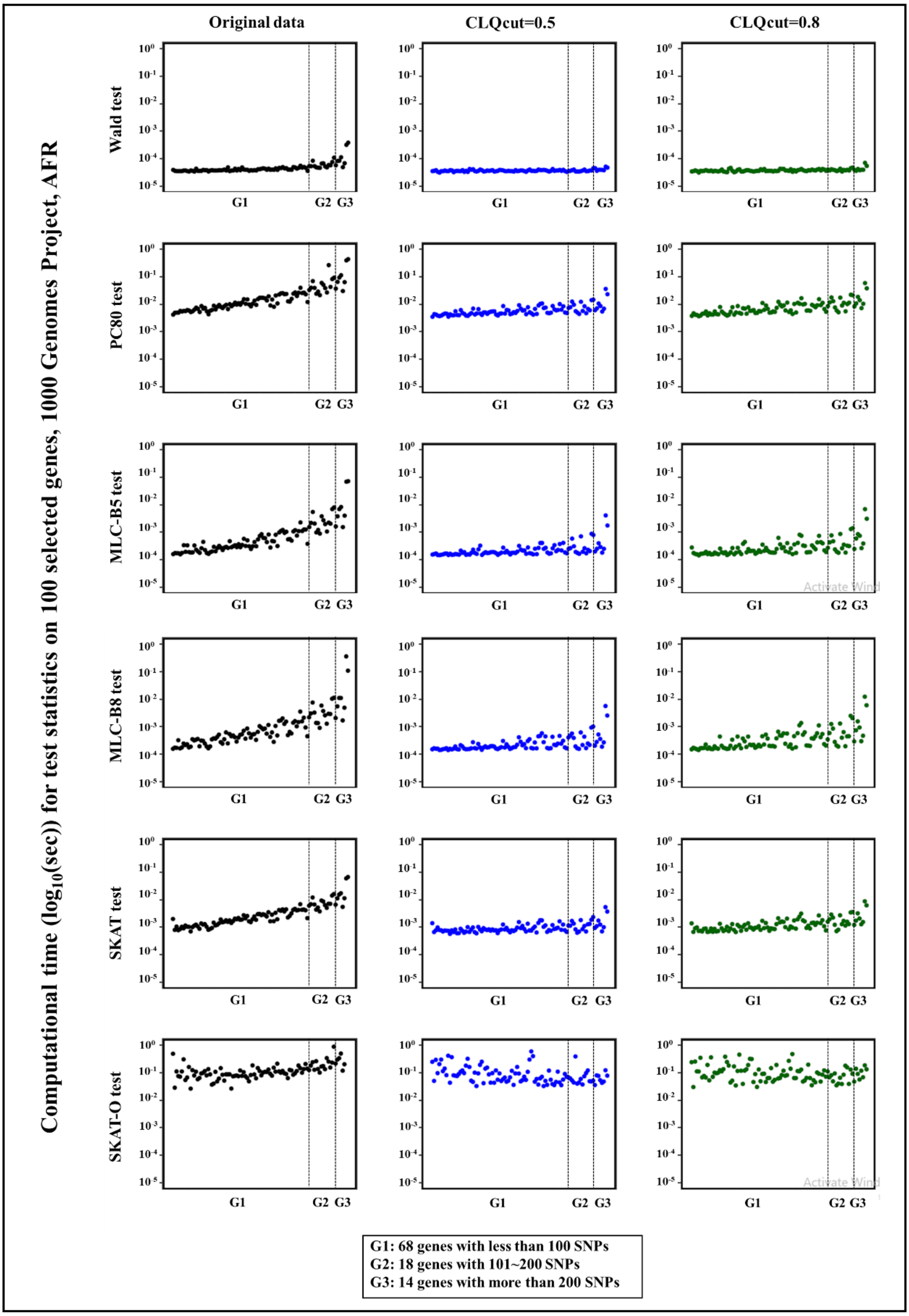

**FIGURE S23** The computational time for test statistics in a single replication of 2casual model on 100 selected genes, 68 genes with less than 101 SNPs, 18 genes with 101~200 SNPs, and 14 genes with more than 200 SNPs, using the original data and the DRLPC processed data with two threshold values for CLQcut (0.5, 0.8) and a threshold values 0.8 for PCcut, 1000 Genomes Project, AFR, chromosome 22. The x-axix represents the original gene size, sorted based on the number of SNPs.

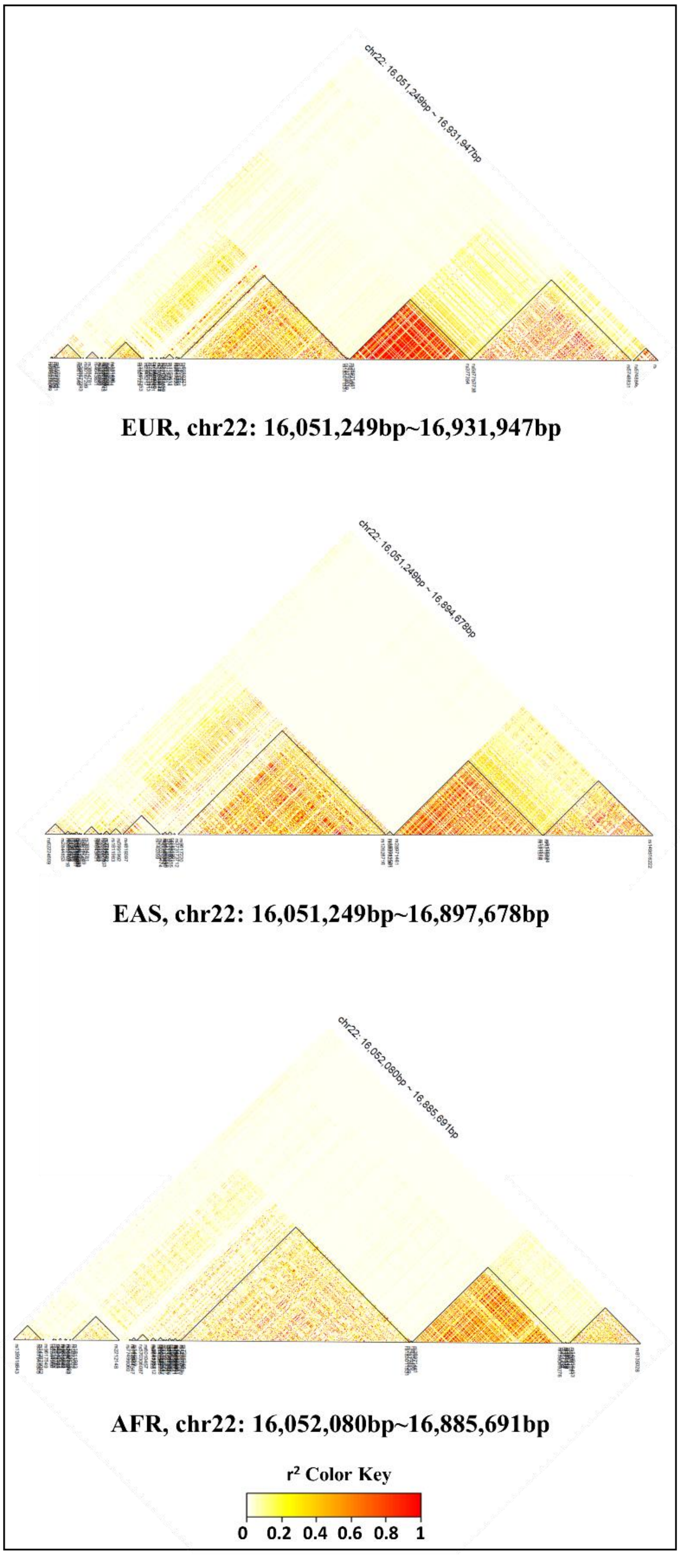

**FIGURE S24** LD($r^2$) heatmap of the first 1000 consecutive SNP on chromosome 22, using the LDheatmap function from the gpart package, across three super-populations, 1000 Genomes Project.
